# Different Roles of D1/D2 Medium Spiny Neurons in the Nucleus Accumbens in Pair Bond Formation of Male Mandarin Voles

**DOI:** 10.1101/2024.07.24.604911

**Authors:** Lizi Zhang, Yishan Qu, Larry J Young, Wenjuan Hou, Limin Liu, Jing Liu, Yuqian Wang, Lu Li, Xing Guo, Yin Li, Caihong Huang, Zijian Lv, Yitong Li, Rui Jia, Ting Lian, Hao Feng, Zhixiong He, Fadao Tai

**Affiliations:** Institute of Brain and Behavioural Sciences, College of Life Sciences, Shaanxi Normal University, Xi’an, Shaanxi, China; Department of Psychiatry and Behavioral Sciences, Silvio O. Conte Center for Oxytocin and Social Cognition, Center for Translational Social Neuroscience, Emory National Primate Research Center, Emory University, Atlanta, 30033, USA; Center for Social Neural Networks, University of Tsukuba, Tsukuba, 305-8555, Japan; Research Center for Prevention and Treatment of Respiratory Disease, School of Clinical Medicine, Xi’an Medical University, Xi’an, Shaanxi, China

**Author notes:** **Corresponding authors:** Fadao Tai and Zhixiong He;, 620 Xi Chang’ an Street, Chang’ an District, Xi’an city, Shaanxi province, China. These authors contributed equally to this work (Lizi Zhang, Yishan Qu).

**Keywords:** Dopamine, Nucleus accumbens, Pair bond, D1/D2 medium spiny neurons, Synaptic transmission

## Abstract

The mesolimbic dopamine (DA) system has been implicated in pair bond formation. However, involvements of DA release, real time activities, and electrophysiological activities of D1/D2 medium spiny neurons (MSNs) in the nucleus accumbens (NAc) shell in pair bonding remain unclear. This work verified that male mandarin voles after pair bonding released higher levels of DA in the NAc shell and displayed higher levels of D1 MSNs activity and lower levels of D2 MSNs activity upon sniffing their partners compared to upon sniffing an unknown female. Moreover, pair bonding induced differential alterations in both synaptic plasticity and neuronal intrinsic excitability in both D1 MSNs and D2 MSNs. In addition, chemogenetic inhibition of ventral pallidum (VP) -projecting D2 MSNs in the NAc shell enhanced pair bond formation, while chemogenetic activation of VP-projecting D2 MSNs in the NAc shell inhibited pair bond formation. These findings suggest that different neuronal activity of NAc shell D1 MSNs / D2 MSNs regulated by increasing DA release after pair bonding may be a neurobiological mechanism underlying pair bond formation.

## Introduction

In monogamous animals, pair bonding is a selective, lasting relationship between two adult individuals forming a breeding pair (Bales et al., 2021; López-Gutiérrez et al., 2021). Pair bonding is promoted by enduring social attraction between partners, and after sexual activity, partners remain together and jointly defend resources (Carter & Perkeybile, 2018; Numan & Young, 2016). In monogamous animals, pair bonding is usually characterized by partner preference (Carp et al., 2016), which is defined as the selective social preference for a pair mate rather than a stranger. In addition, both male and female prairie voles develop partner preferences following mating, and common neural pathways are engaged during pair bonding (Numan & Young, 2016). This relationship provides a secure psychological base for both partners and acts as a social buffer against life’s stresses in humans. Research showed that married people—especially in stable attachments—live longer than unmarried people (Jia & Lubetkin, 2020; Rendall, Weden, Favreault, & Waldron, 2011). In addition, the intimacy of couples affects not only their psychological status (for example by reducing instances of depression) and the rate of disease progression, but also their immune system and cardiovascular health (Ford & Young, 2021a; Frasure-Smith et al., 2000; Leserman et al., 1999; Leserman et al., 2000). Therefore, transformations of the neurobiological mechanisms in pair bonding are a powerful model for understanding social relationships in our own species, which carries implications for social attachments and romantic partnerships (Ford & Young, 2021b; Volkmar, 2001; K. A. Young, Liu, & Wang, 2008).

Most of the available knowledge regarding the neurobiology of pair bonding originates from studies on monogamous prairie voles (*Microtus ochrogaster*) (Borie et al., 2022; Gobrogge & Wang, 2015; K. A. Young, Gobrogge, Liu, & Wang, 2011). In prairie voles, mesolimbic dopamine (DA), which originates in the ventral tegmental area (VTA), plays an important role in pair bonding (Curtis & Wang, 2005). The nucleus accumbens (NAc) shows increased release of DA during sexual behavior (Aragona, Liu, Curtis, Stephan, & Wang, 2003; Gingrich, Liu, Cascio, Wang, & Insel, 2000), and this increased DA signaling is necessary for the formation of the pair bond. Non-selective DA receptor antagonists in the NAc inhibit mating-induced partner preferences (Aragona et al., 2003; Z. Wang et al., 1999). Because DA plays an important role in behaviors associated with reward, behavioral reinforcement, and addiction (Perra, Clements, Bernier, & Morikawa, 2011), we predict that the level of real-time DA release in voles may be different depending on whether they encounter their own partner or an unfamiliar vole of the opposite sex.

DA exerts its effect on the formation of the pair bond by binding with its receptors, dopamine type 1 receptor (D1R) and dopamine type 2 receptor (D2R). Moreover, DA affects pair bonding through binding with its receptor in the NAc shell, but not the core (Aragona et al., 2006b). Medium spiny neurons (MSNs) are the main type of efferent neuron in the NAc. NAc MSNs receive dopaminergic inputs to regulate the function of the NAc, which plays an important role in motivation, reward, and drug addiction (Cooper, Robison, & Mazei-Robison, 2017; Floresco, 2015; Russo & Nestler, 2013; Salgado & Kaplitt, 2015). MSNs are the main neurons in the NAc to receive, integrate, and transmit information, and can be divided into D1 MSNs and D2 MSNs according to different axon projections and distribution of different types of dopamine receptors (Gerfen & Surmeier, 2011; Surmeier, Ding, Day, Wang, & Shen, 2007; Yager, Garcia, Wunsch, & Ferguson, 2015). Pharmacological studies have shown that D1R and D2R play specific roles in the formation and maintenance of NAc-mediated pair bonding. Blocking D2R with its specific antagonist, while not blocking D1R, prevents pair bonding in prairie voles (Gingrich et al., 2000), suggesting the importance of D2R in the formation of the pair bond. In contrast, D1R is involved in the maintenance of the pair bond. D1R is upregulated in the NAc following pair bonding, and D1R signaling helps to maintain partner preference by inducing aggressive behavior toward strangers in paired male prairie voles (Aragona et al., 2006b). The excitability of MSNs is directly related to this change in social behavior. Willett et al. found that in prairie voles, MSNs exhibited notable differences in excitability compared with MSNs in rats; the amplitude of miniature excitatory postsynaptic current (mEPSC) and neuronal excitability decreased with increasing partner preference (Willett et al., 2018). A recent study found that cohabitation and mating in female prairie voles can alter both spontaneous and evoked synaptic activity of NAc MSNs; however, which type of MSNs is altered remains unknown (Borie et al., 2022). Although D1R and D2R have been found to play different roles in the formation and maintenance of pair bonding using pharmacological methods, whether D1/D2 MSNs display different levels of real time activity and show different electrophysiological properties after pair bond formation remains unclear.

The ventral pallidum (VP) receives projections from NAc shell D2 MSNs and D1 MSNs (Kupchik et al., 2015) and performs a major role in reward, motivation (Smith, Tindell, Aldridge, & Berridge, 2009), and partner preference formation (Lim et al., 2004). For example, microinjection of a psychostimulant into the VP is sufficient to form a conditioned place preference (Gong, Neill, & Justice, 1996). VP is also involved in both sexual behavior and social affiliation. In human males, VP activity increases during sexual arousal (Rauch et al., 1999). In addition, paired male Callicebus Titi monkeys *(Callicebus nigrifrons)* housed together with their partners show elevated VP glucose metabolism, suggesting that VP neural activity is related to pair bond maintenance (Bales, Mason, Catana, Cherry, & Mendoza, 2007). Additionally, inhibition of synaptic transmission from D2 MSNs to the VP can enhance motivation (Gallo et al., 2018). Activation of D1 MSNs to the VP mediates aversive, but not reward responses (Liu et al., 2022). However, the role of NAc shell D1 and D2 MSNs projecting to the VP in the formation of a pair bond still remains unclear. Therefore, it is important to examine whether NAc shell D2 and D1 MSNs projections to VP play different roles in the formation of the pair bond.

Mandarin voles (*Microtus mandarinus*) are socially monogamous rodents that are widely distributed across China (Li et al., 2021); they are an ideal animal model for examining the neurobiology of pair bonding. Previous results confirmed that male mandarin voles show significant preference to co-housed partners after 5–7 days of cohabitation (Li et al., 2021). Therefore, this study used male mandarin voles and examined the real time release of DA as well as the activity of two types of MSNs in the NAc shell upon sniffing both their partner and a stranger after pair bonding using *in vivo* fiber photometry. Alterations in the neural excitability, synaptic plasticity, and sensitivity to DA of NAc shell D1/D2 MSNs after pair bonding were assessed using whole cell patch-clamp recordings and pharmacological methods. Finally, a chemogenetic approach was used to activate or inhibit D2 MSNs and D1 MSNs projections to VP during pair bonding to disclose different roles of these two types of projections in pair bonding. This study advances the understanding of the neurobiological mechanisms underlying the formation of pair bonds and provides insights into the brain mechanism underlying enduring attachment between mating partners.

## Results

### DA release in the NAc shell was higher upon sniffing their partner than sniffing a stranger after pair bonding

After 7 days of cohabitation with females, male mandarin voles showed significant preference for their partners in a partner preference test, displaying reliable pair bonding (Figure S1). To explore alterations in neural activities induced by the formation of this pair bond, neural activities were measured before (at 3 days of cohabitation) and after (at 7 days of cohabitation) the formation of the pair bond. To determine the changes in DA release after pair bonding, extracellular DA was monitored by unilateral injection of rAAV-dLight 1.1, which encoded a fluorescent DA sensor into the NAc shell (Figure 1A–C). The sites of virus expression and optical fiber layout are shown in Figure 1B. The Control group was injected with AAV coding enhanced green fluorescent protein (EGFP). Histological method confirmed that both DA sensor and EGFP were expressed in the NAc shell (Figure 1B, Figure S2B).

**Figure 1.**
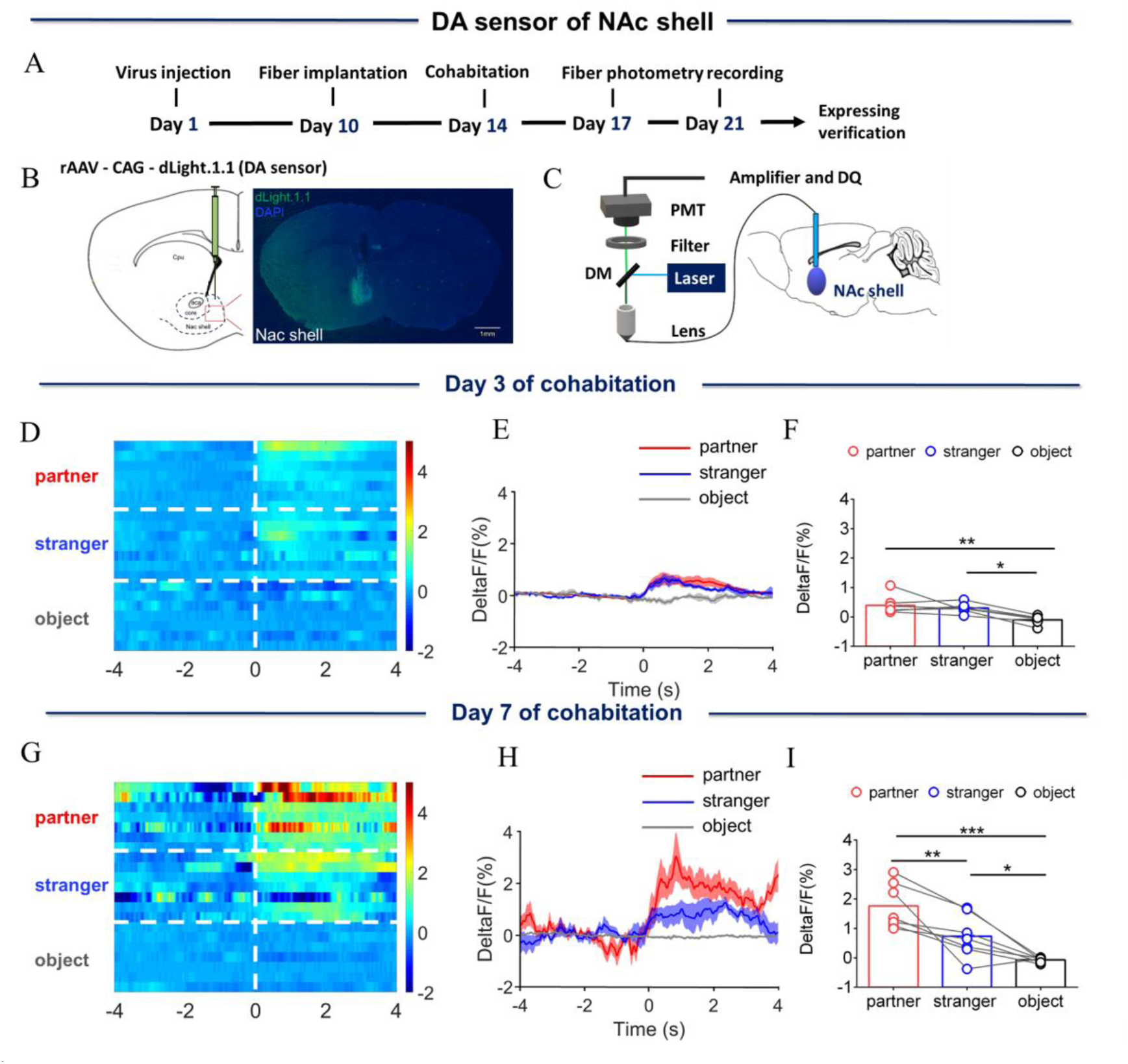
Dynamics of extracellular dopamine (DA) concentration within the nucleus accumbens (NAc) shell upon sniffing their partner or an unknown female. (A) Timeline of experiments. (B) Schematic diagrams depicting virus injection and recording sites and histology showing the expression of DA sensor within the NAc shell. Scale bar: 1 mm. (C) Schematic of the procedure used to record extracellular DA concentration in the NAc shell using fiber photometry. (D) Heat map illustrating the extracellular DA concentrations (ΔF/F, %) of the NAc shell when sniffing their partner, an unknown female, and an unrelated object. (E) Mean fluorescence signal changes of DA sensor during sniffing their partner (red line), an unknown female (blue line), or an object (gray line) after 3 days of cohabitation. The shaded areas along the differently colored lines represent the margin of error. (F) Quantification (One-Way Repeated Measures ANOVA) of changes in DA signals during sniffing of their partner, an unknown female, and an object after 3 days of cohabitation. (G) Heat map illustrating the extracellular DA concentration (ΔF/F, %) of the NAc shell when sniffing their partner, an unknown female, and an object. (H) Mean fluorescence signal changes of the DA sensor when sniffing their partner (red line), an unknown female (blue line), or an object (gray line) after 7 days of cohabitation. (I) Quantification (One-Way Repeated Measures ANOVA) of changes in extracellular DA concentration when sniffing their partner, an unknown female, and an object after 7 days of cohabitation. Error bars = SEM * represents *p* < 0.05, ** represents *p* < 0.01, and *** represents *p* < 0.001. See Supplementary Table 1 for detailed statistics.

Next, the dynamics of endogenous DA was measured in male mandarin voles when they sniffed their partner or a female stranger. Fluorescence signals were found to have increased in the NAc shell relative to baseline after cohabitation for 3 days when male voles sniffed either their partner or an unknown female (Figure S4A) (Paired t test: partner, t (6) = 3.290, p = 0.0166; stranger, t (6) = 4.884, p = 0.0028; object, t (6) = 1.745, p = 0.1315). Signal intensities showed no difference between sniffing their partner or an unknown female after three days cohabitation (Figure 1D–F) (One-Way Repeated Measures ANOVA: F (2.000, 12.00) = 8.702, p = 0.0001; partner vs. object, p = 0.0060; stranger vs. object, p = 0.0239). In addition, the fluorescence signals of the DA release in the NAc shell did not show differences between sniffing partners or strangers after 1 h or 3 days of cohabitation (Figure S3) (S3-F, One-Way Repeated Measures ANOVA: F (2.000, 12.00) = 19.59, p = 0.0002; partner vs. object, p = 0.0004, stranger vs. object, p = 0.0006. S2-I, One-Way Repeated Measures ANOVA: F (2.000, 12.00) = 10.85, p = 0.0087; partner vs. object, p = 0.0056, stranger vs. object, p = 0.0044. S3-J, Two-Way Repeated Measures ANOVA: group × treatment: F (2, 12) = 0.5558, p = 0.5877; group: F (2, 12) = 20.53, p = 0.0001; treatment: F (1, 6) = 0.1467, p = 0.7149).

However, after cohabitation for 7 days, although the extracellular DA concentration increased relative to baseline upon sniffing their partner or an unknown female (Figure S4B) (Paired t test: partner, t (6) = 6.020, p = 0.0009; stranger, t (6) = 2.611, p = 0.0401; object, t (6) = 2.209, p = 0.0692), the extracellular DA concentration was significantly higher upon sniffing their partner than a stranger (Figure 1G–I) (One-Way Repeated Measures ANOVA: F (2.000, 12.00) = 21.41, p = 0.0002; partner vs. stranger, p = 0.0097; partner vs. object, p < 0.0001, stranger vs. object, p = 0.0422). DA release did not change upon sniffing an object. In addition, changes in DA release upon sniffing same-sex cagemates and unknown males after cohabitation for 3 and 7 days were also measured. DA release in the NAc shell increased relative to baseline after cohabitation for 3 and 7 days when male voles sniffed an unknown male vole (Figure S5A, B) (A, Paired t test: partner, t (5) = 1.464, p = 0.2030; stranger, t (5) = 3.327, p = 0.0209; object, t (5) = 1.249, p = 0.2669. B, Paired t test: partner, t (5) = 1.998, p = 0.1021; stranger, t (5) = 2.602, p = 0.0481; object, t (5) = 0.1254, p = 0.9051), but no difference was found in signal intensities between sniffing same-sex cagemates and unknown males (Figure S6D–I) (One-Way Repeated Measures ANOVA: F (2.000, 10.00) = 4.864, p = 0.0335; stranger vs. object, p = 0.0396). Carrots are voles’ daily food. We found that when voles eat carrots, DA release increases in the NAc shell. The fluorescence signal of DA concentration was examined when voles ate carrot after cohabitation for 3 and 7 days, to rule out that the difference in fluorescence signals was caused by the difference in virus expression at different time points. The results showed that the fluorescence signal in the NAc shell increased upon eating carrot (Figure S7B–D) (Two-Way Repeated Measures ANOVA: group × treatment: F (1, 7) = 0.0274, p = 0.8732; group: F (1, 7) = 0.0557, p = 0.8201; treatment: F (1, 7) = 32.79, p = 0.0007), but no significant difference in fluorescence signal was found while eating carrot between voles that had cohabitated for 3 and 7 days. In control animals expressing EGFP in the NAc shell, changes in fluorescence intensities showed no difference between male voles sniffing their partner and an unknown female (Figure S2) (E, One-Way Repeated Measures ANOVA: F (2.000, 8.00) = 4.092, p = 0.0597. G, One-Way Repeated Measures ANOVA: F (2.000, 8.00) = 0.6107, p = 0.5665. I, One-Way Repeated Measures ANOVA: F (2.000, 8.00) = 0.0800, p = 0.9238. K, One-Way Repeated Measures ANOVA: F (2.000, 8.00) = 2.786, p = 0.1207). Furthermore, no significant differences were found between the fluorescence signals of DA release upon side-by-side contact with their partner and a stranger after 3 days of cohabitation (Figure S8A) (Two-tailed unpaired t-test: t (6) = 0.08005, p = 0.9388). However, the fluorescence signals of DA release upon side-by-side contact with their partner were higher than that during the same behavior with a stranger after 7 days of cohabitation (Figure S8B) (Two-tailed unpaired t-test: t (9) = 2.733, p = 0.0231).

### D2 MSNs activity in the NAc shell decreased while D1 MSNs activity increased upon sniffing their partner after pair bonding

The above results provide evidence that DA released within the NAc shell may play an important role in the formation of the pair bond among mandarin voles. Whether MSNs display different activities *in vivo* after the formation of partner preference was examined next. Fourteen days prior to cohabitation, rAAV-D1/D2-GCaMP6m, a D1/D2 genetically encoded fluorescent calcium sensor, was injected into the NAc shell. Dynamics of real-time calcium signals in the NAc shell in freely moving voles were recorded via a fiber photometry system in pair bonded or non-bonded mandarin voles (Figure 2A, B, Figure 3A, B). Immunofluorescence images showed that 84.42% of D1-GCaMP6m cells and 85.96% of D2-GCaMP6m cells were D1R-mRNA and D2R-mRNA positive (Figure 2C, Figure 3C).

**Figure 2.**
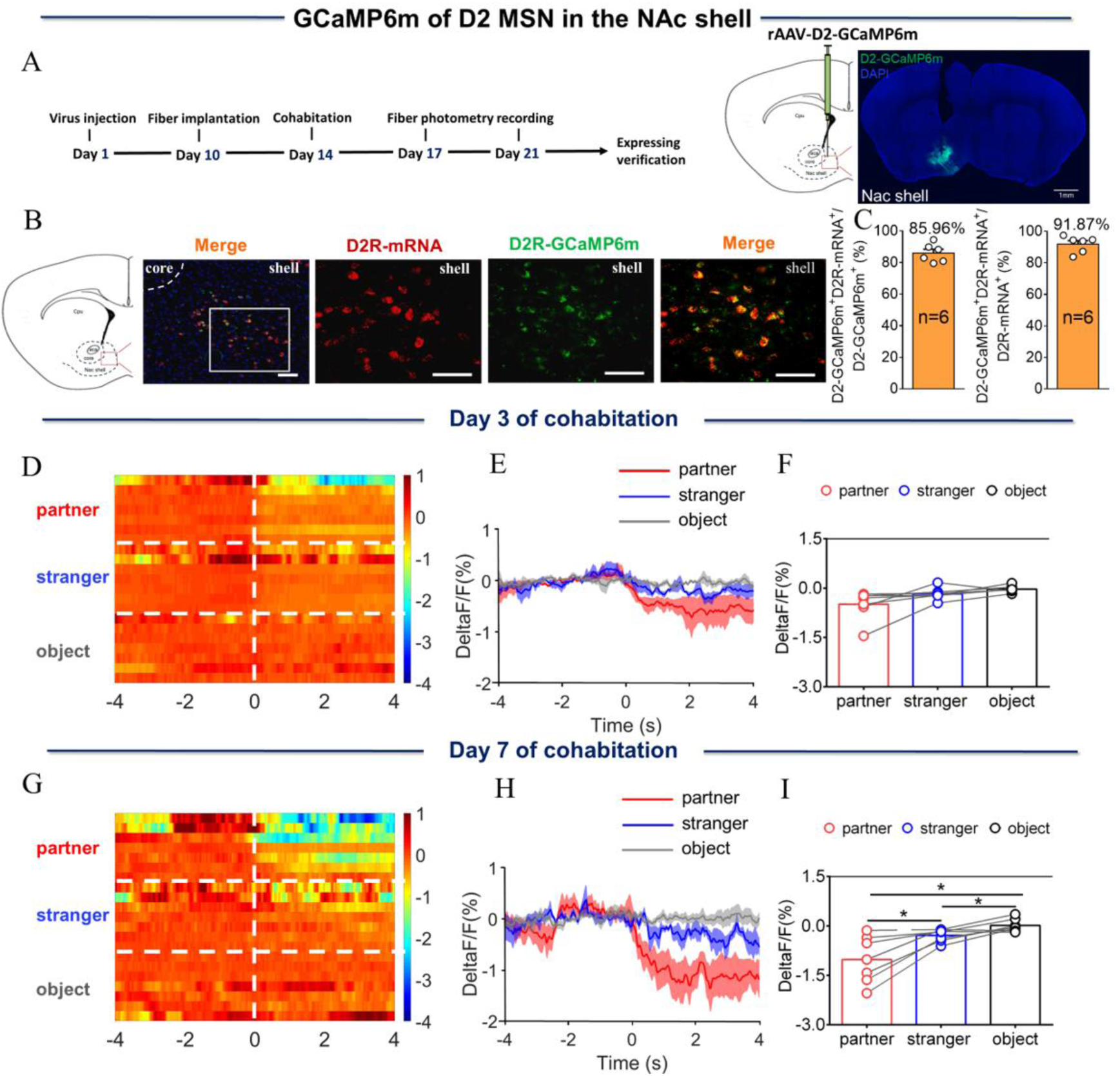
NAc shell D2 medium spinous neurons (MSNs) showing decreased activity upon sniffing their partner or an unknown female after cohabitation. (A) Left: Timeline of experiments; right: Schematic diagrams depicting virus injection and recording sites as well as histology showing the expression of D2-GCaMP6m within the NAc shell. Scale bar: 1 mm. (B) Overlaps of D2-GCaMP6m (green), D2R-mRNA (red), and DAPI (blue) in the NAc shell. Scale bar: 100 μm. (C) Statistical chart showing that D2-GCaMP6m was relatively restricted to D2R-mRNA positive neurons (n = 6 voles). (D) Heat map illustrating the calcium response (ΔF/F, %) of the NAc shell when sniffing a partner, an unknown female, or an object after cohabitation for 3 days. (E, H) Mean fluorescence signal changes of the calcium response during sniffing their partner (red line), an unknown female (blue line), or an object (gray line) after cohabitation for 3 days (E) and 7 days (H). The shaded areas along the different colors of lines show the margins of error. (F, I) Quantification (One-Way Repeated Measures ANOVA) of changes in calcium signals when sniffing their partner, an unknown female, or an object after cohabitation for 3 days (F) (n = 7 voles) and 7 days (I) (n = 7 voles). (G) Heat map illustrating calcium signals (ΔF/F, %) of the NAc shell when sniffing their partner, an unknown female, or an object after cohabitation for 7 days. All error bars = SEM. * represents *p* < 0.05. See Supplementary Table 1 for detailed statistics.

**Figure 3.**
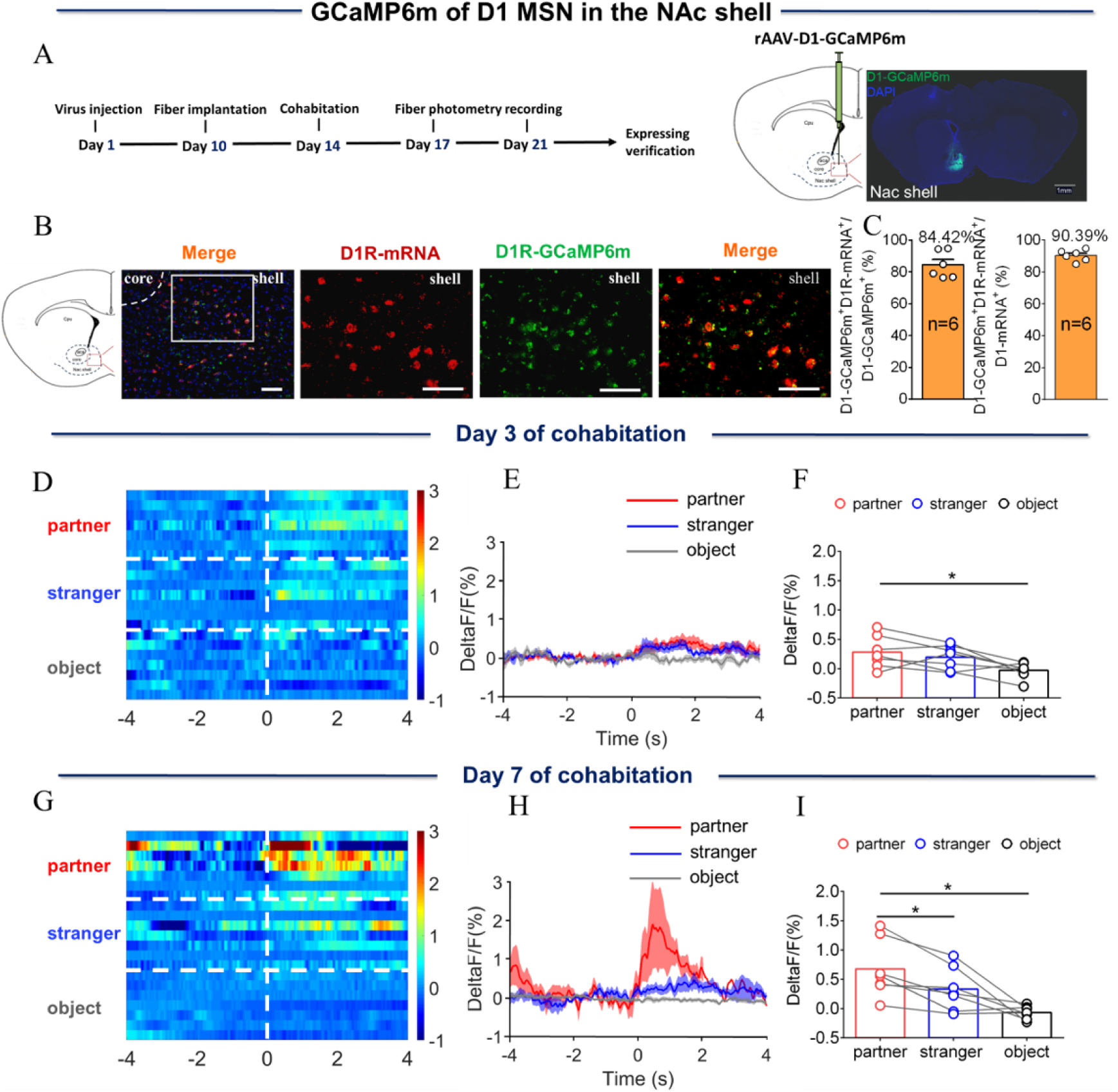
NAc shell D1 MSNs showing increased activity when sniffing their partner or an unknown female after cohabitation. (A) Left: Timeline of experiments; right: Schematic diagrams depicting the virus injection and recording sites as well as histology showing the expression of D1-GCaMP6m within the NAc shell. Scale bar: 1 mm. (B) Overlap of D1-GCaMP6m (green), D1R-mRNA (red), and DAPI (blue) in the NAc shell. Scale bar: 100 μm. (C) Statistical chart showing that D1-GCaMP6m was relatively restricted to D1R-mRNA positive neurons (n = 6 voles). (D) Heat map illustrating the calcium signals (ΔF/F, %) of the NAc shell when sniffing their partner, an unknown female, or an object after cohabitation for 3 days. (E, H) Mean fluorescence changes of calcium signals when sniffing their partner (red line), an unknown female (blue line), or an object (gray line) after cohabitation for 3 days (E) and 7 days (H). The shaded area along the different colored lines represents error margins. (F, I) Quantification (One-Way Repeated Measures ANOVA) of changes in calcium signals when sniffing their partner, an unknown female, or an object after cohabitation for 3 days (F) (n = 7 voles) and 7 days (I) (n = 7 voles). (G) Heat map illustrating the calcium signals (ΔF/F, %) of NAc shell when sniffing their partner, an unknown female, or an object after cohabitation for 7 days. All error bars = SEM. * represents *p* < 0.05. See Supplementary Table 1 for detailed statistics.

The results showed that the fluorescence signal of D1/D2 MSNs in the NAc shell was significantly increased relative to baseline when eating carrot after cohabitation for 3 and 7 days; voles who had experienced cohabitation for 3 and 7 days displayed no significant difference in fluorescence signals upon eating carrot (Figure S7E–J) (Two-Way Repeated Measures ANOVA: group × treatment: F (1, 7) = 1.1413, p = 0.2733; group: F (1, 7) = 1.428, p = 0.2711; treatment: F (1, 7) = 8.499, p = 0.0225). D2 MSNs activity was significantly reduced relative to baseline when sniffing either their partner or an unknown female after 3 and 7 days of cohabitation (Figure S4C, D) (C, Paired t test: partner, t (6) = 2.936, p = 0.0261; stranger, t (6) = 2.499, p = 0.0466; object, t(6) = 0.8186, p = 0.4443. D, Paired t test: partner, t (6) = 3.792, p = 0.0091; stranger, t (6) = 3.935, p = 0.0077; object, t (6) = 0.1841, p = 0.8600). In contrast, after 7 days of cohabitation, the activity when sniffing their partner was significantly lower than when sniffing a stranger (Figure 2G–I) (One-Way Repeated Measures ANOVA: F (1.212, 7.274) = 15.89, p = 0.0039; partner vs. stranger, p = 0.0366; partner vs. object, p = 0.0140, stranger vs. object, p = 0.0434). After 3 days of cohabitation, D2 MSNs activity significantly decreased relative to baseline when sniffing an unknown male (Figure S5C) (Paired t test: partner, t (5) = 2.479, p = 0.00559; stranger, t (5) = 2.654, p = 0.0452; object, t (5) = 0.5164, p = 0.6276). After 7 days of cohabitation, D2 MSNs activity significantly decreased relative to baseline when sniffing a same sex cagemate or an unknown male (Figure S5D) (Paired t test: partner, t (5) = 3.267, p = 0.0223; stranger, t (5) =2.789, p = 0.0385; object, t (5) = 1.035, p = 0.3479). However, no significant difference in fluorescence signals was found when sniffing a same sex cagemate or an unknown male (Figure S9G–I) (One-Way Repeated Measures ANOVA: F (2.000, 10.00) = 4.940, p = 0.0322; stranger vs. object, p = 0.0426). However, D1 MSNs activity was significantly increased relative to baseline when sniffing their partner or an unknown female after 3 days of cohabitation (Figure S4E, F) (E, Paired t test: partner, t (6) = 2.700, p = 0.0356; stranger, t (6) = 2.492, p = 0.0470; object, t (6) = 0.5216, p = 0.6207. F, Paired t test: partner, t (6) = 3.591, p = 0.0115; stranger, t (6) = 2.381, p = 0.0547; object, t (6) = 1.590, p = 0.1630). After 7 days of cohabitation, D1 MSNs activity significantly increased relative to baseline when sniffing their partner and was significantly higher when sniffing their partner than when sniffing an unknown female (Figure 3G–I) (One-Way Repeated Measures ANOVA:). After cohabitation for 3 or 7 days, D1 MSNs activity significantly increased relative to baseline when sniffing an unknown male (Figure S5E, F) (E, Paired t test: partner, t (5) = 1.788, p = 0.1338; stranger, t (5) = 2.613, p = 0.0475; object, t (5) = 0.4349, p = 0.6818. F, Paired t test: partner, t (5) = 2.009, p = 0.1008; stranger, t (5) = 2.983, p = 0.0307; object, t (5) = 1.049, p = 0.3420). No significant difference in fluorescence signals was found upon sniffing a same sex cagemate or an unknown male (Figure S10G–I) (One-Way Repeated Measures ANOVA: F (2.000, 10.00) = 5.533, p = 0.0241; stranger vs. object, p = 0.0278). D2 and D1 MSNs activity did not change upon sniffing an object. In addition, no significant changes were detected in the fluorescence signal upon sniffing their partner or a stranger in D1/D2 MSNs of voles injected with control virus (EGFP) without GCaMP6m sequence in the construct (Figures S11 and S12) (S11-E, One-Way Repeated Measures ANOVA: F (2.000, 8.00) = 0.7086, p = 0.5208. S11-G, One-Way Repeated Measures ANOVA: F (2.000, 8.00) = 1.450, p = 0.2901. S11-I, One-Way Repeated Measures ANOVA: F (2.000, 8.00) = 0.7471, p = 0.5041. S11-K, One-Way Repeated Measures ANOVA: F (2.000, 8.00) = 1.344, p = 0.3140. S12-E, One-Way Repeated Measures ANOVA: F (2.000, 8.00) = 4.162, p = 0.0577. S12-G, One-Way Repeated Measures ANOVA: F (2.000, 8.00) = 0.6909, p = 0.5287. S12-I, One-Way Repeated Measures ANOVA: F (2.000, 8.00) = 0.5571, p = 0.5936. S12-K, One-Way Repeated Measures ANOVA: F (2.000, 8.00) = 2.178, p = 0.1757). Similarly, the fluorescence signals of D2 MSNs and D1 MSNs upon side-by-side contact with their partner or an unknown female after 3 days of cohabitation did not differ. However, the fluorescence signals of D2 MSNs and D1 MSNs upon side-by-side contact with their partner were higher than that upon the same behavior toward an unknown female after 7 days of cohabitation (Figure S8C–F) (S8-C: Two-tailed paired t-test, t (6) = 0.6936, p = 0.5139; S8-D: Two-tailed unpaired t-test, t (10) = 2.813, p = 0.0184; S8-E: Two-tailed unpaired t-test, t (6) = 1.943, p = 0.1000; S8-F: Two-tailed unpaired t-test, t (7) = 2.392, p = 0.0481).

### Cohabitation with a partner alters the electrophysiological properties and synaptic transmission of D2/D1 MSNs in the NAc shell

DA binding with its target receptor (D2R) in the NAc is necessary for the formation of a pair bond. D2 MSNs were identified by infection of the NAc shell with rAAV-D2-mCherry virus and changes in synaptic transmission in D2 MSNs were recorded using whole-cell patch-clamp (Figure 4A). The frequency and amplitude of D2 MSNs spontaneous excitatory postsynaptic current (sEPSC) were found to be significantly higher in paired males than in naive males (Figure 4B, C) (Two-tailed unpaired t-test: frequency: t (20) = -3.634, p = 0.002; amplitude: t (20) = -2.619, p = 0.016). In addition, the frequency, but not the amplitude, of D2 MSNs spontaneous inhibitory postsynaptic currents (sIPSCs) was significantly increased in paired males compared to naive males (Figure 4B, D) (Two-tailed unpaired t-test: frequency: t (26) = -2.096, p = 0.046; amplitude: t (26) = -0.528, p = 0.602). Synapse-driven homeostatic plasticity of intrinsic excitability—long-term changes in synaptic activity that modify intrinsic excitability—has been found in the NAc shell. However, this study failed to detect a significant change in intrinsic excitability in D2 MSNs in both naive males and paired males (Figure 4E–H) (4F: Two-Way Repeated Measures ANOVA, group × treatment: F (8, 198) = 0.070, p = 0.999; group: F (1, 198) = 0.0003, p = 0.986; treatment: F (8, 198) = 16.767, p < 0.0001; 4G, Two-Way Repeated Measures ANOVA, group × treatment: F (8, 198) = 0.103, p = 0.999; group: F (1, 198) = 0.078, p = 0.780; treatment: F (8, 198) = 10.095, p < 0.0001; 4H, Two-tailed unpaired t-test, t (22) = 0.100, p = 0.921). To further examine the changes in excitatory and inhibitory synaptic transmission of D2 MSNs, evoked excitatory and inhibitory postsynaptic currents (PSCs) were recorded from the same cells. An increased E/I ratio of PSCs was found in D2 MSNs of paired males (Figure 4J) (Two-tailed unpaired t-test: t (10) = -3.499, p = 0.012). Finally, whether the sEPSC of D2 MSNs could be evoked by bath-applied DA (5 µM) in naive and paired males was also tested. The results showed a significant decrease in both the frequency and amplitude of D2 sEPSC in naive (Figure 4K, L) (Two-Way Repeated Measures ANOVA: frequencies: group × treatment: F (1, 14) = 0.0177, p = 0.896; group: F (1, 14) = 8.105, p = 0.0129; treatment: F (1, 14) = 7.189, p = 0.0179, Control_ Baseline vs Control DA, p = 0.0050, Cohabitation _ Baseline vs Cohabitation DA, p = 0.2296, Control_ Baseline vs Cohabitation Baseline, p = 0.0277, Control _DA vs Cohabitation DA, p = 0.0376; amplitude: group × treatment: F (1, 14) = 0.7937, p = 0.4042; group: F (1, 14) = 5.911, p = 0.0291; treatment: F (1, 14) = 14.78, p = 0.0018, Control_ Baseline vs Control DA, p = 0.007, Cohabitation Baseline vs Cohabitation _DA, p = 0.098, Control Baseline vs Cohabitation Baseline, p = 0.0472, Control DA vs Cohabitation DA, p = 0.0241), but not in paired mandarin voles (Figure 4L).

**Figure 4.**
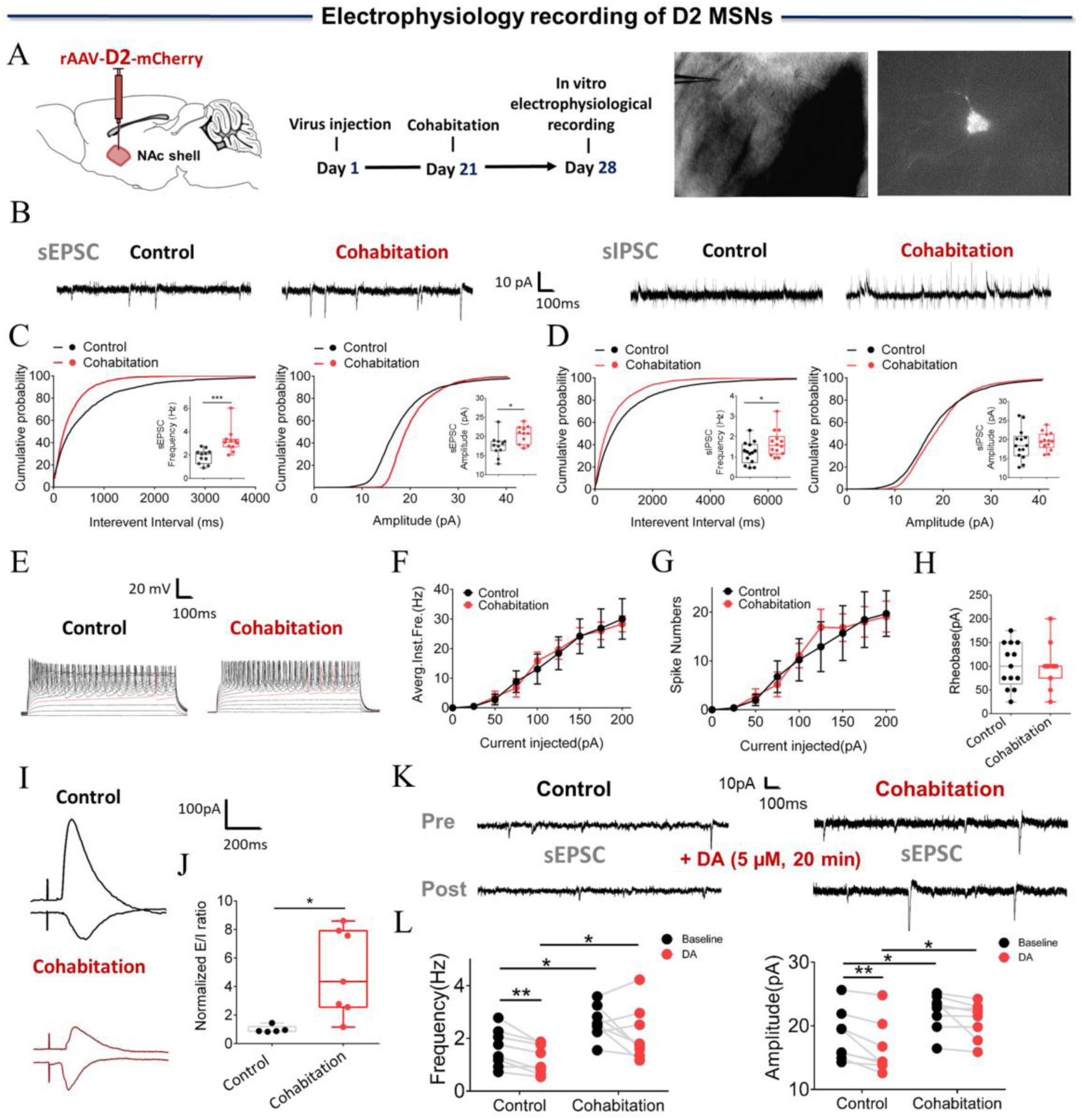
Synaptic transmission and neuronal excitability of D2 MSNs in the NAc shell changes after cohabitation. (A) Timeline of experiments (left), schematic diagrams depicting virus injection and recording sites (middle), and an D2 positive neuron with a micropipette (right). (B) Representative spontaneous excitatory postsynaptic current (sEPSC) and spontaneous inhibitory postsynaptic currents (sIPSC) traces from paired and naive voles. (C) NAc shell D2 MSNs in paired voles exhibited sEPSCs with higher frequencies (Cohabitation: n = 11 cells from four voles; Control: n = 11 cells from four voles) and peak amplitudes (Cohabitation: n = 11 cells from four voles; Control: n = 11 cells from four voles) than those observed in naive voles. (D) NAc shell D2 MSNs in paired voles exhibited sIPSCs with higher frequencies (Cohabitation: n = 14 cells from four voles; Control: n = 14 cells from four voles) than those observed in naive voles. (Amplitude: Cohabitation: n = 14 cells from four voles; Control: n = 14 cells from four voles) (E–H) The neuronal excitability of D2 MSNs in the NAc shell of paired voles was similar to that of naive voles. (Cohabitation: n = 11 cells from four voles; Control: n = 13 cells from six voles). (I and J) The excitation-inhibition ratio was higher in paired voles than in naive voles. (Cohabitation: n = 7 cells from four voles; Control: n = 5 cells from four voles). (K and L) D2-sEPSCs are evoked over a background of bath-applied DA in naive voles and cohabitated (7 days) voles. (Cohabitation: n = 8 cells from four voles; Control: n = 8 cells from three voles). Error bars = SEM. * represents *p* < 0.05, ** represents *p* < 0.01, and *** represents *p* < 0.001. See Supplementary Table 1 for detailed statistics.

Pharmacological assessment showed that D1R is involved in the maintenance of the pair bond; therefore, whole-cell patch-clamp recording was performed to determine whether D1 MSNs synaptic transmission in the NAc shell was altered by cohabitation. D1 MSNs showed reduced frequency and amplitude of sEPSC after cohabitation (Figure 5B, C) (Two-tailed unpaired t-test: frequency: t (33) = 2.816, p = 0.008; amplitude: t (33) = 3.322, p = 0.002). In addition, in sIPSC in D1 MSNs, the frequency decreased drastically, but not the amplitude (Figure 5B, D) (Two-tailed unpaired t-test: frequency: t (24) = 2.324, p = 0.024; amplitude: t (24) = 0.100, p = 0.921). Next, the intrinsic excitability was examined between naive and paired males, and a significant increase in intrinsic excitability of D1 MSNs was found (Figure 5E–H) (5F: Two-Way Repeated Measures ANOVA, group × treatment: F (8, 126) = 0.778, p = 0.623; group: F (1, 126) = 7.757, p = 0.006; treatment: F (8, 126) = 29.810, p <0.0001; 5G, Two-Way Repeated Measures ANOVA, group × treatment: F (8, 126) = 0.845, p = 0.565; group: F (1, 126) = 5.440, p = 0.021; treatment: F (8, 126) = 15.757, p < 0.0001; 5H, Two-tailed unpaired t-test, t (14) = 2.744, p = 0.016); differences in the E/I ratio of D1 MSNs were not found (Figure 5J) (Two-tailed unpaired t-test: t (12) = -1.429, p = 0.179). Then, it was tested whether the sEPSC of D1 MSNs can be evoked by bath-applied DA in naive and paired males. The results showed a significant increase in the frequency of D1 sEPSC in naïve males (Figure 5K, L) (Two-Way Repeated Measures ANOVA: frequencies: group × treatment: F (1, 18) = 4.784, p = 0.0422; group: F (1, 18) = 9.954, p = 0.0055; treatment: F (1, 18) =5.164, p = 0.0356, Control_ Baseline vs Control DA, p = 0.006, Cohabitation _ Baseline vs Cohabitation DA, p = 0.953, Control Baseline vs Cohabitation _ Baseline, p = 0.015, Control DA vs Cohabitation DA, p = 0.006; amplitude: group ×treatment: F (1, 18) = 0.007, p = 0.9343; group: F (1, 18) = 5.842, p = 0.0265; treatment: F (1, 18) = 0.6498, p = 0.4307, Control_ Baseline vs Control DA, p = 0.5285, Cohabitation Baseline vs Cohabitation DA, p = 0.63468, Control Baseline vs Cohabitation Baseline, p = 0.0282, Control DA vs Cohabitation DA, p = 0.029), but not in paired males (Figure 5L).

**Figure 5.**
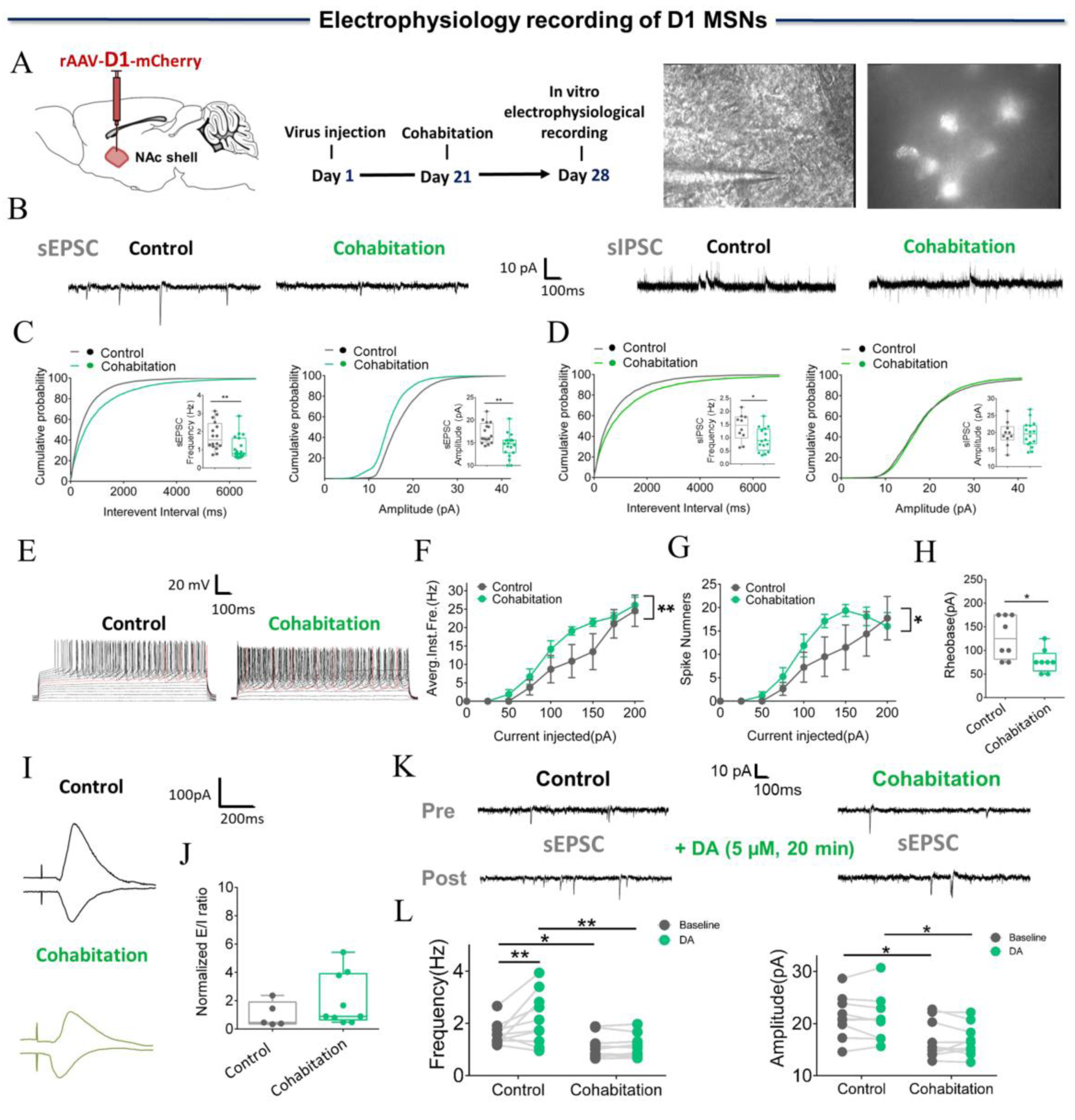
Synaptic transmission and neuronal excitability of D1 MSNs in NAc shell changes after cohabitation. (A) Timeline of experiments (left), schematic diagrams depicting virus injection and recording sites (middle), and an D1 positive neuron with a micropipette (right). (B) Representative sEPSC and sIPSC traces from paired and naive voles. (C) NAc shell D1 MSNs in paired voles exhibited sEPSCs with lower frequencies (Cohabitation: n = 19 cells from five voles; Control: n = 16 cells from four voles) and peak amplitudes (Cohabitation: n = 19 cells from five voles; Control: n = 16 cells from four voles) than those observed in naive voles. (D) NAc shell D1 MSNs in paired voles exhibited sIPSCs with lower frequencies (Cohabitation: n = 16 cells from four voles; Control: n = 10 cells from three voles) than those observed in naive voles (amplitude: Cohabitation: n = 16 cells from four voles; Control: n = 10 cells from three voles). (E–H) Neuronal excitability of D1 MSNs in the NAc shell of paired voles was higher than those observed in naive voles (Cohabitation: n = 8 cells from three voles; Control: n = 8 cells from four voles). (I, J) Excitation-inhibition ratio of paired voles was similar to naive voles. (Cohabitation: n = 9 cells from four voles; Control: n = 5 cells from three voles). (K, L) D1-sEPSCs are evoked over a background of bath-applied DA in naive voles and cohabitated (7 days) voles. (Cohabitation: n = 10 cells from five voles; Control: n = 10 cells from four voles). Error bars = SEM. * represents *p* < 0.05, and ** represents *p* < 0.01. See Supplementary Table 1 for detailed statistics.

### Effects of chemogenetic activation or inhibition of D2/D1 MSNs in the NAc shell projecting to the VP upon the formation of partner preference

MSNs in the NAc shell project extensively to the VP. To activate or inhibit these projections using a chemogenetic approach, AAV-DIO-hM3Dq-mCherry or AAV-DIO-hM4Di-mCherry were injected into the NAc shell and rAAV (Retro)-D2-Cre was injected into the VP to selectively express ‘Gq-DREADD’ or ‘Gi-DREADD’ in NAc shell VP-projecting D2 MSNs (Figure 6A). Immunohistochemical staining showed that 90.11% of mCherry cells co-expressed D2R (Figures 6B, C, and S14). To determine whether the ligand clozapine N-oxide (CNO) could activate or inhibit VP-projecting D2 MSNs, whole cell patch-clamp recordings were performed. The results showed that addition of 10 μM CNO remarkably increased the number of action potentials in the Gq-DREADD-transfected neurons (Figures 6D and 7D). In contrast, CNO caused a significant decrease of membrane potentials in Gi-DREADD-transfected neurons (Figures 6D and 7D). These results signal the specificity and validation of this virus strategy.

**Figure 6.**
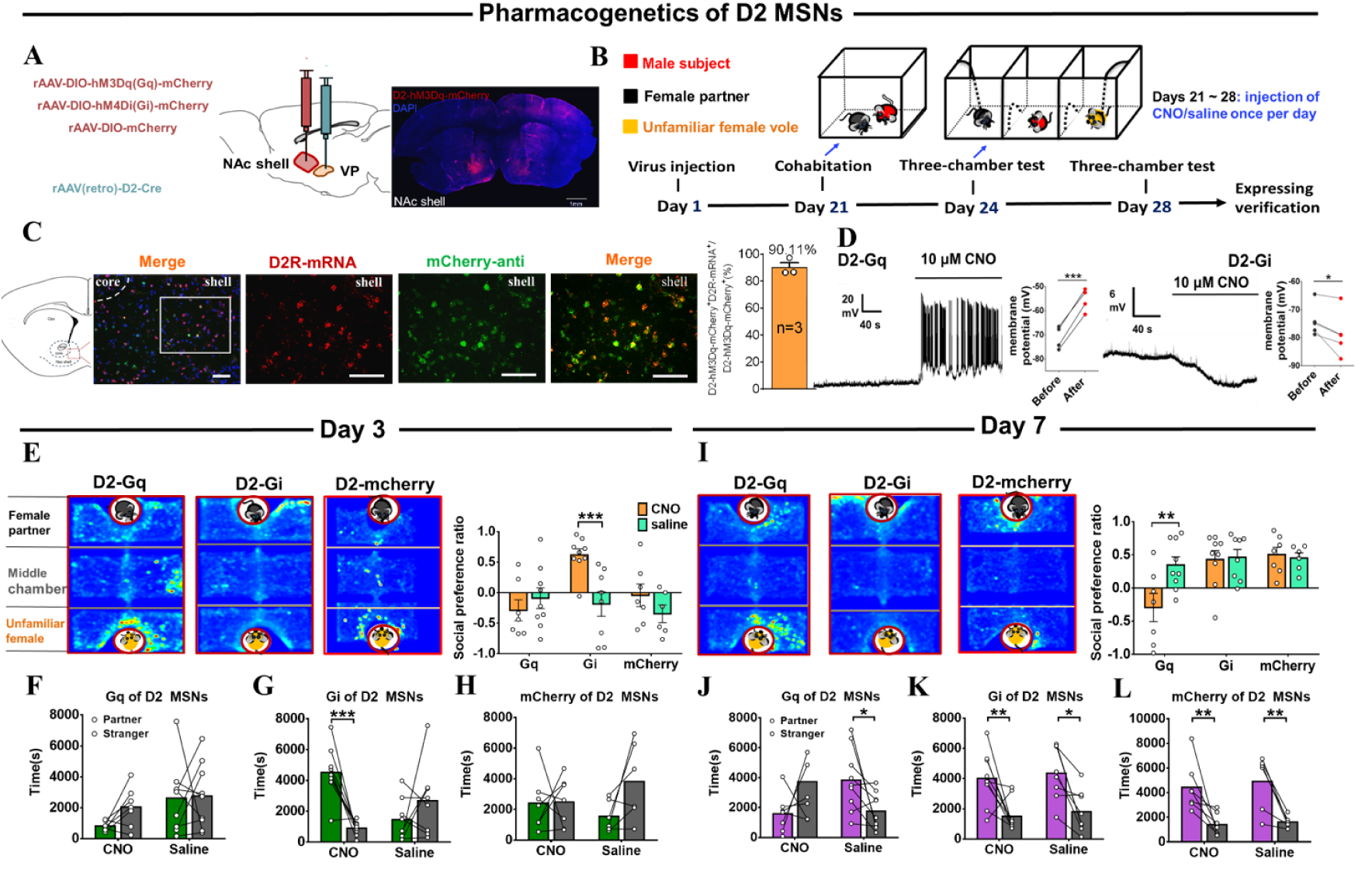
Effects of chemogenetic manipulation of NAc shell VP-projecting D2 MSNs on the formation of a partner preference. (A) Schematic of the chemogenetic viral strategy and injection sites as well as histology showing the expression of D2-hM3Dq-mcherry within the NAc shell. Scale bar: 1 mm. (B) Timeline of experiments. (C) Immunohistology image showing co-localization of hM3Dq-mCherry-anti expression (green), D2R-mRNA (red), and DAPI (blue) in the NAc shell. Scale bar: 100 μm. Statistical chart showing that D2-mRNA cells were relatively restricted to D2-hM3Dq-mCherry cells (n = 3 voles). (D) Representative traces from a Gq-DREADD (left) neuron and Gi-DREADD (right) neuron after CNO bath. (E) Representative heatmaps of the partner preference test after 3 days of cohabitation. Left: Gq group. Middle: Gi group. Right: mcherry group. (F–H) Quantification of side-by-side times in partner preference tests after cohabitation for 3 days. (I) Representative heatmaps of the partner preference test after cohabitation for 7 days. Left: Gq group. Middle: Gi group. Right: mcherry group. (J–L) Quantification of side-by-side times in the partner preference test after cohabitation for 7 days. (D2-hM3Dq: CNO: n = 7, saline: n = 9; D2-hM4Di: CNO: n = 9, saline: n = 8; D2-mCherry: CNO: n = 7, saline: n = 6). Error bars = SEM. * represents *p* < 0.05, ** represents *p* < 0.01, and *** represents *p* < 0.001. See Supplementary Table 1 for detailed statistics.

**Figure 7.**
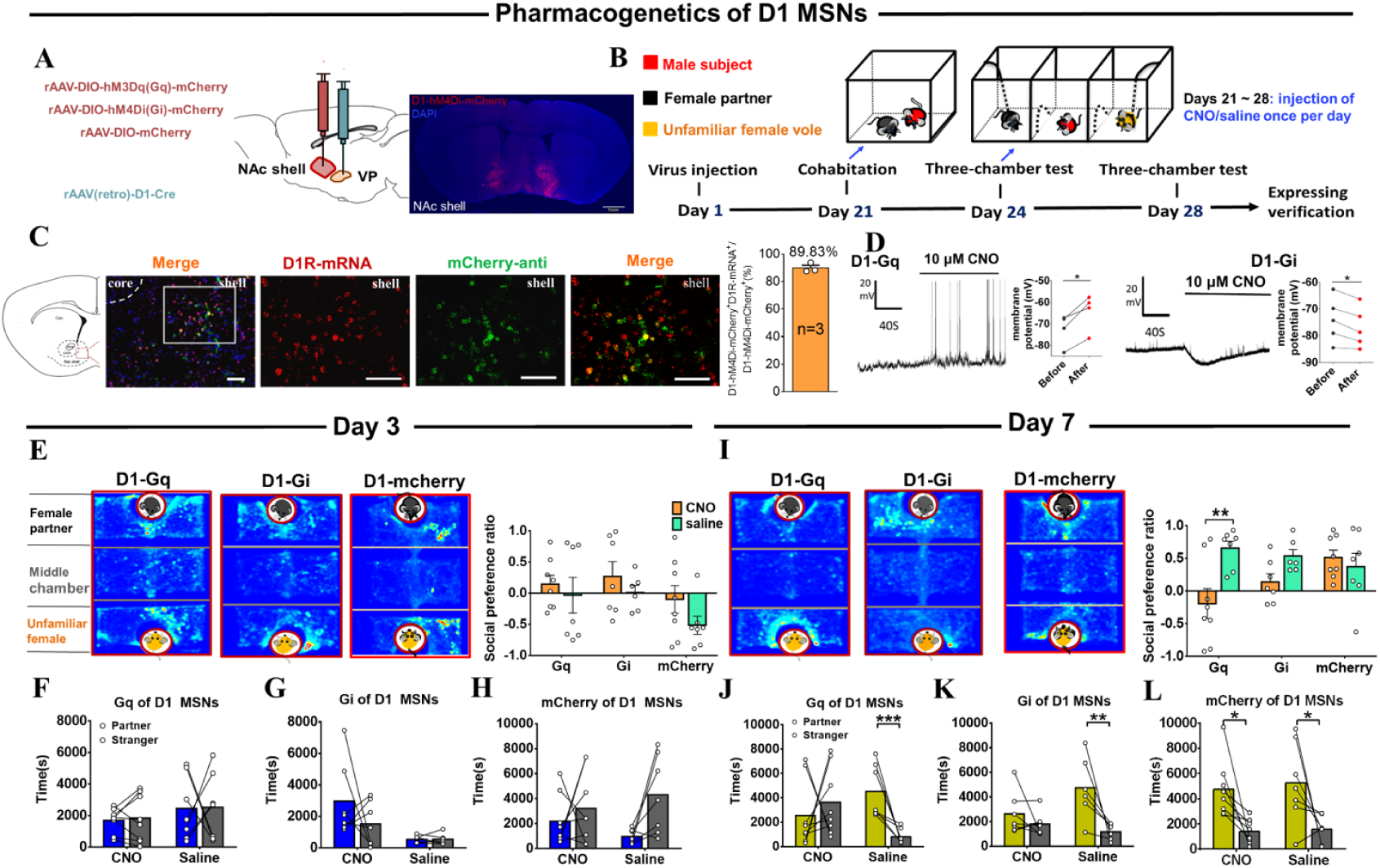
Effects of chemogenetic manipulation of NAc shell VP-projecting D1 MSNs on the formation of partner preference. (A) Schematic of chemogenetic viral strategy and injection sites as well as histology showing the expression of D1-hM4Di-mcherry within the NAc shell. Scale bar: 1 mm. (B) Timeline of experiments. (C) Immunohistology image showing co-localization of hM4Di-mCherry-anti expression (green), D1R-mRNA (red), and DAPI (blue) in the NAc shell. Scale bar: 100 μm. Statistical chart showing that D1-mRNA cells were relatively restricted to D1-hM4Di-mCherry cells (n = 3 voles). (D) Representative traces of a Gq-DREADD (left) neuron and Gi-DREADD (right) neuron after CNO bath. (E) Representative heatmaps of the partner preference test after 3 days of cohabitation. Left: Gq group. Middle: Gi group. Right: mcherry group. (F–H) Quantification of side-by-side times in the partner preference test after 3 days of cohabitation. (I) Representative heatmaps of the partner preference test after 7 days of cohabitation. Left: Gq group. Middle: Gi group. Right: mcherry group. (J–L) Quantification of side-by-side times in the partner preference test after 7 days of cohabitation. (D1-hM3Dq: CNO: n = 8, saline: n = 7; D1-hM4Di: CNO: n = 7, saline: n = 6; D1-mCherry: CNO: n = 8, saline: n = 7). Error bars = SEM. * represents *p* < 0.05, ** represents *p* < 0.01, and *** represents *p* < 0.001. See Supplementary Table 1 for detailed statistics.

During cohabitation, CNO (1 mg/kg) or saline was intraperitoneally injected daily (Figure 6A). The partner preference test was conducted on day 3 and 7 of cohabitation. Subsequent behavioral studies showed that CNO-treated mandarin voles with D2 MSNs Gi-DREADD virus infection showed increased side-by-side alignment with their partners compared with strangers in a partner preference test (Figure 6E and G) (6E, Two-way ANOVA: group × treatment: F (2, 40) = 4.942, p = 0.012; group: F (2, 40) = 4.265, p = 0.021; treatment: F (1, 40) = 4.967, p = 0.032; Gi_Saline vs Gi_CNO, p < 0.001; Gq_Saline vs Gq_CNO, p = 0.380; mCherry_Saline vs mCherry_CNO, p = 0.052. 6G, Kruskal-Wallis: H = 15.198, p = 0.002, CNO_Partner vs CNO_Stranger, p < 0.0001; saline_Partner vs saline_Stranger, p = 0.328); this shows that inhibition of VP-projecting D2 MSNs prompted the formation of partner preferences. Nevertheless, CNO-treated voles with VP-projecting D2 MSNs Gq-DREADD virus infection spent less time with their partners than with unknown females in the partner preference test; moreover, such activation of VP-projecting D2 MSNs also blocked partner preference after 7 days of cohabitation (Figure 6I, J) (6I, Two-way ANOVA: group × treatment: F (2, 40) = 3.798, p = 0.031; group: F (2, 40) = 6.797, p = 0.003; treatment: F (1, 40) = 3.375, p = 0.074; Gi_Saline vs Gi_CNO, p = 0.861; Gq_Saline vs Gq_CNO, p = 0.002; mCherry_Saline vs mCherry_CNO, p = 0.805. 6J, Kruskal-Wallis: H = 8.083, p = 0.044; CNO_Partner vs CNO_Stranger, p = 0.384; saline_Partner vs saline_Stranger, p = 0.033). In control virus subjects, CNO had no detectable effects on the behavioral performance in tests (Figure 6L) (Two-Way Repeated Measures ANOVA: group × treatment: F (1, 11) = 0.043, p = 0.839; group: F (1, 11) = 0.306, p = 0.591; treatment: F (1, 11) = 25.166, p = 0.0004; CNO_Partner vs CNO_Stranger, p = 0.009; saline_Partner vs saline_Stranger, p = 0.009).

Based on the results of chemogenetic manipulations described above, it was concluded that activation of D2 MSNs projecting to VP disrupted the formation of the pair bond while its inhibition improved the formation of partner preference in mandarin voles.

The chemogenetic approach was similarly used to test whether VP-projecting D1 MSNs in NAc shell regulate partner preference. AAV-DIO-hM3Dq-mCherry or AAV-DIO-hM4Di-mCherry and rAAV (Retro)-D1-Cre virus were bilaterally injected into the NAc shell and VP, respectively, to inhibit VP-projecting D1 MSNs (Figure 7A). Similarly, male mandarin voles received daily intraperitoneal injections of CNO (1 mg/kg) or saline during cohabitation, and partner preference was tested on days 3 and 7. In the partner preference test, CNO-treated voles with VP-projecting D1 MSNs Gi-DREADD virus infection showed trends to reduce side-by-side contact with their partners compared to saline-treated voles in the partner preference test after 7 days of cohabitation (Figure 7E, G, I, and K) (7E, Two-way ANOVA: group × treatment: F (2, 37) = 0.186, p = 0.831; group: F (2, 37) = 2.729, p = 0.078; treatment: F (1, 37) = 2.839, p = 0.100. 7G, Kruskal-Wallis: H = 9.597, p = 0.022; CNO_Partner vs CNO_Stranger, p = 0.087; saline_Partner vs saline_Stranger, p = 0.821. 7I, Two-way ANOVA: group × treatment: F (2, 37) = 4.934, p = 0.013; group: F (2, 37) = 0.873, p = 0.426; treatment: F (1, 37) = 7.633, p = 0.009; Gq_Saline vs Gq_CNO, p = 0.001; Gi_Saline vs Gi_CNO, p = 0.109; mCherry_Saline vs mCherry_CNO, p = 0.522. 7K, Two-Way Repeated Measures ANOVA: group × treatment: F (1, 11) = 4.281, p = 0.063; group: F (1, 11) = 1.513, p = 0.244; treatment: F (1, 11) = 11.027, p = 0.007; CNO_Partner vs CNO_Stranger, p = 0.377; saline_Partner vs saline_Stranger, p = 0.004). In addition, CNO-treated voles with VP-projecting D1 MSNs Gq-DREADD virus infection showed a significant reduction in side-by-side time spent with their partner than saline treated voles in the partner preference test after 7 days of cohabitation, and showed no partner preference (Figure 7E, F, I, and J) (7F, Kruskal-Wallis: H = 0.743, p = 0.863. 7J, Kruskal-Wallis: H = 12.819, p = 0.005; CNO_Partner vs CNO_Stranger, p = 0.256; saline_Partner vs saline_Stranger, p = 0.001). In control virus subjects, CNO had no significant effect on behavioral performance during testing (Figure 7L) (Two-Way Repeated Measures ANOVA: group × treatment: F (1, 13) = 0.033, p = 0.859; group: F (1, 13) = 0.325, p = 0.578; treatment: F (1, 13) = 15.559, p = 0.002; CNO_Partner vs CNO_Stranger, p = 0.033; saline_Partner vs saline_Stranger, p = 0.029). These results indicate that activation of the VP-projecting D1 MSNs impaired partner preference, but inhibition of these neurons did not produce significant effects on partner preference.

## Discussion

Although previous studies have demonstrated an association between the DA system and pair bond formation using pharmacological methods, the detailed neural mechanisms by which DA modulates partner preference remain elusive. By using AAV virus vectors and whole-cell patch-clamp recording, the present study found that, after 7 days of cohabitation, extracellular DA release in the NAc shell increased with social interaction; this increase was higher in response to a familiar partner than in response to an unfamiliar female. The observed increase in DA release resulted in alterations in activities of D2 and D1 neurons in the NAc shell selectively in response to their partner after extended cohabitation. Furthermore, the electrophysiological properties of D1 and D2 neurons in the NAc shell were also altered by cohabitation, displaying neural plasticity. Finally, opposite effects of DA on D1 and D2 neurons are supported by circuit manipulation, namely activation of D2 MSNs projections to VP or inhibiting/activating D1 MSNs projections to VP inhibits pair bonding, while inhibition of D2 MSNs projections to VP promotes pair bonding. These results reveal a neurobiological mechanism underlying the attachment to a partner and circuit mechanism underlying the formation of a pair bond.

To validate the choice of 3 days of cohabitation as an indicative timepoint of the DA response “before the pair bond”, an additional experiment was conducted to measure DA release in the NAc upon sniffing their partner and a stranger after 1 h and 3 days of cohabitation. The results affirmed that 1 h of cohabitation was not sufficient to form pair bonding. Fluorescence signals of the DA release in the NAc shell did not show differences upon sniffing their partner and a stranger after 1 h and 3 days of cohabitation (Figure S3). After 3 days of cohabitation, males showed similar levels of DA release upon sniffing their partner and a stranger than males after 1 h of cohabitation. Although according to partner preference tests, certain individuals preferred to stay with their partner after 3 days of cohabitation because of individual variation, the statistical result did not show significant preference for a certain partner. This result indicates that 3 days of cohabitation were not sufficient to form a stable pair bond at the behavioral and molecular levels.

In the present study, after 7 days of cohabitation, male mandarin voles showed more real-time DA release while they were sniffing their partner or had side-by-side contact with their partner than during the same behaviors directed toward an unknown female (Figure 1, S8A, B). Previous studies corroborated the important role of DA in the formation of a pair bond (Aragona et al., 2003; Z. X. Wang, Aragona, Liu, & Curtis, 2004). Moreover, local injections of the DA agonist apomorphine into the NAc shell enhanced partner preference formation; microinjection of the DA antagonist haloperidol into the NAc shell could prevent the formation of mating-induced partner preferences (Aragona et al., 2003). These results are also in line with previous reports, where mating was found to increase DA levels in the NAc among both male and female prairie voles (Aragona et al., 2003; Gingrich et al., 2000). Similarly, the results of the present study are also consistent with a recent study showing that upon sniffing and huddling with their partner, DA release was higher than during the same behaviors directed toward an unfamiliar vole (Pierce et al., 2024). In addition, the obtained results showed that extracellular DA levels upon sniffing their partner after 7 days of cohabitation were significantly higher compared to the extracellular DA after 3 days of cohabitation (Figure S13D). Furthermore, there was no significant difference in the fluorescence signals between same-sex strangers and cagemates after 3 or 7 days of cohabitation (Figures S6, S9, and S10). Changes in fluorescence signals were also not found during non-social behavioral bouts (Figure S5A–C). Combined with this experimental evidence, it is suggested that differences in the levels of DA release upon sniffing a partner and a stranger after 7 days of cohabitation may be due to the formation of a pair bond.

In addition, changes that were detected in the DA signal during side-by-side contact were much smaller than those detected upon sniffing (Figures 1 and S8). It is possible that the DA level was significantly elevated during the initial encounter, thus DA increased strongly during initial examination (Dai et al., 2022). One prominent feature of the examination-associated DA response is its fast adaptation, which diminishes exponentially later on (Dai et al., 2022). If sniffing is considered as an appetitive goal-directed behavior, side-by-side alignment may be a consummatory behavior. DA release generally precedes the onset of consummatory behaviors; as this behavior lasts, DA release may decrease significantly (Dai et al., 2022). In addition, DA is critical for the appetitive drive for social interaction, but not for low-effort, unconditioned consummatory behaviors (Pierce et al., 2024). Side-by-side contact is a long-lasting behavior that happens in a quiescent state. These may be the reasons why upon sniffing, the DA signal is stronger than that upon side-by-side alignment.

Furthermore, the present study found that D2 MSNs showed more enhanced suppression of activity after 7 days of cohabitation upon sniffing their partner compared with the activity after 3 days of cohabitation (Figure S13E). Similarly, D1 MSNs showed significant difference in response to sniffing their partner after 7 days of cohabitation compared to the response after 3 days of cohabitation (Figure S13F). These results are supported by pharmacological research, showing that activation of D2R in the NAc shell of female prairie voles accelerates partner preferences without mating, and the blockage of D2R antagonizes this behavior (L. J. Young & Wang, 2004). In addition, selective aggression toward a stranger was stopped by blocking D1R in the NAc (Resendez, Kuhnmuench, Krzywosinski, & Aragona, 2012). In the present study, increasing DA release in the NAc, decreasing calcium activity of D2 MSNs, and increasing calcium activity of D1 MSNs during social interaction with their partner after pair bond formation may be one mechanism underlying the preference for the partner after the formation of a pair bond.

In addition, electrophysiological properties and synaptic plasticity of MSNs expressing D1R and D2R after pair bonding were characterized for the first time. The NAc is assumed to play a role in emotion- and motivation-related learning and memory (Cardinal & Everitt, 2004; Kelley, 2004). DA in the NAc is assumed to promote pair bonding by facilitating the association between sensory cues of the partner and the reward of sexual behavior (Walum & Young, 2018). The sexual experience gained during cohabitation and pair bond formation may alter the electrophysiological properties of D1 and D2 MSNs. In a previous study, both D1R and D2R could specifically regulate the formation of partner preference (K. A. Young et al., 2011). Activation of D1R and D2R has different effects on the cyclic AMP (cAMP) signaling pathway, which has been proved to underly the specific regulation of pair bond formation by different types of DA receptors (Aragona & Wang, 2007). The physiological and direct cellular consequences of DA action on MSNs are the positive or negative effects of cAMP production, mediated by D1R or D2R, respectively (Lobo & Nestler, 2011). Although DR in the NAc shell is known to have direct connections with pair bond formation according to studies using animal models (Aragona et al., 2006b; L. J. Young & Wang, 2004), experimental evidence supporting a link between functional NAc shell MSNs and partner preference is still lacking. Although one recent study also found that pair bonding increases the amplitude of electrically induced EPSCs (Borie et al., 2022), the present study provides direct experimental evidence that the formation of partner preference induces different changes in both the neuronal activities and synaptic transmission of D1/D2 MSNs in male mandarin voles.

The present study found that the frequencies of sEPSC and sIPSCs were significantly enhanced after the formation of a pair bond in NAc shell D2 MSNs. The excitatory/inhibitory balance of D2 MSNs was enhanced after cohabitation. These results are not consistent with the findings from fiber photometry of calcium signals. One study showed that NAc D2 MSNs were linked to both ‘liking’ (food consumption) and ‘wanting’ (food approach) but with opposing actions; The high D2 MSNs activity signaled ‘wanting’, and the low D2 MSNs activity enhanced ‘liking’. D2 MSNs face a tradeoff between increasing ‘wanting’ by being more active or allowing ‘liking’ by remaining silent (Guillaumin, Viskaitis, Bracey, Burdakov, & Peleg-Raibstein, 2023). Therefore, the increase in frequencies of sEPSC and sIPSC in D2 MSNs may reflect two processes, liking and wanting differently and respectively. We suggest that hedonia and motivation might influence D2 MSNs activity during cohabitation and contribute to the processing of pair bond formation in a more dynamic and complex manner than previously expected. The difference may also be related to the interactions between neurotransmitters such as DA, glutamate, and gamma-aminobutyric acid (GABA) in the basal ganglia (Gerfen & Surmeier, 2011; Kalivas, Lalumiere, Knackstedt, & Shen, 2009; Meyer-Lindenberg, Domes, Kirsch, & Heinrichs, 2011; Pistillo, Clementi, Zoli, & Gotti, 2015; Shamay-Tsoory & Abu-Akel, 2016). DA modulates neurotransmitter release at both GABAergic and glutamatergic striatal synapses (Lovinger, 2010). It has been reported that the mEPSC amplitude is related to partner preference in prairie voles, but no difference in mEPSC frequency was detected between the NAc of voles and rats (Willett et al., 2018), which were not consistent with present results. The possible reasons for this variance are differences in brain regions (core and shell), and the types of MSNs were further distinguished in this research, which was neglected by previous studies. The frequency of mEPSC in overall MSNs may produce no changes, given that a decreased frequency of sEPSC in D1 MSNs and an increased frequency of sEPSC in D2 MSNs were observed. Further, the formation of a pair bond increased the D2 MSNs excitation/inhibition balance and amplified the D2 MSNs output in the present study, which would affect information processing in the NAc. Moreover, the frequencies of sEPSC and sIPSC were significantly reduced in the NAc shell D1 MSNs after pair bonding, whereas the intrinsic excitability increased after cohabitation with females. The bidirectional modifications (reduced synaptic inputs vs. increased excitability) observed in D1 MSNs might result from homeostatic regulation. The overall synaptic transmission may produce no net changes, given that reductions in both excitatory and inhibitory synaptic transmission of D1 MSNs were observed. Also, increases in the intrinsic excitability of D1 MSNs would result in an overall excitation gain on D1 MSNs. There are several explanations for the result that the same concentration of DA applied in a bath with slices from the cohabitation group did not change the frequency of D2/D1 sEPSC or the amplitude of sEPSCs. During cohabitation, the NAc shell DA concentration increased with mating behaviors (Aragona et al., 2003). The DA receptor has been activated, resulting in reduced DA receptor sensitivity to DA in the cohabitation group. In addition, DA exerts its role by binding to G protein-coupled receptors (i.e., D1R and D2R) that produce positive or negative effects via cAMP (Lobo & Nestler, 2011). In addition to DA, other neurotransmitters, such as glutamic acid, also produce effects via the same pathway. It remains unclear how these neurotransmitters interact in the MSNs to mediate activities. How other neural circuits (basolateral amygdala-NAc) jointly change the neuronal activity of MSNs after cohabitation has not yet been explored, which suggests a promising avenue for exploration in the future. Further studies of the distinct intracellular or synaptic mechanisms of both types of MSNs and their dynamics at various times during pair bond formation are necessary to better understand the cell type-specific functions and physiological consequences of pair bonding.

Moreover, the mesolimbic DA system regulates pair bond formation through the NAc-VP circuit (Numan & Young, 2016). The D1/D2 MSNs from the NAc shell equally innervate the VP through inhibitory projections (Kupchik et al., 2015). Combined with the findings in changes in DA release, calcium signals, and electrophysiological properties, it can be speculated that the activation of VP-projecting D2 MSNs caused by the DA release and the negative regulation loop of VP-projecting D1 MSNs in the NAc shell may result in higher excitability of VP and the promotion of pair bond formation. Further, the effects of DA on NAc seem to promote synaptic plasticity, enabling mating partners to continuously activate the NAc-VP attraction circuit, finally leading to lasting social attraction and bonding.

In the present study, DREADDs approaches were used to inhibit or excite NAc MSNs to VP projection and it was found that D1 and D2 NAc MSNs projecting to VP play different roles in the formation of a pair bond. Chemogenetic inhibition of VP-projecting D2 MSNs promoted partner preference formation, while activation of VP-projecting D2 MSNs inhibited it (Figure 6). Chemogenetic activation of D2 MSNs produced the opposite effect of DA on the D2 MSNs on partner preference, while inhibition of these neurons produced the same effect of DA on D2 MSNs. DA binding with D2R is coupled with Gi and produces an inhibitory effect (Lobo & Nestler, 2011). It is generally assumed that the activation of D2R produces aversive and negative reinforcement. These results were consistent with the reduced activity of D2 MSNs upon sniffing their partner in the fiber photometry test and the increased frequency and amplitude of sIPSC in the present study. Our results also align with other previous studies, which showed that chemogenetic inhibition of NAc D2 MSNs is sufficient to enhance reward-oriented motivation in motivational tasks (Carvalho Poyraz et al., 2016; Gallo et al., 2018). Inhibition of D2 MSNs during self-administration enhanced the response and motivation to obtain cocaine (Bock et al., 2013). This also suggests that the mechanism underlying attachment to a partner and drug addiction is similar.

Besides, in the present study, the formation of partner preference was inhibited following activation or inhibition of VP-projecting D1 MSNs, which is not consistent with conventional understanding of prairie vole behavior. Alternatively, DA binding with D1R is coupled with Gs and produces an excitatory effect (Lobo & Nestler, 2011), while the activation of D1R produces reward and positive reinforcement (Hikida, Kimura, Wada, Funabiki, & Nakanishi, 2010; Kwak & Jung, 2019; Tai, Lee, Benavidez, Bonci, & Wilbrecht, 2012). For example, activation of D1 MSNs enhances the cocaine-induced conditioned place preference (Lobo et al., 2010). In addition, D1R activation by DA promotes D1 MSNs activation, which in turn promotes reinforcement. However, a recent study found that NAc-ventral mesencephalon D1 MSNs promote reward and positive reinforcement learning; in contrast, NAc-VP D1 MSNs led to aversion and negative reinforcement learning (Liu et al., 2022). It is consistent with our results that activation of NAc-VP D1 MSNs pathway reduced time spent side-by-side and impaired partner preference after 7 days of cohabitation. In contrast to the inhibition of D2 MSNs, we found that inhibition of D1 MSNs did not elicit corresponding increases in partner preference. One possible explanation is that almost all D1 MSNs projecting to the VTA/substantia nigra (SN) send collaterals to the VP (Pardo-Garcia et al., 2019). For example, optogenetically stimulating VP axons may inadvertently cause effects in the VTA/SN through the antidromic activation of axon collaterals (Yizhar, Fenno, Davidson, Mogri, & Deisseroth, 2011). Therefore, chemogenetic inhibition of D1 MSNs may also inhibit DA neurons in the VTA, subsequently inhibiting the formation of a pair bond.

The dopamine and different types of dopamine receptors in the NAc may play different roles in regulation of pair bond formation and maintenance. The chemogenetic manipulation have revealed that VP-projecting D2 MSNs are necessary and more important in pair bond formation compared to VP-projecting D1 MSNs. It is consistent with previous pharmacological experiments that blocking of D2R with its specific antagonist, while D1R was not blocked, can prevent the formation of a pair bond in prairie voles (Gingrich et al., 2000). This indicates that D2R is crucial for the initial formation of the pair bond. D2R is involved in the reward aspects related to mating. In female prairie voles, D2R in the NAc is important for partner preference formation. The activation of D2R may help to condition the brain to assign a positive valence to the partner’s cues during mating, facilitating the development of a preference for a particular mate. In addition, the cohabitation caused the DA release, the high affinity Gi-coupled D2R was activated first, which inhibited D2 MSNs activity and promoted the pair bond formation. And then, after 7 days of cohabitation, the pair bonding was already established, the significantly increased release of dopamine significantly activated Gs-coupled D1R with the low affinity to dopamine, which increased D1 MSNs activity and maintained the formation of partner preference. While D1R is also present and involved in the overall process, its role in the initial formation of the pair bond is not as dominant as D2R (Aragona et al., 2006a). However, it still participates in the neurobiological processes related to pair bond formation. For example, in male mandarin voles, after 7 days of cohabitation with females, D1R activity in the NAc shell was affected during pair bond formation. The extracellular DA concentration was higher when sniffing their partner compared to a stranger, and this increase in DA release led to an increase in D1R activity in the NAc shell. In prairie voles, dopamine D1 receptors seem to be essential for pair bond maintenance. Neonatal treatment with D1 agonists can impair partner preference formation later in life, suggesting an organizational role for D1 in maintaining the bond (Aragona et al., 2006a). In pair-bonded male prairie voles, D1R is involved in inducing aggressive behavior toward strangers, which helps to maintain the pair bond by protecting it from potential rivals. In the NAc shell, D1 agonist decreases the latency to attack same-sex conspecifics, while D1 antagonism increases it (Aragona et al., 2006a). In summary, D2R is more crucial for pair bond formation, being involved in reward association and necessary for the initial development of the bond. D1R, on the other hand, is more important for pair bond maintenance, being involved in aggression and mate guarding behaviors and having an organizational role in maintaining the bond over time. We therefore suggest that D2 MSNs are more predominantly involved in the formation of a pair bond compared with D1 MSNs.

However, certain limitations of the present study should not be ignored. Firstly, in the present study, it was observed that about 20% of D1 promoter MSNs were labeled by D2-mRNA and about 14% D2 promoter MSNs were labeled by D1-mRNA vice (Figure S16). One reason for this difference may be that a small part of MSNs can co-express both D1R and D2R in the NAc shell (Valjent, Bertran-Gonzalez, Hervé, Fisone, & Girault, 2009). Another reason may be the non-specific expression of the virus. This means that a small group of MSNs were misclassified and miscounted. Although the effect of co-expressed neurons cannot be completely eliminated, the obtained results still reflect the changes in activity and physiological function of the vast majority of MSNs after the formation of a pair bond. In addition to DA, a variety of neurotransmitters or modulators are also involved in the formation of a pair bond, such as oxytocin, glutamate, and GABA. In the future, it is of great importance to study the synergistic regulation of pair bonding by multiple transmitters and systems in the NAc shell. Secondly, it is important to further examine whether other neural circuits also influence the formation of partner preference, such as basolateral amygdala projections to NAc. Moreover, in general, natal philopatry among mammals is female-biased in the wild(Brody & Armitage, 1985; Greenwood, 1983; Ims, 1990; Solomon & Jacquot, 2002). Social mammals are rarely characterized by exclusively male natal philopatry (Solomon & Jacquot, 2002). Males often disperse from natal area to a new place. Thus, male rodents may play a dominant role in the formation and maintenance of mating relationships. This is a reason why we investigate pair bonding in males firstly. Certainly, female mate selection, as well as sexual receptivity or refusal through olfactory cues from males, thereby affects the formation and maintenance of pair bonding (Hoglen & Manoli, 2022). This is also the reason why we should focus on the mechanisms underlying pair bonding formation in females in future research. Despite these limitations, the obtained results reveal the neurobiological mechanism underlying attachment to a partner as well as the circuit mechanism underlying the formation of a pair bond (Figure S17).

## Author Contributions

F.T. designed the research; L.Z. and Y.Q. performed the research; L.Z., Y.Q., W.H., L.L., Y.L., T.L., R.J. and H.F. performed the fiber photometry; L.Z. and Y.Q. performed *in vitro* electrophysiology recordings; L.Z., Z.L., J.L., Y.W., L.L., X.G., C.H., and Y.L. performed chemogenetic experiments; Y.Q. performed immunohistochemistry; L.Z. and Z.H. analyzed the data; L.J.Y., L.Z., F.T., and Z.H. wrote the paper.

L.Z. and Y.Q. contributed equally to this work.

## Acknowledgements

This work was supported by STI2030-Major Projects (2022ZD0205101), the National Natural Science Foundation of China (31970424 and 31901082), the Natural Science Foundation of Shaanxi Province, China (2020JQ-412), the China Postdoctoral Science Foundation (2019M653534), the Fundamental Research Funds for Central University (GK201903065 and GK202007008), the Department of Science and Technology of Shaanxi province (No. 2022PT-44), the Natural Science Basic Research Plan in Shaanxi Province of China (2023-JC-YB-207), and the Fundamental Research Funds for Central University of China (GK202301012). The contribution by LJY was also supported by an NIH grant.

## Methods

### Animals

Subjects used in the present study were the F2 generation of mandarin voles bred in the laboratory. The voles were weaned 21 days after birth and lived in same-sex colonies in different polycarbonate cages (44 × 22 × 16 cm) and were housed in a 22–24 °C, 12-h light/dark cycle with food and water *ad libitum*. The voles used in the experiment were ∼70–90 days old. All laboratory procedures were conducted in accordance with the Guidelines for the Care and Use of Laboratory Animals in China and the regulations of the Animal Care and Use Committee of Shaanxi Normal University.

### Key resources table-Virus

| Designation | Identifiers | Additional information | Source or reference |
| --- | --- | --- | --- |
| rAAV-CAG-dLight1.1-WPRE-hGH_polyA | PT-1138 | Serotype titer 2/9, 5.00 × 10<sup>12</sup> vg/ml | BrainVTA |
| rAAV-D1-GCaMp6m-WPRE-hGH_polyA | PT-2749 | Serotype titer 2/9, 2.55 × 10<sup>12</sup> vg/ml | BrainVTA |
| rAAV-D2-GCaMp6m-WPRE-hGH_polyA | PT-2750 | Serotype titer 2/9, 2.69 × 10<sup>12</sup> vg/ml | BrainVTA |
| rAAV-D1-EGFP-WPRE-bGH_polyA | PT-0214 | Serotype titer 2/9, 2.78 × 10<sup>12</sup> vg/ml | BrainVTA |
| rAAV-D2-EGFP-WPRE-hGH_polyA | PT-3245 | Serotype titer 2/9, 2.26 × 10<sup>12</sup> vg/ml | BrainVTA |
| rAAV-D1-mCherry-WPRE-bGH_polyA | PT-0757 | Serotype titer 2/9, 3.66 × 10<sup>12</sup> vg/ml | BrainVTA |
| rAAV-D2-mCherry-WPRE-bGH_polyA | PT-0367 | Serotype titer 2/9, 2.44 × 10<sup>12</sup> vg/ml | BrainVTA |
| rAAV-D1-CRE-WPRE-hGH_polyA | PT-1217 | Serotype titer 2/R, 4.72 × 10<sup>12</sup> vg/ml | BrainVTA |
| rAAV-D2-CRE-WPRE-bGH_polyA | PT-0571 | Serotype titer 2/R, 5.00 × 10<sup>12</sup> vg/ml | BrainVTA |
| rAAV-CAG-EGFP | PT-0305 | Serotype titer 2/9, 5.28 × 10<sup>12</sup> vg/ml | BrainVTA |
| rAAV-EF1α-DIO-hM3D(Gq)-mCherry | PT-0042 | Serotype titer 2/9, 2.70 × 10<sup>12</sup> vg/ml | BrainVTA |
| rAAV-EF1α-DIO-hM4D(Gi)-mCherry | PT-0043 | Serotype titer 2/9, 5.53 × 10<sup>12</sup> vg/ml | BrainVTA |
| rAAV-EF1α-DIO-mCherry | PT-0013 | Serotype titer 2/9, 5.19 × 10<sup>12</sup> vg/ml | BrainVTA |

### Stereotaxic surgery and virus infusions

Male mandarin voles were anesthetized with 1.5–3.0% isoflurane inhalant gas (R5-22-10, RWD, China) and placed in stereotaxic instruments (68045, RWD, China). The brain coordinates of virus injection were as follows: NAc shell: AP: +1.5, ML: ±0.99, DV: −4.2; VP: AP: +0.8, ML: ±1.2, DV: −4.9. Next, a 10 µL Hamilton microsyringe (7635-01, HAMILTON) was used to inject the virus using a microsyringe pump (KDS legato 130, RWD, China) at a rate of 80 nl/min. After the injection, the needle was left in place for 10 min, and was then slowly removed to prevent virus leakage. For fiber photometry recording, voles were mounted on a stereotaxic apparatus and a fiber optical cannula (diameter 2.5 mm, NA 0.37, depth 6 mm, RWD, China) was implanted into the site ∼0.15 mm above the NAc shell 10 days after virus injection. Only voles with the correct locations of virus expression and optical fibers were used for further analyses.

Firstly, real-time release of DA and activity of two types of MSNs in the NAc shell upon sniffing their partner, an unknown female, or an object on days 3 and 7 of cohabitation were measured using an *in vivo* fiber photometry system. Male mandarin voles were injected with rAAV-CAG-dLight1.1 (400 nl), rAAV-D1-GCaMP6m (400 nl), rAAV-D2-GCaMP6m (400 nl), rAAV-CAG-EGFP (300 nl), rAAV-D1-EGFP (400 nl), and rAAV-D2-EGFP (400 nl) into the NAc shell. The viruses used in the present study were obtained from brainVTA company and listed in Key resources table - Virus. The non-coding promoter sequence of the mouse D1R/D2R gene was predicted and amplified for validation by the company. The sequence was then constructed and packaged by the AAV virus vector by brainVTA company. The detailed sequence information can be obtained from brainVTA company.

Alterations in the neural excitability and synaptic plasticity of NAc shell D1/D2 MSNs after cohabitation were then examined using whole cell patch-clamp recordings. Animals were divided into sexually naive and cohabitation groups. Voles were injected with rAAV-D1-mCherry (400 nl) or rAAV-D2-mCherry (400 nl) into the NAc shell to identify D1/D2 MSNs. Twenty-one days after virus injection, the males were cohabitated with females for 7 days.

Finally, a chemogenetic approach was used to activate or inhibit D1/D2 MSNs projecting to the VP on days 3 and 7 of cohabitation to disclose different roles of these two types of projections in the formation of a pair bond. AAV-EF1α-DIO-hM3Dq(Gq)–mCherry (400 nl), AAV–EF1α–DIO– hM4Di(Gi)-mCherry (400 nl), or AAV-EF1α-DIO-mCherry (400 nl) were bilaterally injected into the NAc shell and rAAV (Retro)-D1-Cre (400 nl) and rAAV (Retro)-D2-Cre (400 nl) were injected into the VP. The D1/D2 promoter sequences originated from mice. Subjects were divided into Gq, Gi, and mCherry groups. After cohabitation, the partner preference test was conducted on days 3 and 7 days of cohabitation.

All female voles were ovariectomized and primed through subcutaneous administration of estradiol benzoate (17-β-Estradiol-3-Benzoate, Sigma, 2 μg dissolved in sesame oil starting 3 days prior to the experiments) (Borie et al., 2022).

### Fiber photometry

The fiber photometry system (ThinkerTech, Nanjing, China) was used as previously described (Li et al., 2021). Briefly, the emission light from modulated blue 480 LED (50 mW) was reflected with a dichroic mirror, and then delivered to the brain by an optical objective lens coupled with an optical commutator, which excited the D1/D2-GCaMP6m or the DA sensor located in the NAc shell. The excitation light was passed through another band pass filter, into a CMOS detector (Thorlabs, Inc; DCC3240M), and finally recorded by Labview software (TDMSViewer; ThinkerTech, Nanjing, China). Fourteen days after virus injection, male voles were cohabited with females. Previous results confirmed that male mandarin voles show significant preference to co-housed partners after 5–7 days of cohabitation (Feng et al., 2021); therefore, the real time release of DA and the activity of D1/D2 MSNs in the NAc shell were observed on days 3 and 7 days of cohabitation (partner: DA sensor: day 3, n = 7; day 7, n = 7. D2-GCaMP6m: day 3, n = 7; day 7, n = 7. D1-GCaMP6m: day 3, n = 7; day 7, n = 7; same sex cagemate: DA sensor: day 3, n = 6; day 7, n = 6. D2-GCaMP6m: day 3, n = 6; day 7, n = 6. D1-GCaMP6m: day 3, n = 6; day 7, n = 6).

After 3 and 7 days of cohabitation, subjects were anesthetized with isoflurane and connected to a multi-mode optic fiber patch cord (ThinkerTech, Nanjing, China; NA: 0.37, OD: 200 μm) connected to a fiber photometry apparatus. The cage for social interaction is an open field (44 × 22 × 16 cm) in which subjects were exposed to their partner or a stranger. Males were exposed to their partner or an unfamiliar female (each exposure lasted for 30 min) in random order in a clean social interaction cage. The changes in fluorescence signals during these social interactions with their partner, an unfamiliar vole of the opposite sex, or an object (Rubik’s Cube) were collected and digitalized by CamFiberPhotometry software (ThinkerTech). To rule out that the difference in fluorescence signals was caused by the difference in virus expression at different time points, we used the same experimental strategy in new male mandarin voles and measured the fluorescence signal changes upon eating carrot after 3 and 7 days of cohabitation (The male mandarin voles were fasted for four hours before the test.). Fiber photometry was performed to measure the release of DA and the activity of D1/D2 MSNs in the NAc shell during social interaction with a partner or a stranger, including sniffing and side-by-side contact (The side-by-side contact behavior is defined as significant physical contact with a social vole and huddling in a quiescent state.). In the same sex cagemate group, male mandarin voles were cohabited with a same-sex vole for 3 and 7 days. Males were then exposed to their same sex cagemate or an unfamiliar male (each exposure lasted for 30 min and was conducted in random order) in a clean social interaction cage. The changes in fluorescence signals upon sniffing the same sex cagemate, an unfamiliar male vole, or an object were collected and digitalized by CamFiberPhotometry software (ThinkerTech).

All data were analyzed in MatLab 2019a; the ΔF/F values represent the release of DA and activity of D1/D2 MSNs during sniffing and side-by-side contact. The baseline (F_0_) for all events and behaviors was the averaged fluorescence signal of -10 to 0 s prior to exposure to the stimulated voles. The fluorescence signals after the start of sniffing and side-by-side contact are recorded as F. These fluorescence signals during the entire 30 min session were recorded and analyzed. The fluorescence change (ΔF/F) values were calculated as (F – F_0_) / F_0_. Then, the time window was determined based on the duration of a behavioral bout. To estimate the calcium response, the average (ΔF/F) was calculated during the 4 s time window following the beginning of sniffing and side-by-side contact. Otherwise, the data points where the sniffing bout occurred within 4 s of a prior sniffing bout were excluded from data analysis. The sniffing and side-by-side contact that occurred in a fiber photometry experiment were examined using jwatcher, a behavior event scoring software (https://www.jwatcher.ucla.edu/, Dan Blumstein’s Lab & the late Christopher Evans’ lab, Sydney).

### Slice preparation and whole-cell patch clamp recording

Four weeks after virus injection, voles were anesthetized with 1.5–3.0% isoflurane inhalant gas and the brains were harvested into oxygenated artificial cerebrospinal fluid (ACSF) containing 125 mM sodium chloride (NaCl), 2.5 mM potassium chloride (KCl), 25 mM glucose, 25 mM sodium hydrogen carbonate (NaHCO_3_), 1.25 mM monosodium phosphate (NaH_2_PO_4_), 2 mM calcium chloride (CaCl_2_), and 1 mM magnesium chloride (MgCl_2_), gassed with 5% CO_2_ / 95%O_2_. Sagittal NAc slices (300 μm) were cut on a vibratome (VT 1200S, Leica, Germany) with oxygenated ACSF (32–34 °C), incubated at 32 °C for 30 min, followed by 1 h of incubation at room temperature before recording. Voltage- and current-clamp whole-cell recordings were performed using standard techniques at room temperature. Electrodes were pulled from 1.5 mm borosilicate-glass pipettes on a P-97 puller (Sutter Instruments).

Whole-cell patch-clamp recordings were obtained with a Multiclamp 700B amplifier, digitized at 10 kHz using a Digidata 1440 A acquisition system with Clampex 10.2, and analyzed with pClamp 10.5 software (Molecular Devices). Only cells that maintained a stable access resistance (<30 MΩ) throughout the entire recording were analyzed. mCherry MSNs in the NAc shell were identified under IR-DIC optics and fluorescence microscopy. sIPSCs (D2 MSNs: Cohabitation, n = 14 cells from four voles; Control, n = 14 cells from four voles) (D1 MSNs: Cohabitation, n = 16 cells from four voles; Control, n = 10 cells from three voles) were isolated by adding D-AP5 (50 µM, Sigma), and CNQX (20 µM, Sigma) in ACSF while recording at a holding potential of 0 mV. sEPSCs (D2 MSNs: Cohabitation, n = 11 cells from four voles; Control, n = 11 cells from four voles) (D1 MSNs, Cohabitation: n = 19 cells from five voles; Control, n = 16 cells from four voles) were isolated by adding PTX (100 µM, Sigma) in ACSF while recording at a holding potential of −70 mV. For evoked glutamatergic and GABAergic PSC recordings, a concentric electrode was used to stimulate axon terminal input to the NAc shell. Evoked EPSCs were isolated by voltage-clamping neurons at the reversal potential of the inhibition (−70 mV), whereas evoked IPSCs were recorded at the reversal potential of the excitation (0 mV). A total of 20 recording events with intervals of 30 s at each holding potential were used for analysis. The E-I ratio was calculated from baseline subtracted traces as the average EPSC amplitude divided by the average IPSC amplitude. All data were normalized to the mean of the control E-I ratio (D2 MSNs: Cohabitation, n = 7 cells from four voles; Control, n = 5 cells from three voles) (D1 MSNs: Cohabitation, n = 9 cells from four voles; Control, n = 5 cells from three voles). For the bath-applied DA test, sEPSCs were isolated by adding PTX (100 µM) in ACSF while recording at a holding potential of −70 mV. After 10 min of recording, slices were perfused with 5 μM DA (Sigma). The total recording time for each cell was 40 min and the last 10 min of the recording were assessed (D2 MSNs: Cohabitation, n = 8 cells from four voles; Control, n = 8 cells from three voles) (D1 MSNs: Cohabitation, n = 10 cells from five voles; Control: Control, n = 10 cells from four voles). Electrode resistance was ∼5–7 MΩ when filled with internal solution consisting of 130 mM CsMeSO_3_, 8 mM CsCl, 1 mM MgCl2, 0.3 mM EGTA, 10 mM HEPES, 4 mM ATP (magnesium salt), 0.3 mM GTP (sodium salt), and 10 mM phosphocreatine (pH 7.4, 300 mOsm).

To assess the neuronal action potential, MSNs were stimulated with step-current pulses (1000 ms, ranging from 0 to 250 pA) by adding PTX (100 µM, Sigma), CNQX (20 µM, Sigma), and D-AP5 (50 µM, Sigma). (D2 MSNs: Cohabitation, n = 11 cells from four voles; Control, n = 13 cells from six voles) (D1 MSNs: Cohabitation, n = 8 cells from three voles; Control, n = 8 cells from four voles). To determine the activation and inhibition effect of CNO (BrainVTA, CNO-02) in Gi or Gq virus infected neurons, spontaneous firing of action potentials in the cell was recorded in the current-clamp mode. After 3 min of baseline recording, NAc slices from mice with injection of Gq and Gi viruses were perfused with 10 μM CNO for 7 min. The total recording time for each cell was 10 min. Electrode resistance was ∼5–7 MΩ when filled with internal solution consisting of: 130 mM K-Gluconate, 5 mM NaCl, 10 mM HEPES, 0.5 mM EGTA, 2 mM Mg-ATP, and 0.3 mM Na-GTP (pH 7.3, 280 mOsm).

### Chemogenetics

Three weeks after virus injection, male mandarin voles were cohabited with females. For the D1-mCherry-hM3Dq/hM4Di group (hM3Dq: CNO, n = 8; saline, n = 7. hM4Di: CNO, n = 7; saline, n = 6) or the D2-mCherry-hM3Dq/hM4Di group (hM3Dq: CNO, n = 7; saline, n = 9. hM4Di: CNO, n = 9; saline, n = 8), male voles were injected with CNO (1 mg/kg, i.p. injection) or saline once per day during the 7-days cohabitation period. On days 3 and 7 of cohabitation, a partner preference test (3 h) was conducted 3 h after CNO injection. For the D1-mCherry group (CNO: n = 8; saline: n = 7) or the D2-mCherry group (CNO: n = 7; saline: n = 6), the same test procedure was used.

### Partner preference test

Partner preference formation is a reliable indicator of pair bonding and is characterized by selective contact and mating with partners rather than with strangers. The three-chamber partner preference test was first developed in Dr Sue Carter’s lab and has since been adopted by many other laboratories (K. A. Young et al., 2011). The apparatus consists of a three-chamber (60 × 40 × 20 cm) arena where the middle chamber is connected to two identical chambers. Partner and unfamiliar voles of the opposite sex were placed into the two chambers on opposite sites. Before testing, subjects were adapted to the test arena for 30 min, and partner and opposite sex voles were also habituated for 10 min. During the 3 h of partner preference test, two stimulus animals were firstly confined to their own chambers; then, the subjects were placed into the middle chamber and allowed to move freely. Real-time side-by-side contact behaviors were videotaped and quantified by smart 3.0. A longer duration of side-by-side contact spent with partners (but not unfamiliar opposite-sex female voles) indicates successful formation of partner preferences. The social preference ratio was calculated as follows: (time spent on partner side - time spent on stranger side) / (time spent on partner side + time spent on stranger side).

### Fluorescence *in situ* hybridization

Fluorescence *in situ* hybridization (FISH) was conducted using RNAscope® Multiplex Fluorescent Reagent Kit v2 (Advanced Cell Diagnostics 323100) following the manufacturer’s protocol. Voles were perfused transcardially with 0.1 M PBS (pH 7.4) followed by 4% paraformaldehyde in 0.1 M PBS. Then, brains were collected and postfixed for 24 h in 4% paraformaldehyde at 4 °C followed by 24 h in 20% sucrose and 24h in 30% sucrose, and were cut into 12 μm (6 series/brain) coronal sections using a cryostat (Leica Biosystems, Germany). Sections were mounted on slides and stored at -80 °C. Probes used in this study were RNAscope® Prob-Mo-Ppib (#533491), RNAscope® Negative Control Prob-DapB (#310043), RNAscope™ Probe-Mo-Drd1-C3 (#588161-C3), and RNAscope Probe-Mo-Drd2-C2 (#534471-C2). To verify specificity and effectiveness of the rAAV-GCaMP6m virus, MSNs (co-labeled by D1R/D2R mRNA and D1-GCaMP6m/D2-GCaMP6m virus) were quantified. 20× images including the NAc shell were acquired and boxed areas (300 × 300 μm^2^) were selected. Three representative sections were chosen per brain. The NAc ^D1/D2 mRNA (AF594)^ positive cells and co-labeled cells were then manually marked and counted using the “multi-point” function of image J (V1.8.0, National Institutes of Health, USA). The specificity ratio (%) was calculated as follows: (co-labeled cells of D1R or D2R mRNA and D1-GCaMP6m or D2-GCaMP6m) / (D1-GCaMP6m or D2-GCaMP6m positive cells). The effectiveness ratio (%) was calculated as follows: (co-labeled cells of D1R or D2R mRNA and D1-GCaMP6m or D2-GCaMP6m) / (D1R or D2R mRNA positive cells).

To validate D1R-mCherry or D2R-mCherry antibodies, FISH was conducted using RNA-Protein Co-Detection Ancillary Kit (Advanced Cell Diagnostics 323180) following the manufacturer’s protocol. Briefly, after pretreatment with the RNA-Protein Co-Detection Ancillary Kit, brain slices were incubated in primary antibody (1:300, ab183628, abcom, USA) at 4 °C over night. Sections were then incubated with secondary antibody (anti-rabbit goat conjugated with Alexa Fluor 488, JacksonImmuno, USA, 111-545-003, 1: 500) after hybridization with amplifiers. Sections were counterstained with DAPI (RNAscope® Multiplex Fluorescent Reagent Kit v2, Advanced Cell Diagnostics 323100) for 30 s at room temperature. Glass slides were fixed with antifade solution and coverslipped images were acquired under an Olympus microscope (OLYMPUS BX-43, OLYMPUS, Japan). To verify the specificity of the rAAV-mCherry virus, the MSNs co-labeled by D1R/D2R mRNA and D1-mCherry/D2-mCherry virus were quantified. 20× images including the NAc shell were acquired and boxed areas (300 × 300 μm^2^) were selected. Then, the D1-mCherry/D2-mCherry positive cells, NAc ^D1/D2 mRNA (AF594)^ positive cells, and co-labeled cells were manually marked. These positive or merged cells were counted using the “multi-point” function of image J. The numbers from three representative sections per brain were averaged as the value of each brain. The specificity ratio (%) was calculated as follows: (co-labeled cells of D1R or D2R mRNA and D1-mCherry or D2-mCherry positive cells) / (D1-mCherry or D2-mCherry positive cells).

### Statistical analysis

All data are represented as means ± SEMs. All data were assessed for normality using a one-sample Kolmogorov-Smirnov test, and the Levene’s test was used to confirm homogeneity of variance. Comparisons between two groups were performed either by unpaired or paired t-tests. One-way analyses of variance (ANOVAs), two-way ANOVAs, or two-way repeated-measures ANOVAs were used to compare multiple groups under multiple testing conditions as appropriate. Post-hoc comparisons were conducted using Sidak. Kruskal-Wallis analyses were used to compare data when multiple groups of data did not meet the normality and homogeneity of variance requirements. Statistical procedures were performed using SPSS20.0. All statistical graphs / charts were plotted via GraphPad prism6.0. All experiments and statistical analyses used the double-blind method. Significant levels were set at *p* < 0.05.

---

**Supplementary Table 1.** Statistical Analysis.

| Figure | Part | test type | Exact P-value, F-value with degree of freedom for ANOVAs, t-value with degree of freedom for t-tests |
| --- | --- | --- | --- |
| F1 | F | One-Way Repeated Measures ANOVA | F (2.000, 12.00) = 8.702, p = 0.0001; partner vs. object, p = 0.0060; stranger vs. object, p = 0.0239 |
|  | I | One-Way Repeated Measures ANOVA | F (2.000, 12.00) = 21.41, p = 0.0002; partner vs. stranger, p = 0.0097; partner vs. object, p < 0.0001, stranger vs. object, p = 0.0422 |
| F2 | F | One-Way Repeated Measures ANOVA | F (1.373, 8.235) = 6.662, p = 0.0255; partner vs. stranger, p = 0.2092; partner vs. object, p = 0.0659, stranger vs. object, p = 0.2708 |
|  | I | One-Way Repeated Measures ANOVA | F (1.212, 7.274) = 15.89, p = 0.0039; partner vs. stranger, p = 0.0366; partner vs. object, p = 0.0140, stranger vs. object, p = 0.0434 |
| F3 | F | One-Way Repeated Measures ANOVA | F (2.000, 12.00) = 5.297, p = 0.0184; partner vs. object, p = 0.0249 |
|  | I | One-Way Repeated Measures ANOVA | F (1.242, 7.451) = 11.11, p = 0.0093; partner vs. stranger, p = 0.0278, partner vs. object, p = 0.0326 |
| F4 | C | Two-tailed unpaired t-test | frequency: t (20) = -3.634, p = 0.002; amplitude: t (20) = -2.619, p = 0.016 |
|  | D | Two-tailed unpaired t-test | frequency: t (26) = -2.096, p = 0.046; amplitude: t (26) = -0.528, p = 0.602 |
|  | F | Two-Way Repeated Measures ANOVA | group × treatment: F (8, 198) = 0.070, p = 0.999; group: F (1, 198) = 0.0003, p = 0.986; treatment: F (8, 198) = 16.767, p < 0.0001 |
|  | G | Two-Way Repeated Measures ANOVA | group × treatment: F (8, 198) = 0.103, p = 0.999; group: F (1, 198) = 0.078, p = 0.780; treatment: F (8, 198) = 10.095, p < 0.0001 |
|  | H | Two-tailed unpaired t-test | t (22) = 0.100, p = 0.921 |
|  | J | Two-tailed unpaired t-test | t (10) = -3.499, p = 0.012 |
|  | L | Two-Way Repeated Measures ANOVA with Sidak's multiple comparison test | frequencies: group × treatment: F (1, 14) = 0.0177, p = 0.896; group: F (1, 14) = 8.105, p = 0.0129; treatment: F (1, 14) = 7.189, p = 0.0179, Control_Baseline vs Control DA, p = 0.0050, Cohabitation_Baseline vs Cohabitation DA, p = 0.2296, Control_Baseline vs Cohabitation Baseline, p = 0.0277, Control_DA vs Cohabitation DA, p = 0.0376; amplitude: group × treatment: F (1, 14) = 0.7937, p = 0.4042; group: F (1, 14) = 5.911, p = 0.0291; treatment: F (1, 14) = 14.78, p = 0.0018, Control_Baseline vs Control DA, p = 0.007, Cohabitation Baseline vs Cohabitation DA, p = 0.098, Control Baseline vs Cohabitation Baseline, p = 0.0472, Control DA vs Cohabitation DA, p = 0.0241 |
| F5 | C | Two-tailed unpaired t-test | frequency: t (33) = 2.816, p = 0.008; amplitude: t (33) = 3.322, p = 0.002 |
|  | D | Two-tailed unpaired t-test | frequency: t (24) = 2.324, p = 0.024; amplitude: t (24) = 0.100, p = 0.921 |
|  | F | Two-Way Repeated Measures ANOVA | group × treatment: F (8, 126) = 0.778, p = 0.623; group: F (1, 126) = 7.757, p = 0.006; treatment: F (8, 126) = 29.810, p < 0.0001 |
|  | G | Two-Way Repeated Measures ANOVA | group × treatment: F (8, 126) = 0.845, p = 0.565; group: F (1, 126) = 5.440, p = 0.021; treatment: F (8, 126) = 15.757, p < 0.0001 |
|  | H | Two-tailed unpaired t-test | t (14) = 2.744, p = 0.016 |
|  | J | Two-tailed unpaired t-test | t (12) = -1.429, p = 0.179 |
|  | L | Two-Way Repeated Measures ANOVA with Sidak's multiple comparison test | frequencies: group × treatment: F (1, 18) = 4.784, p = 0.0422; group: F (1, 18) = 9.954, p = 0.0055; treatment: F (1, 18) = 5.164, p = 0.0356, Control_Baseline vs Control DA, p = 0.006, Cohabitation_Baseline vs Cohabitation DA, p = 0.953, Control Baseline vs Cohabitation_Baseline, p = 0.015, Control DA vs Cohabitation DA, p = 0.006; amplitude: group × treatment: F (1, 18) = 0.007, p = 0.9343; group: F (1, 18) = 5.842, p = 0.0265; treatment: F (1, 18) = 0.6498, p = 0.4307, Control_Baseline vs Control DA, p = 0.5285, Cohabitation Baseline vs Cohabitation DA, p = 0.63468, Control Baseline vs Cohabitation Baseline, p = 0.0282, Control DA vs Cohabitation DA, p = 0.029 |
| F6 | E | Two-way ANOVA with Sidak's multiple comparison test | group × treatment: F (2, 40) = 4.942, p = 0.012; group: F (2, 40) = 4.265, p = 0.021; treatment: F (1, 40) = 4.967, p = 0.032; Gi_Saline vs Gi_CNO, p < 0.001; Gq_Saline vs Gq_CNO, p = 0.380; mCherry_Saline vs mCherry_CNO, p = 0.052 |
|  | F | Kruskal-Wallis | H = 4.465, p = 0.215 |
|  | G | Kruskal-Wallis | H = 15.198, p = 0.002, CNO_Partner vs CNO_Stranger, p < 0.0001; saline_Partner vs saline_Stranger, p = 0.328 |
| | H | Two-Way Repeated Measures ANOVA with Sidak's multiple comparison test | group × treatment: F (1, 11) = 2.039, p = 0.181; group: F (1, 11) = 0.135, p = 0.720; treatment: F (1, 11) = 2.293, p = 0.158; CNO_Partner vs CNO_Stranger, p = 0.998; saline_Partner vs saline_Stranger, p = 0.136 |
| | I | Two-way ANOVA with Sidak's multiple comparison test | group × treatment: F (2, 40) = 3.798, p = 0.031; group: F (2, 40) = 6.797, p = 0.003; treatment: F (1, 40) = 3.375, p = 0.074; Gi_Saline vs Gi_CNO, p = 0.861; Gq_Saline vs Gq_CNO, p = 0.002; mCherry_Saline vs mCherry_CNO, p = 0.805 |
|  | J | Kruskal-Wallis | H = 8.083, p = 0.044; CNO_Partner vs CNO_Stranger, p = 0.384; saline_Partner vs saline_Stranger, p = 0.033 |
| | K | Two-Way Repeated Measures ANOVA with Sidak's multiple comparison test | group × treatment: F (1, 15) = 0.002, p = 0.968; group: F (1, 15) = 0.447, p = 0.514; treatment: F (1, 15) = 21.427, p = 0.0003; CNO_Partner vs CNO_Stranger, p = 0.009; saline_Partner vs saline_Stranger, p = 0.012 |
| | L | Two-Way Repeated Measures ANOVA with Sidak's multiple comparison test | group × treatment: F (1, 11) = 0.043, p = 0.839; group: F (1, 11) = 0.306, p = 0.591; treatment: F (1, 11) = 25.166, p = 0.0004; CNO_Partner vs CNO_Stranger, p = 0.009; saline_Partner vs saline_Stranger, p = 0.009 |
| F7 | E | Two-way ANOVA with Sidak's multiple comparison test | group × treatment: F (2, 37) = 0.186, p = 0.831; group: F (2, 37) = 2.729, p = 0.078; treatment: F (1, 37) = 2.839, p = 0.100 |
|  | F | Kruskal-Wallis | H = 0.743, p = 0.863 |
|  | G | Kruskal-Wallis | H = 9.597, p = 0.022; CNO_Partner vs CNO_Stranger, p = 0.087; saline_Partner vs saline_Stranger, p = 0.821 |
| | H | Two-Way Repeated Measures ANOVA with Sidak's multiple comparison test | group × treatment: F (1, 13) = 1.388, p = 0.260; group: F (1, 13) = 0.008, p = 0.928; treatment: F (1, 13) = 5.001, p = 0.044; CNO_Partner vs CNO_Stranger, p = 0.700; saline_Partner vs saline_Stranger, p = 0.071 |
| | I | Two-way ANOVA with Sidak's multiple comparison test | group × treatment: F (2, 37) = 4.934, p = 0.013; group: F (2, 37) = 0.873, p = 0.426; treatment: F (1, 37) = 7.633, p = 0.009; Gq_Saline vs Gq_CNO, p = 0.001; Gi_Saline vs Gi_CNO, p = 0.109; mCherry_Saline vs mCherry_CNO, p = 0.522 |
|  | J | Kruskal-Wallis | H = 12.819, p = 0.005; CNO_Partner vs CNO_Stranger, p = 0.256; saline_Partner vs saline_Stranger, p = 0.001 |
| | K | Two-Way Repeated Measures ANOVA with Sidak's multiple comparison test | group × treatment: F (1, 11) = 4.281, p = 0.063; group: F (1, 11) = 1.513, p = 0.244; treatment: F (1, 11) = 11.027, p = 0.007; CNO_Partner vs CNO_Stranger, p = 0.377; saline_Partner vs saline_Stranger, p = 0.004 |
| | L | Two-Way Repeated Measures ANOVA with Sidak's multiple comparison test | group × treatment: F (1, 13) = 0.033, p = 0.859; group: F (1, 13) = 0.325, p = 0.578; treatment: F (1, 13) = 15.559, p = 0.002; CNO_Partner vs CNO_Stranger, p = 0.033; saline_Partner vs saline_Stranger, p = 0.029 |
| FS1 | B | Paired t test | t (8) = 0.623, p = 0.551 |
|  | C | Paired t test | t (8) = 5.294, p = 0.0007 |
| FS2 | E | One-Way Repeated Measures ANOVA | F (2.000, 8.00) = 4.092, p = 0.0597 |
|  | G | One-Way Repeated Measures ANOVA | F (2.000, 8.00) = 0.6107, p = 0.5665 |
|  | I | One-Way Repeated Measures ANOVA | F (2.000, 8.00) = 0.0800, p = 0.9238 |
|  | K | One-Way Repeated Measures ANOVA | F (2.000, 8.00) = 2.786, p = 0.1207 |
| FS3 | F | One-Way Repeated Measures ANOVA | F (2.000, 12.00) = 19.59, p = 0.0002; partner vs. object, p = 0.0004, stranger vs. object, p = 0.0006 |
|  | I | One-Way Repeated Measures ANOVA | F (2.000, 12.00) = 10.85, p = 0.0087; partner vs. object, p = 0.0056, stranger vs. object, p = 0.0044 |
|  | **J** | **Two-Way Repeated Measures ANOVA** | **group × treatment: $F(2, 12) = 0.5558, p = 0.5877$; group: $F(2, 12) = 20.53, p = 0.0001$; treatment: $F(1, 6) = 0.1467, p = 0.7149$** |
| **FS4** | **A** | partner: Paired t test | $t(6) = 3.290, p = 0.0166$ |
| | | stranger: Paired t test | $t(6) = 4.884, p = 0.0028$ |
| | | object: Paired t test | $t(6) = 1.745, p = 0.1315$ |
| | **B** | partner: Paired t test | $t(6) = 6.020, p = 0.0009$ |
| | | stranger: Paired t test | $t(6) = 2.611, p = 0.0401$ |
| | | object: Paired t test | $t(6) = 2.209, p = 0.0692$ |
| | **C** | partner: Paired t test | $t(6) = 2.936, p = 0.0261$ |
| | | stranger: Paired t test | $t(6) = 2.499, p = 0.0466$ |
| | | object: Paired t test | $t(6) = 0.8186, p = 0.4443$ |
| | **D** | partner: Paired t test | $t(6) = 3.792, p = 0.0091$ |
| | | stranger: Paired t test | $t(6) = 3.935, p = 0.0077$ |
| | | object: Paired t test | $t(6) = 0.1841, p = 0.8600$ |
| | **E** | partner: Paired t test | $t(6) = 2.700, p = 0.0356$ |
| | | stranger: Paired t test | $t(6) = 2.492, p = 0.0470$ |
| | | object: Paired t test | $t(6) = 0.5216, p = 0.6207$ |
| | **F** | partner: Paired t test | $t(6) = 3.591, p = 0.0115$ |
| | | stranger: Paired t test | $t(6) = 2.381, p = 0.0547$ |
| | | object: Paired t test | $t(6) = 1.590, p = 0.1630$ |
| **FS5** | **A** | cagemate: Paired t test | $t(5) = 1.464, p = 0.2030$ |
| | | stranger: Paired t test | $t(5) = 3.327, p = 0.0209$ |
| | | object: Paired t test | $t(5) = 1.249, p = 0.2669$ |
| | **B** | cagemate: Paired t test | $t(5) = 1.998, p = 0.1021$ |
| | | stranger: Paired t test | $t(5) = 2.602, p = 0.0481$ |
| | | object: Paired t test | $t(5) = 0.1254, p = 0.9051$ |
| | **C** | cagemate: Paired t test | $t(5) = 2.479, p = 0.00559$ |
| | | stranger: Paired t test | $t(5) = 2.654, p = 0.0452$ |
| | | object: Paired t test | $t(5) = 0.5164, p = 0.6276$ |
| | **D** | cagemate: Paired t test | $t(5) = 3.267, p = 0.0223$ |
| | | stranger: Paired t test | $t(5) = 2.789, p = 0.0385$ |
| | | object: Paired t test | $t(5) = 1.035, p = 0.3479$ |
| | **E** | cagemate: Paired t test | $t(5) = 1.788, p = 0.1338$ |
| | | stranger: Paired t test | $t(5) = 2.613, p = 0.0475$ |
| | | object: Paired t test | $t(5) = 0.4349, p = 0.6818$ |
| | **F** | cagemate: Paired t test | $t(5) = 2.009, p = 0.1008$ |
| | | stranger: Paired t test | $t(5) = 2.983, p = 0.0307$ |
| | | object: Paired t test | $t(5) = 1.049, p = 0.3420$ |
| **FS6** | **F** | One-Way Repeated Measures ANOVA | $F(2.000, 10.00) = 4.864, p = 0.0335$; stranger vs. object, $p = 0.0396$ |
| | **I** | One-Way Repeated Measures ANOVA | $F(2.000, 10.00) = 11.42, p = 0.0026$; cagemate vs. object, $p = 0.0074$; stranger vs. object, $p = 0.0050$ |
| **FS7** | **D** | Two-Way Repeated Measures ANOVA | group × treatment: $F(1, 7) = 0.0274, p = 0.8732$; group: $F(1, 7) = 0.0557, p = 0.8201$; treatment: $F(1, 7) = 32.79, p = 0.0007$ |
| | **G** | Two-Way Repeated Measures ANOVA | group × treatment: $F(1, 5) = 0.7013, p = 0.4405$; group: $F(1, 5) = 0.6822, p = 0.4464$; treatment: $F(1, 5) = 42.67, p = 0.0013$ |
| | **J** | Two-Way Repeated Measures ANOVA | group × treatment: $F(1, 7) = 1.1413, p = 0.2733$; group: $F(1, 7) = 1.428, p = 0.2711$; treatment: $F(1, 7) = 8.499, p = 0.0225$ |
| **FS8** | **A** | Two-tailed unpaired t-test | $t(6) = 0.08005, p = 0.9388$ |
| | **B** | Two-tailed unpaired t-test | $t(9) = 2.733, p = 0.0231$ |
| | **C** | Two-tailed paired t-test | $t(6) = 0.6936, p = 0.5139$ |
| | **D** | Two-tailed unpaired t-test | $t(10) = 2.813, p = 0.0184$ |
| | **E** | Two-tailed unpaired t-test | $t(6) = 1.943, p = 0.1000$ |
| | **F** | Two-tailed unpaired t-test | $t(7) = 2.392, p = 0.0481$ |
| FS9 | F | One-Way Repeated Measures ANOVA | $F(2.000, 10.00) = 4.664, p = 0.0371$; stranger vs. object, $p = 0.0498$ |
| | I | One-Way Repeated Measures ANOVA | $F(2.000, 10.00) = 4.940, p = 0.0322$; stranger vs. object, $p = 0.0426$ |
| FS10 | F | One-Way Repeated Measures ANOVA | $F(2.000, 10.00) = 2.054, p = 0.1790$; |
| | I | One-Way Repeated Measures ANOVA | $F(2.000, 10.00) = 5.533, p = 0.0241$; stranger vs. object, $p = 0.0278$ |
| FS11 | E | One-Way Repeated Measures ANOVA | $F(2.000, 8.00) = 0.7086, p = 0.5208$ |
| | G | One-Way Repeated Measures ANOVA | $F(2.000, 8.00) = 1.450, p = 0.2901$ |
| | I | One-Way Repeated Measures ANOVA | $F(2.000, 8.00) = 0.7471, p = 0.5041$ |
| | K | One-Way Repeated Measures ANOVA | $F(2.000, 8.00) = 1.344, p = 0.3140$ |
| FS12 | E | One-Way Repeated Measures ANOVA | $F(2.000, 8.00) = 4.162, p = 0.0577$ |
| | G | One-Way Repeated Measures ANOVA | $F(2.000, 8.00) = 0.6909, p = 0.5287$ |
| | I | One-Way Repeated Measures ANOVA | $F(2.000, 8.00) = 0.5571, p = 0.5936$ |
| | K | One-Way Repeated Measures ANOVA | $F(2.000, 8.00) = 2.178, p = 0.1757$ |
| FS13 | A | Two-Way Repeated Measures ANOVA | group × treatment: $F(3, 18) = 0.3050, p = 0.8214$; group: $F(3, 18) = 0.7172, p = 0.5545$; treatment: $F(1, 6) = 2.757, p = 0.1479$ |
| | B | Two-Way Repeated Measures ANOVA | group × treatment: $F(3, 18) = 0.8046, p = 0.5076$; group: $F(3, 18) = 1.891, p = 0.1672$; treatment: $F(1, 6) = 0.6753, p = 0.4426$ |
| | C | Two-Way Repeated Measures ANOVA | group × treatment: $F(3, 18) = 0.3340, p = 0.8009$; group: $F(3, 18) = 1.527, p = 0.2418$; treatment: $F(1, 6) = 0.3764, p = 0.5621$ |
| | D | Two-Way Repeated Measures ANOVA | group × treatment: $F(2, 12) = 11.50, p = 0.0016$; group: $F(2, 12) = 25.43, p < 0.0001$; treatment: $F(1, 6) = 18.52, p = 0.0051$ |
| | E | Two-Way Repeated Measures ANOVA | group × treatment: $F(2, 12) = 6.054, p = 0.0152$; group: $F(2, 12) = 15.51, p = 0.0005$; treatment: $F(1, 6) = 4.363, p = 0.0817$ |
| | F | Two-Way Repeated Measures ANOVA | group × treatment: $F(2, 12) = 4.496, p = 0.0349$; group: $F(2, 12) = 11.67, p = 0.0015$; treatment: $F(1, 6) = 7.372, p = 0.0349$ |
| | G | Two-Way Repeated Measures ANOVA | group × treatment: $F(2, 10) = 0.3067, p = 0.7425$; group: $F(2, 10) = 10.83, p = 0.0031$; treatment: $F(1, 5) = 0.6553, p = 0.4550$ |
| | H | Two-Way Repeated Measures ANOVA | group × treatment: $F(2, 10) = 0.0211, p = 0.9792$; group: $F(2, 10) = 8.527, p = 0.0069$; treatment: $F(1, 5) = 0.2301, p = 0.6517$ |
| | I | Two-Way Repeated Measures ANOVA | group × treatment: $F(2, 10) = 1.406, p = 0.2897$; group: $F(2, 10) = 5.215, p = 0.0281$; treatment: $F(1, 5) = 1.111, p = 0.3400$ |

## Supplemental information

**Figure S1.**
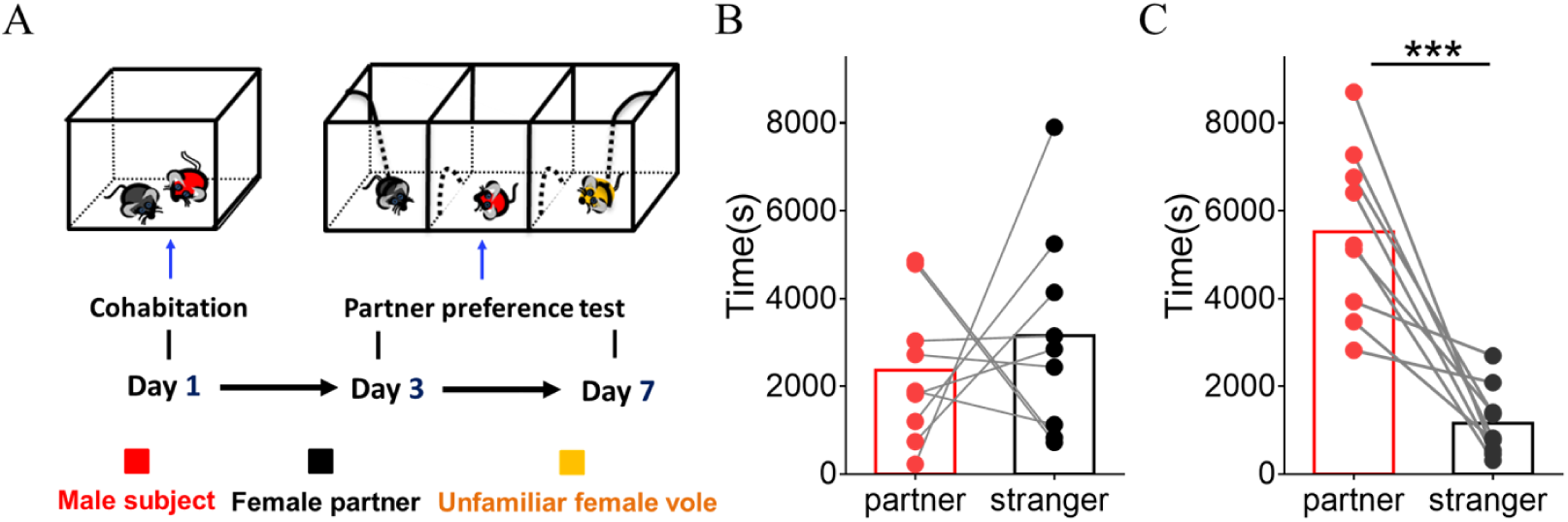
Mandarin voles showed significant preference to partners in the partner preference test after cohabitation. (A) Timeline of experiments. (B) Quantification of side-by-side time in the partner preference test after cohabitation for 3 days (n = 9 voles, Paired t test: t (8) = 0.623, p = 0.551). (C) Quantification of side-by-side time in the partner preference test after 7 days of cohabitation (n = 9 voles, Paired t test: t (8) = 5.294, p = 0.0007). Error bars = SEM. *** represent *p* < 0.001.

**Figure S2.**
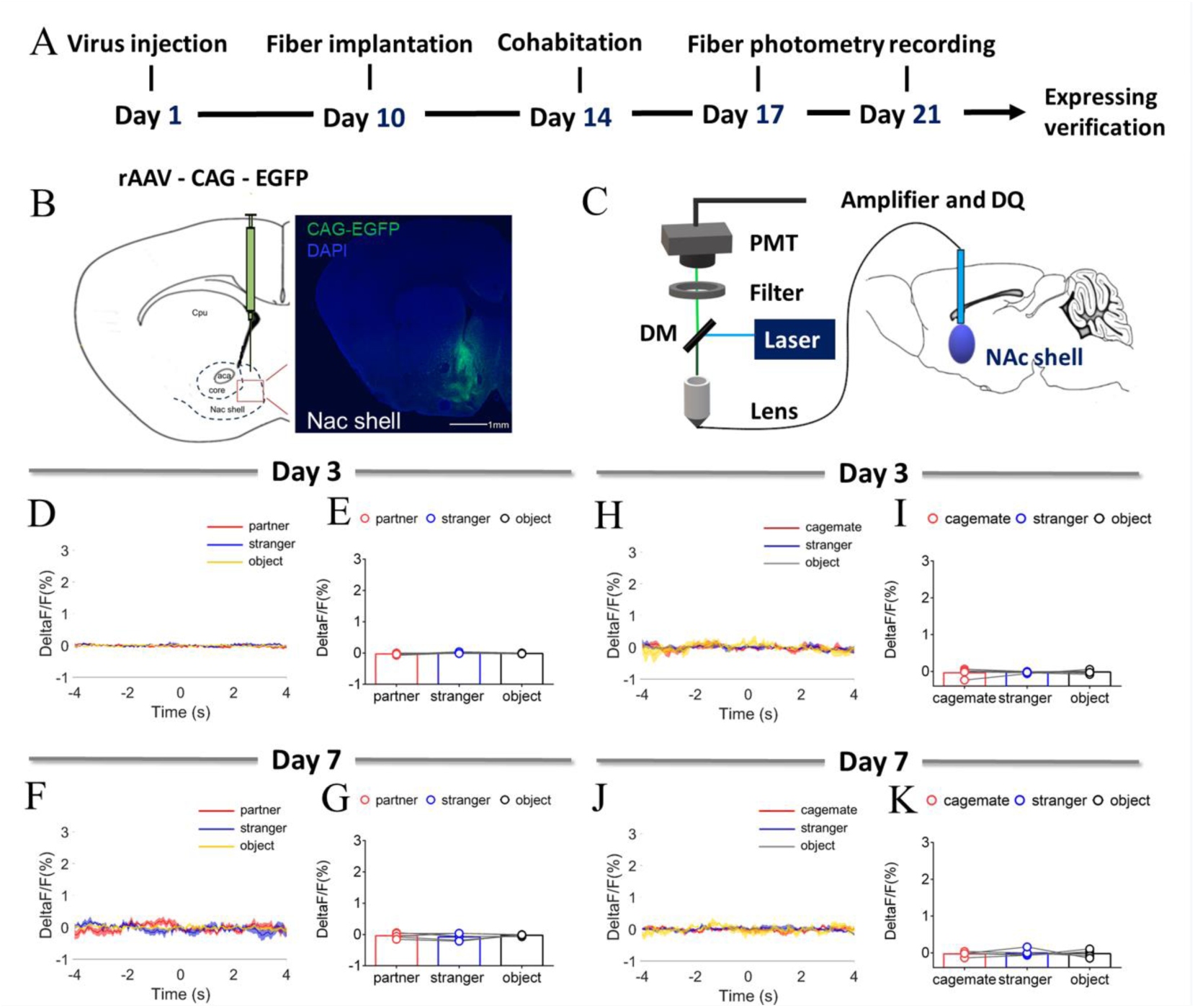
NAc shell EGFP fluorescence signals (with injection of control virus) upon sniffing partner or an unknown female after cohabitation. (A)Timeline of experiments. (B) Schematic diagrams depicting histology showing the expression of EGFP within the NAc shell. Scale bar: 1 mm. (C) The schematic of the fiber photometry. (D, F) Mean fluorescence signals changes of EGFP fluorescence signals during sniffing partner (red line), an unknown female (blue line) or an object (yellow line) after cohabitation for 3 days (D) and 7 days (F). (E, G) Quantification (Repeated One-way ANOVA) of change in EGFP fluorescence signals during sniffing partner, an unknown female or an object after cohabitation for 3 days (E) (n = 5 voles) and 7 days (G) (n = 5 voles). (H, J) Mean fluorescence signal changes of EGFP fluorescence signals during sniffing cagemate (red line), an unknown male (blue line) or an object (yellow line) after cohabitation for 3 days (H) and 7 days (J). (I, K) Quantification (Repeated One-way ANOVA) of change in EGFP fluorescence signals during sniffing cagemate, an unknown male or an object after cohabitation for 3 days (I) (n = 5 voles) and 7 days (K) (n = 5 voles). All error bars = SEM. Supplementary Table 1 for detailed statistics.

**Figure S3.**
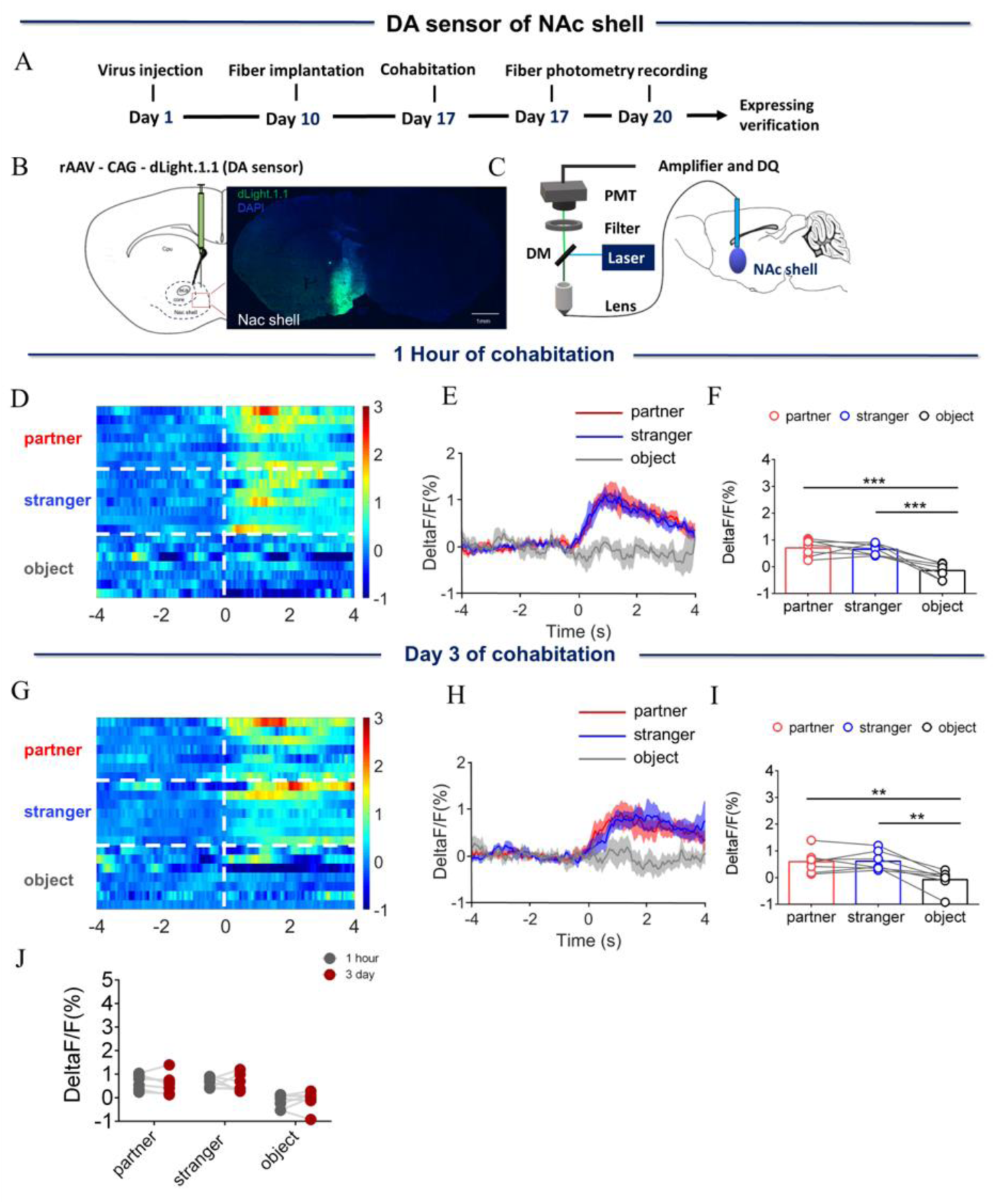
Dynamics of extracellular dopamine (DA) concentration within the nucleus accumbens (NAc) shell upon sniffing their partner or an unknown female after one hour or three days of cohabitation. (A) Timeline of experiments. (B) Schematic diagrams depicting virus injection and recording sites and histology showing the expression of DA sensor within the NAc shell. Scale bar: 1 mm. (C) Schematic of the procedure used to record extracellular DA concentration in the NAc shell using fiber photometry. (D) Heat map illustrating the extracellular DA concentrations (ΔF/F, %) of the NAc shell when sniffing their partner, an unknown female, and an unrelated object. (E) Mean fluorescence signal changes of DA sensor during sniffing their partner (red line), an unknown female (blue line), or an object (gray line) after one hour of cohabitation. The shaded area along the differently colored lines represent the margin of error. (F) Quantification (Repeated One-way ANOVA) of changes in DA signals during sniffing of their partner, an unknown female, and an object after one hour of cohabitation. (G) Heat map illustrating the extracellular DA concentration (ΔF/F, %) of the NAc shell when sniffing their partner, an unknown female, and an object. (H) Mean fluorescence signal changes of the DA sensor when sniffing their partner (red line), an unknown female (blue line), or an object (gray line) after 3 days of cohabitation. (I) Quantification (Repeated One-way ANOVA) of changes in extracellular DA concentration when sniffing their partner, an unknown female, and an object after 3 days of cohabitation. (J) Quantification (Two-Way Repeated Measures ANOVA) change sniffing their partner, an unknown female, and an object after cohabitation for one hour and 3 days (n= 7 voles). Error bars = SEM. * represents *p* < 0.05, ** represents *p* < 0.01, and *** represents *p* < 0.001. See Supplementary Table 1 for detailed statistics.

**Figure S4.**
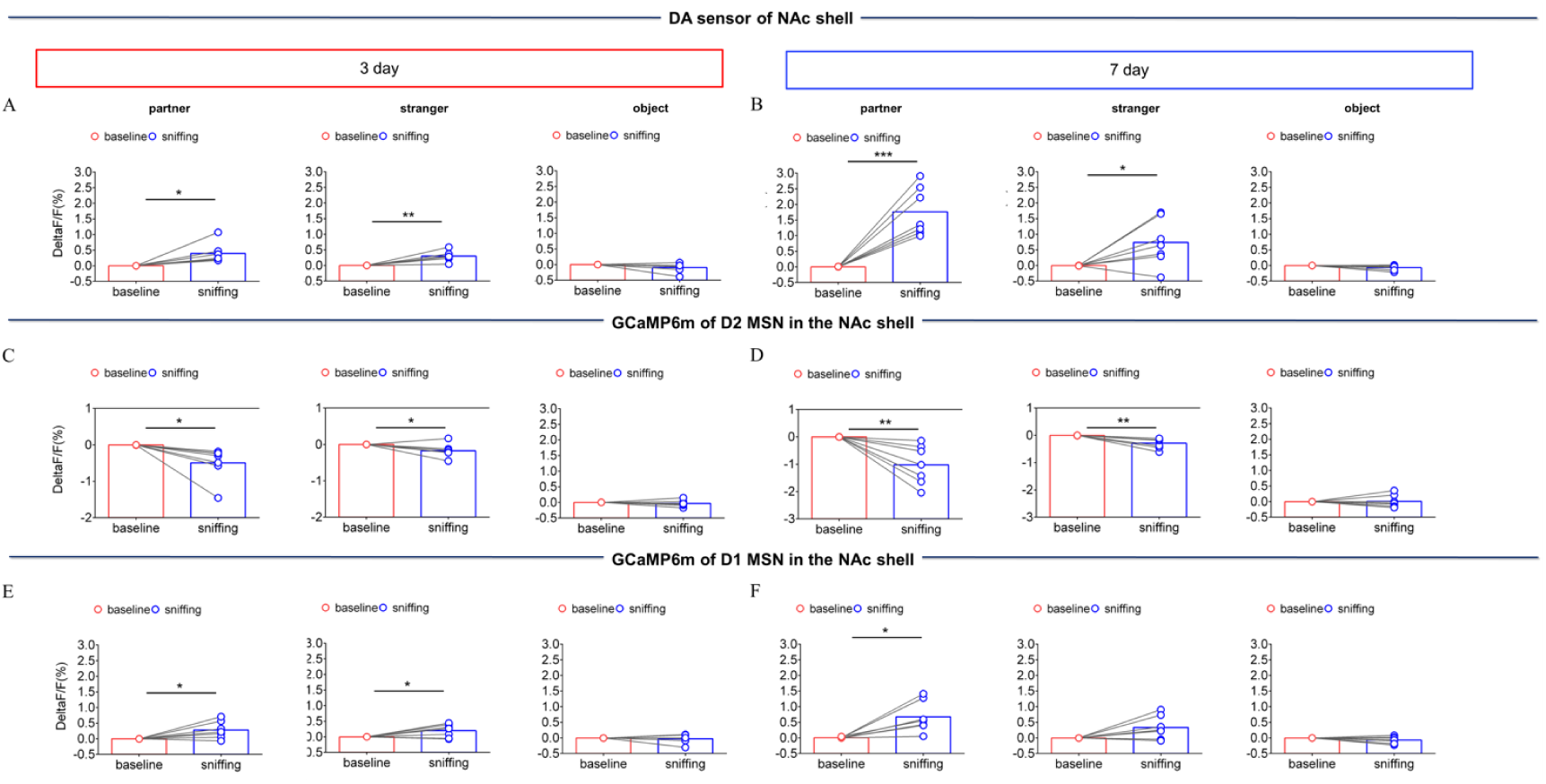
NAc shell Dynamics of extracellular DA concentration, D2 MSNs and D1 MSNs when sniffing their partner or an unknown female after cohabitation. (A) DA sensor: Quantification (Paired t test) of change during sniffing partner (left), female stranger (middle) or object (right) after cohabitation for 3 days (n= 7 voles). (B) DA sensor: Quantification (Paired t test) of changes during sniffing partner (left), an unknown female (middle) or an object (right) after cohabitation for 7 days (n= 7 voles). (C) D2 MSNs: Quantification (Paired t test) of change in calcium signals during sniffing partner (left), an unknown female (middle) or an object (right) after cohabitation for 3 days (n= 7 voles). (D) D2 MSNs: Quantification (Paired t test) of change in calcium signals during sniffing partner (left), an unknown female (middle) or an object (right) after cohabitation for 7 days (n= 7 voles). (E) D1 MSNs: Quantification (Paired t test) of change in calcium signals during sniffing partner (left), an unknown female (middle) or an object (right) after cohabitation for 3 days (n= 7 voles). (F) D1 MSNs: Quantification (Paired t test) of change in calcium signals during sniffing partner (left), an unknown female (middle) or an object (right) after cohabitation for 7 days (n= 7 voles). Error bars = SEM * represents *p* < 0.05, ** represents *p* < 0.01. See Supplementary Table 1 for detailed statistics.

**Figure S5.**
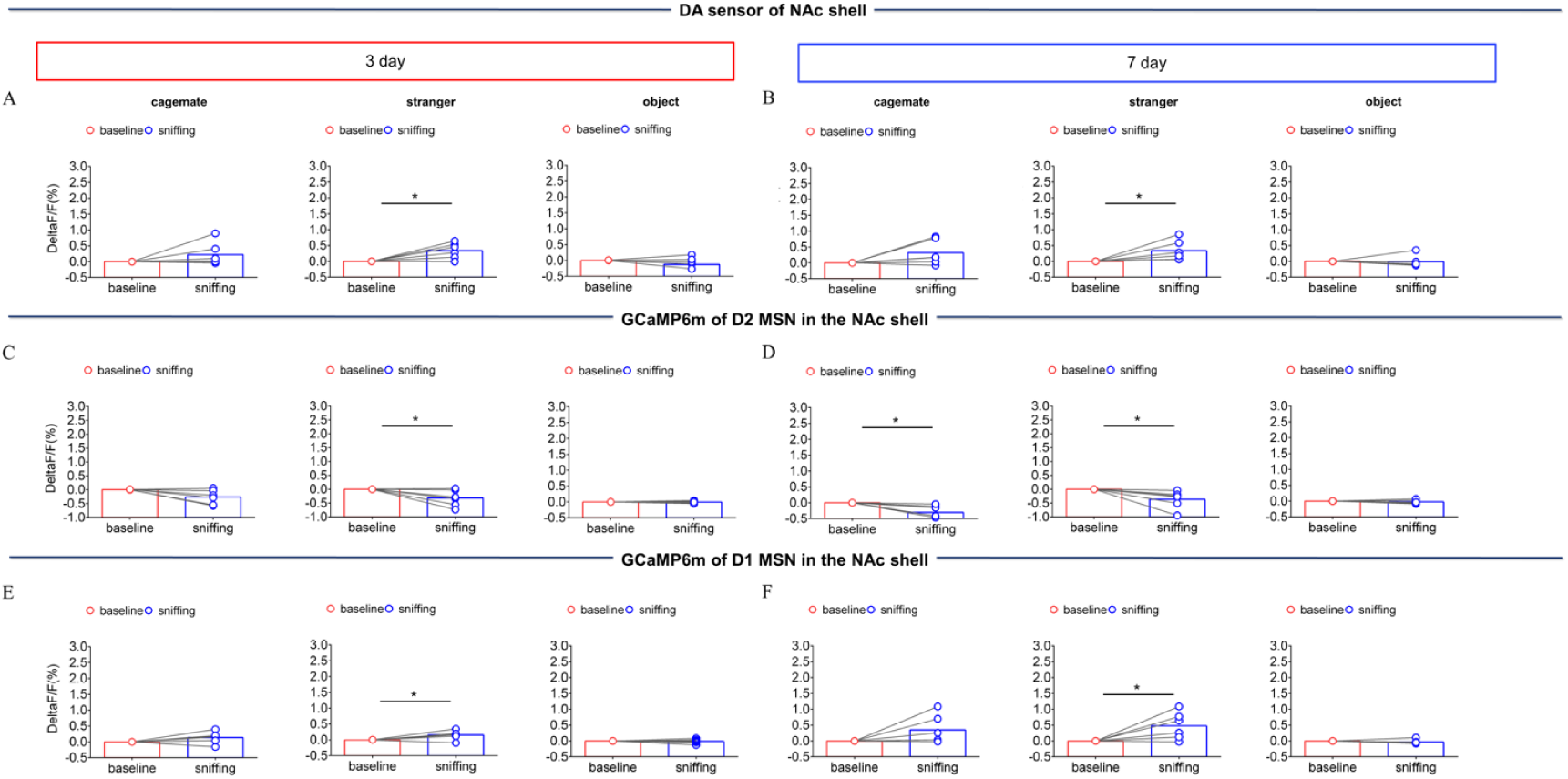
NAc shell Dynamics of extracellular DA concentration, D2 MSNs and D1 MSNs when sniffing their cagemate or an unknown male after cohabitation. (A) DA sensor: Quantification (Paired t test) of changes during sniffing cagemate (left), an unknown male (middle) or an object (right) after cohabitation for 3 days (n= 6 voles). (B) DA sensor: Quantification (Paired t test) of changes during sniffing cagemate (left), an unknown male (middle) or an object (right) after cohabitation for 7 days (n= 6 voles). (C) D2 MSNs: Quantification (Paired t test) of change in calcium signals during sniffing cagemate (left), an unknown male (middle) or an object (right) after cohabitation for 3 days (n= 6 voles). (D) D2 MSNs: Quantification (Paired t test) of change in calcium signals during sniffing cagemate (left), an unknown male (middle) or an object (right) after cohabitation for 7 days (n= 6 voles). (E) D1 MSNs: Quantification (Paired t test) of change in calcium signals during sniffing cagemate (left), an unknown male (middle) or an object (right) after cohabitation for 3 days (n= 6 voles). (F) D1 MSNs: Quantification (Paired t test) of change in calcium signals during sniffing cagemate (left), an unknown male (middle) or an object (right) after cohabitation for 7 days (n= 6 voles). Error bars = SEM. * represent *p* < 0.05. Supplementary Table 1 for detailed statistics.

**Figure S6.**
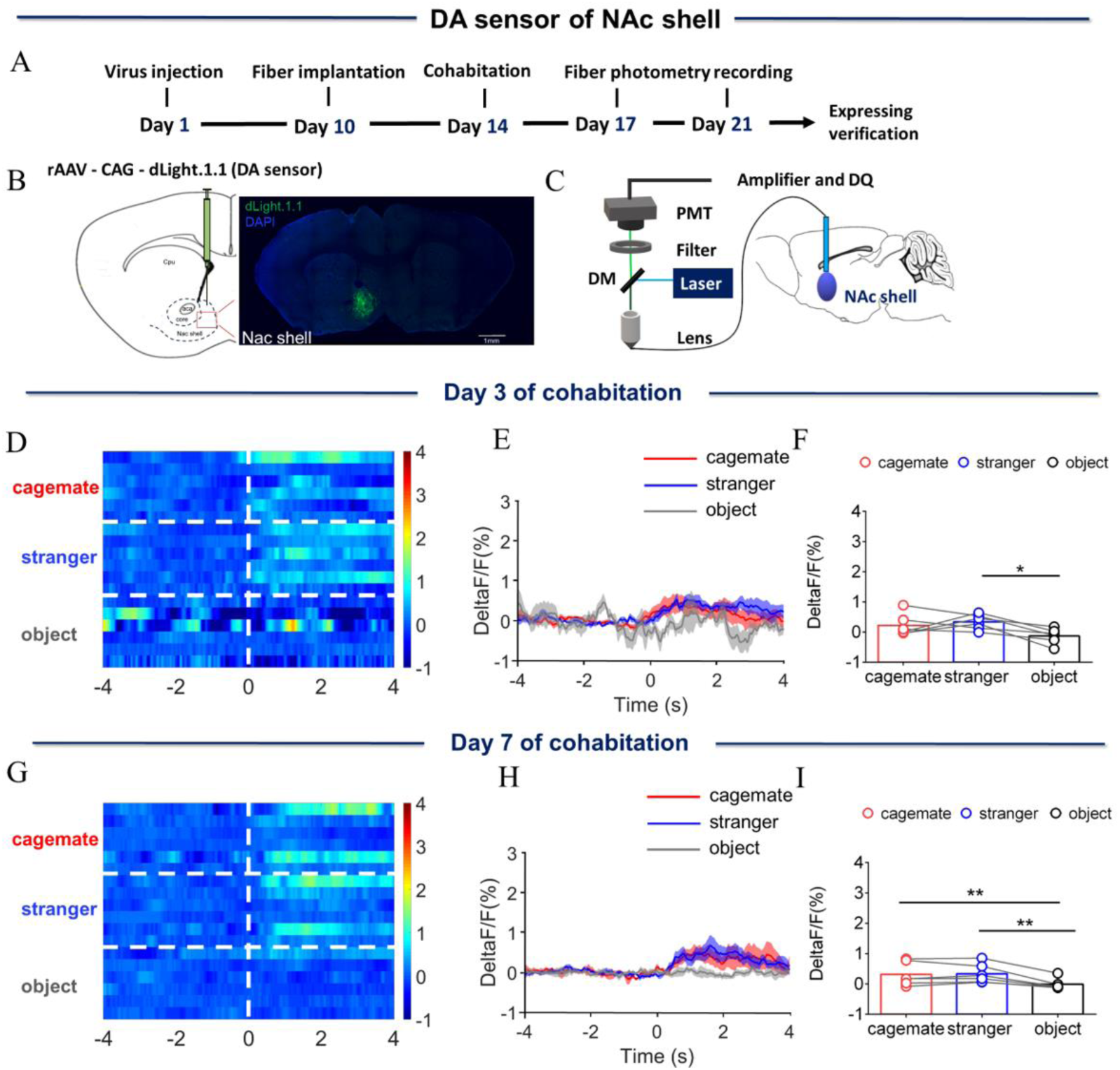
Dynamics of extracellular DA concentration within the NAc shell upon sniffing their cagemate or an unknown male. (A) Timeline of experiments. (B) Schematic diagrams depicting virus injection and recording sites and histology showing the expression of DA sensor within the NAc shell. Scale bar: 1 mm. (C) Schematic of the procedure used to record extracellular DA concentration in the NAc shell using fiber photometry. (D) Heat map illustrating the extracellular DA concentrations (ΔF/F, %) of the NAc shell when sniffing their cagemate, an unknown male, or an object. (E) Mean fluorescence signal changes of DA sensor during sniffing their cagemate (red line), an unknown male (blue line), or an object (gray line) after 3 days of cohabitation. The shaded area along the differently colored lines represent the margin of error. (F) Quantification (repeated one-way ANOVA) of changes in DA signals during sniffing of their cagemate, an unknown male, or an object after 3 days of cohabitation. (G) Heat map illustrating the extracellular DA concentration (ΔF/F, %) of the NAc shell when sniffing their cagemate, an unknown male, or an object. (H) Mean fluorescence signal changes of the DA sensor when sniffing their cagemate (red line), an unknown male (blue line), or an object (gray line) after 7 days of cohabitation. (I) Quantification (repeated one-way ANOVA) of changes in extracellular DA concentration when sniffing their cagemate, an unknown male, or an object after 7 days of cohabitation. Error bars = SEM. * represents *p* < 0.05, ** represents *p* < 0.01. See Supplementary Table 1 for detailed statistics.

**Figure S7.**
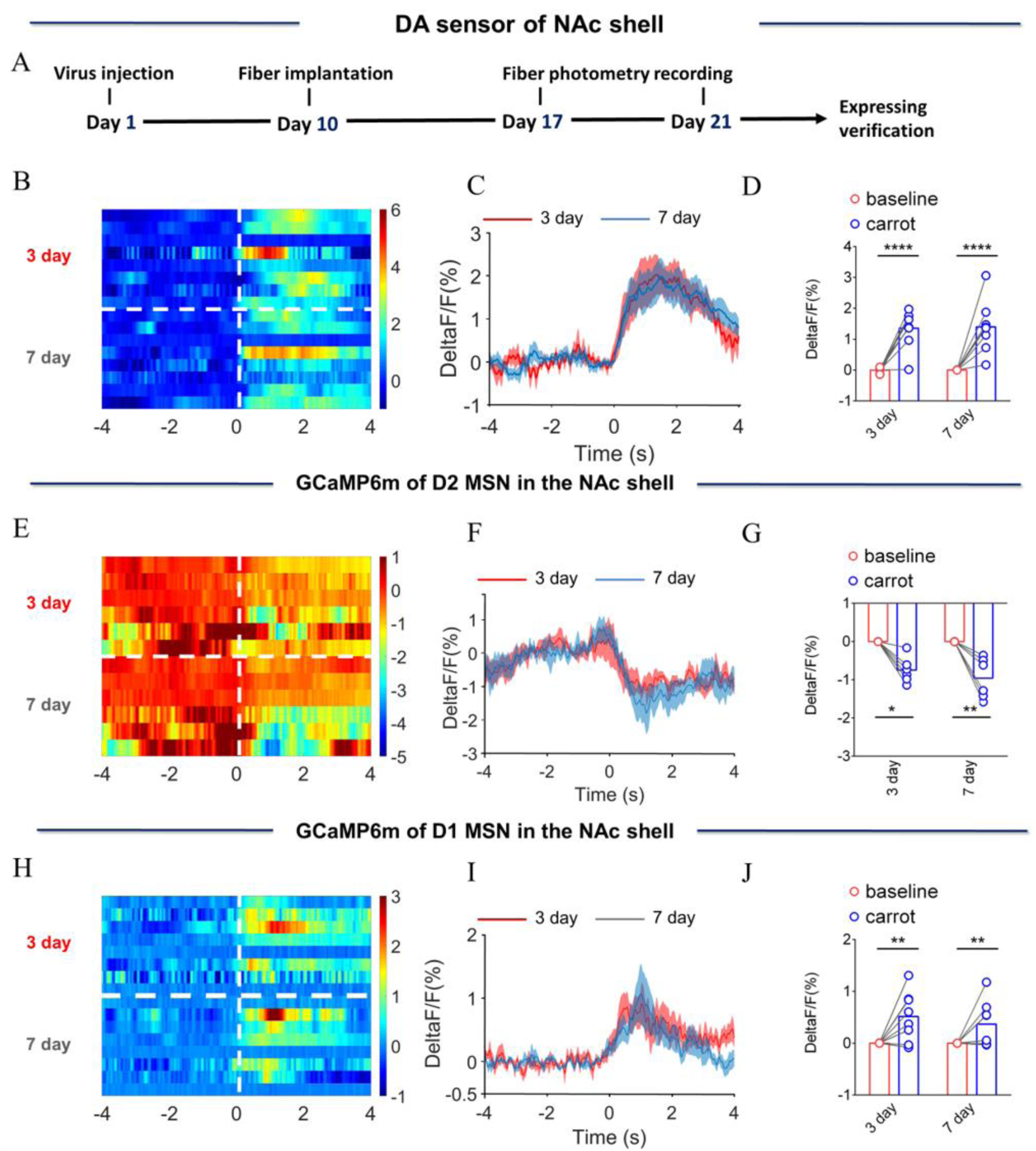
NAc shell Dynamics of extracellular DA concentration, D2 MSNs and D1 MSNs upon eating carrot. (A)Timeline of experiments. (B) Heat map illustrating the extracellular DA concentration (ΔF/F, %) of NAc shell upon eating carrot after cohabitation for 3 days and 7 days. (C) Mean fluorescence signals changes of DA sensor upon eating carrot after cohabitation for 3 (red line) and 7 days (blue line). The shaded area along the different colored lines represent error margins. (D) Quantification (Two-Way Repeated Measures ANOVA) of change in DA signals upon eating carrot after cohabitation for 3 and 7 days. (E) Heat map illustrating the extracellular D2 MSNs (ΔF/F, %) of NAc shell upon eating carrot after cohabitation for 3 and 7 days. (F) Mean fluorescence signals changes of D2 MSNs upon eating carrot after cohabitation for 3 (red line) and 7 days (blue line). The shaded area along the different colored lines represent error margins. (G) Quantification (Two-Way Repeated Measures ANOVA) of change in D2 MSNs signals upon eating carrot after cohabitation for 3 and 7 days. (H) Heat map illustrating the extracellular D1 MSNs (ΔF/F, %) of NAc shell upon eating carrot after cohabitation for 3 and 7 days. (I) Mean fluorescence signals changes of D1 MSNs upon eating carrot after cohabitation for 3 (red line) and 7 days (blue line). The shaded area along the different colored lines represents error margins. (J) Quantification (Two-Way Repeated Measures ANOVA) of change in D1 MSNs upon eating carrot after cohabitation for 3 and 7 days. All error bars = SEM. * represent *p* < 0.05, ** represent *p* < 0.01, *** represent *p* < 0.001. Supplementary Table 1 for detailed statistics.

**Figure S8.**
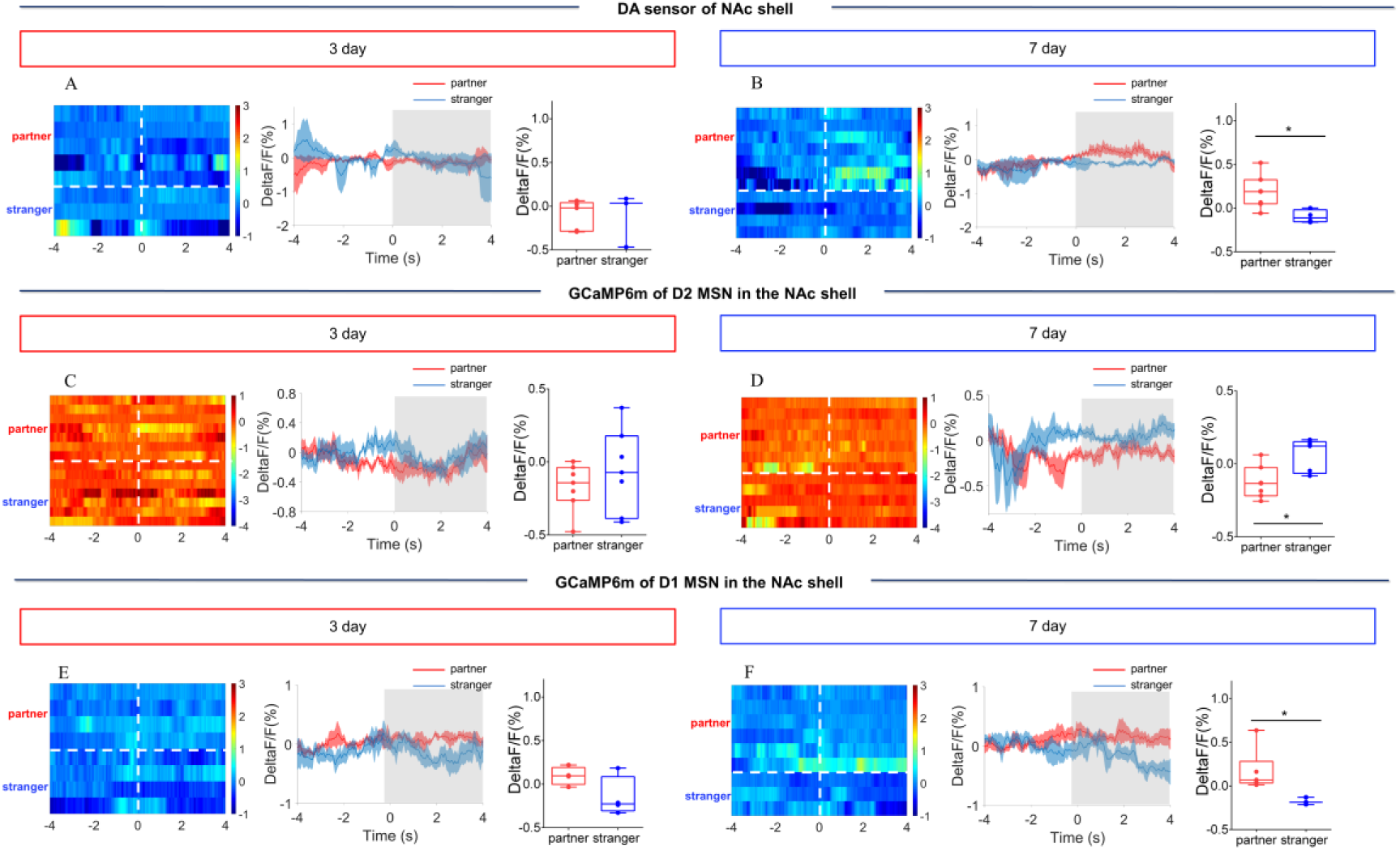
NAc shell Dynamics of extracellular DA release, D2 MSNs and D1 MSNs during side-by-side contact after cohabitation. (A) Mean fluorescence signals changes (middle) and quantification (unpaired t test) (right) of DA sensor during side-by-side contact with partner (n= 5 voles) or an unknown female (n= 3 voles) after 3 days of cohabitation. Left: Heat map illustrating the extracellular DA concentration (ΔF/F, %) of NAc shell during side-by-side contact after cohabitation for 3 days. (B) Mean fluorescence signals changes (middle) and quantification (unpaired t test) (right) of DA sensor during side-by-side contact with partner (n= 7 voles) or an unknown female (n= 4 voles) after 7 days of cohabitation. Left: Heat map illustrating the extracellular DA concentration (ΔF/F, %) of NAc shell during side-by-side contact after cohabitation for 7 days. (C) Mean fluorescence signals changes (middle) and quantification (paired t test) (right) of D2 MSNs during side-by-side contact with partner (n= 7 voles) or an unknown female (n= 7 voles) after 3 days of cohabitation. Left: Heat map illustrating the fluorescence signals of NAc shell D2 MSNs during side-by-side contact after cohabitation for 3 days. (D) Mean fluorescence signals changes (middle) and quantification (unpaired t test) (right) of D2 MSNs during side-by-side contact with partner (n= 7 voles) or an unknown female (n= 5 voles) after 7 days of cohabitation. Left: Heat map illustrating the fluorescence signals of NAc shell D2 MSNs during side-by-side contact after cohabitation for 7 days. (E) Mean fluorescence signals changes (middle) and quantification (unpaired t test) (right) of D1 MSNs during side-by-side contact with partner (n= 4 voles) or an unknown female (n= 4 voles) after 3 days of cohabitation. Left: Heat map illustrating the fluorescence signals of NAc shell D1 MSNs during side-by-side contact after cohabitation for 3 days. (F) Mean fluorescence signals changes (middle) and quantification (unpaired t test) (right) of D1 MSNs during side-by-side contact with partner (n= 6 voles) or an unknown female (n= 3 voles) after 7 days of cohabitation. Left: Heat map illustrating the fluorescence signals of NAc shell D1 MSNs during side-by-side contact after cohabitation for 7 days. Not all animals engaged in side-by-side contact. The number of data points was different from numbers of subjects because some animals did nor showed side-by-side contact, thus we could not detect the changes in fluorescence signals upon occurring of this behavior. All error bars = SEM. * represent *p* < 0.05. Supplementary Table 1 for detailed statistics.

**Figure S9.**
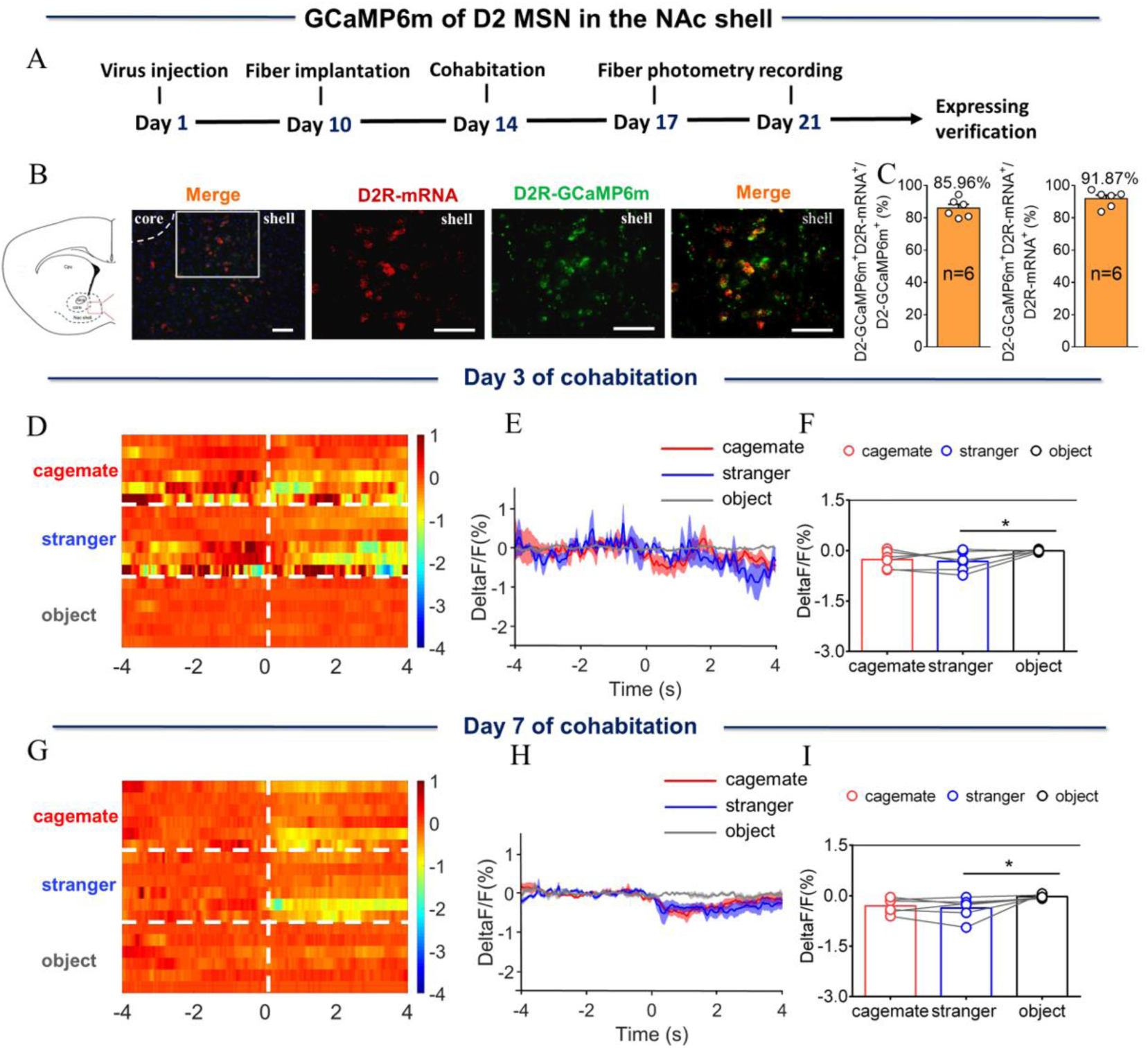
NAc shell D2 MSNs showing decreased activity upon sniffing their cagemate or an unknown male after cohabitation. (A)Timeline of experiments. (B) Overlap of D2-GCaMP6m (green), D2R-mRNA (red), and DAPI (blue) in the NAc shell. Scale bar: 100 μm. (C) Statistical chart showed that D2-GCaMP6m was relatively restricted to D2R-mRNA positive neurons (n=6 voles). (D) Heat map illustrating the calcium signals (ΔF/F, %) of NAc shell during sniffing cagemate, an unknown male or an object. (E) Mean fluorescence signals changes of D2 MSNs during sniffing cagemate (red line), an unknown male (blue line) or an object (gray line) after 3 days of cohabitation. (F) Quantification (Repeated One-way ANOVA) of change in fluorescence signals during sniffing cagemate, an unknown male or an object after 3 days of cohabitation (n=6 voles). (G) Heat map illustrating the calcium signals (ΔF/F, %) of NAc shell during sniffing cagemate, an unknown male or an object. (H) Mean fluorescence signal changes of D2 MSNs during sniffing cagemate (red line), an unknown male (blue line) or an object (gray line) after 7 days of cohabitation. (I) Quantification (Repeated One-way ANOVA) of change in fluorescence signals during sniffing cagemate, an unknown male or an object after 7 days of cohabitation (n=6 voles). All error bars = SEM. * represents *p* < 0.05. Supplementary Table 1 for detailed statistics.

**Figure S10.**
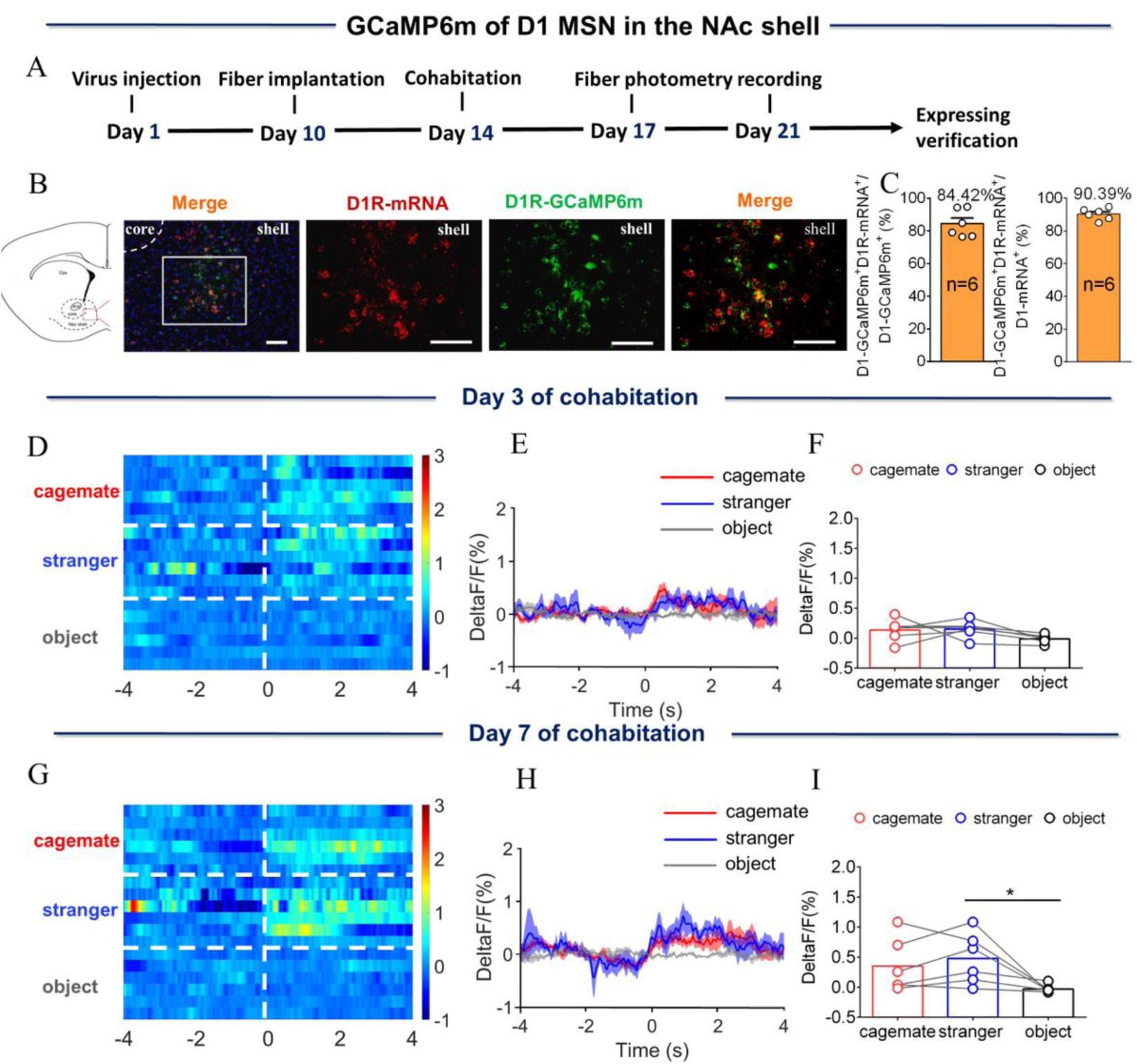
NAc shell D1 MSNs showing increased activity when sniffing their cagemate or an unknown male after cohabitation. (A)Timeline of experiments. (B) Overlap of D1-GCaMP6m (green), D1R-mRNA (red), and DAPI (blue) in the NAc shell. Scale bar: 100 μm. (C) Statistical chart showed that D1-GCaMP6m was relatively restricted to D1R-mRNA positive neurons (n=6 voles). (D) Heat map illustrating the calcium signals (ΔF/F, %) of NAc shell during sniffing cagemate, an unknown male or an object. (E) Mean fluorescence signals changes of D1 MSNs during sniffing cagemate (red line), an unknown male (blue line) or an object (gray line) after 3 days of cohabitation. (F) Quantification (Repeated One-way ANOVA) of change in fluorescence signal during sniffing cagemate, an unknown male or an object after 3 days of cohabitation (n=6 voles). (G) Heat map illustrating the calcium signals (ΔF/F, %) of NAc shell during sniffing cagemate, an unknown male or an object. (H) Mean fluorescence signal changes of D1 MSNs during sniffing cagemate (red line), an unknown male (blue line) or an object (gray line) after 7 days of cohabitation. (I) Quantification (Repeated One-way ANOVA) of change in fluorescence signal during sniffing cagemate, an unknown male or an object after 7 days of cohabitation (n=6 voles). All error bars = SEM. Supplementary Table 1 for detailed statistics.

**Figure S11.**
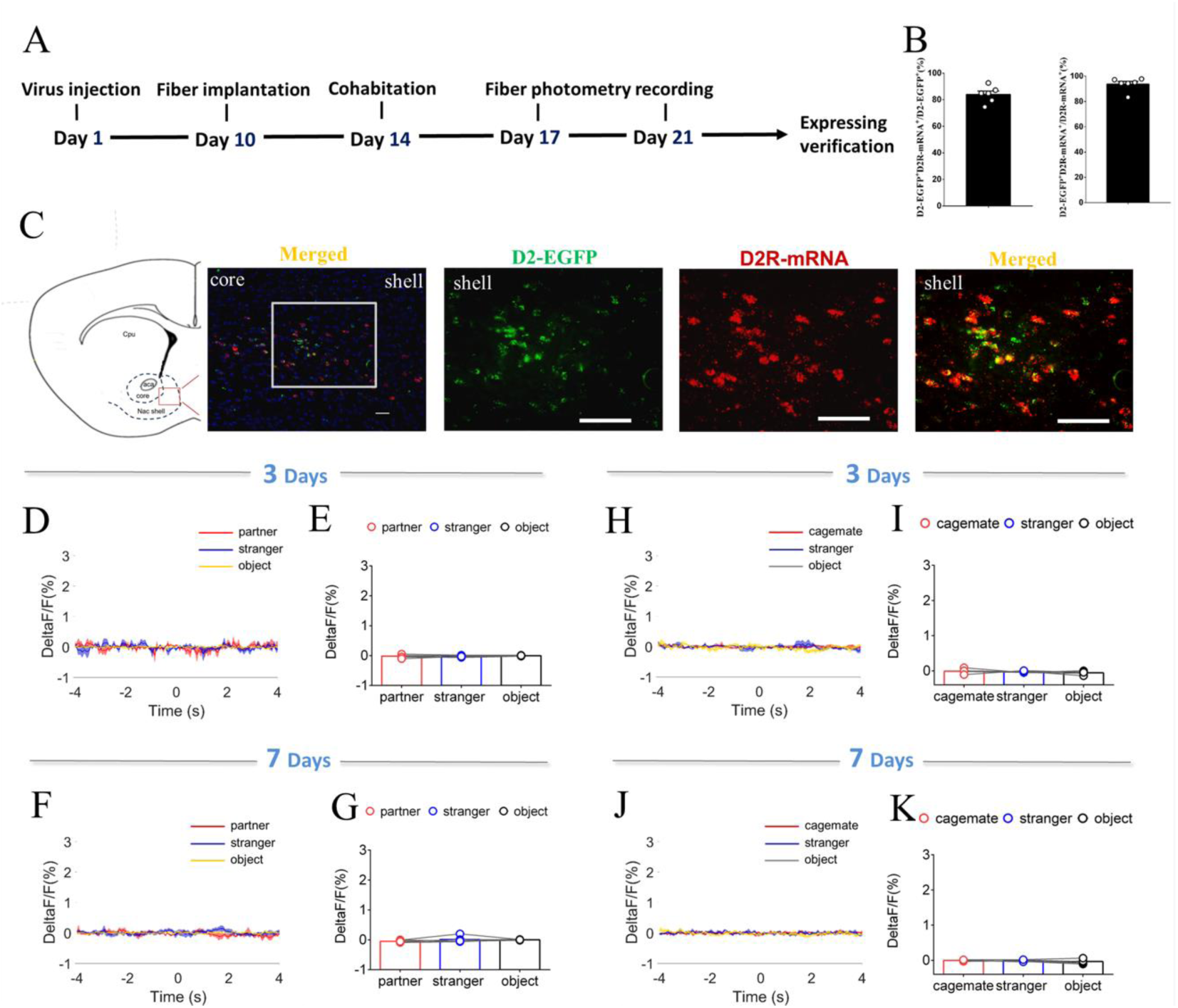
NAc shell D2 EGFP MSNs fluorescence signals after cohabitation. (A)Timeline of experiments. (B) Statistical chart showed that EGFP was relatively restricted to D2R-mRNA positive neurons (n=6 voles). (C) Overlap of EGFP (green), D2R-mRNA (red), and DAPI (blue) in the NAc shell. Scale bar: 100 μm. (D, F) Mean fluorescence signals changes of calcium response during sniffing partner (red line), an unknown female (blue line) or an object (yellow line) after cohabitation for 3 days (D) and 7 days (F). (E, G) Quantification (Repeated One-way ANOVA) of change in calcium signals during sniffing partner, an unknown female or an object after cohabitation for 3 days (E) (n = 5 voles) and 7 days (G) (n = 5 voles). (H, J) Mean fluorescence signal changes of calcium response during sniffing cagemate (red line), an unknown male (blue line) or an object (yellow line) after cohabitation for 3 days (H) and 7 days (J). (I, K) Quantification (Repeated One-way ANOVA) of change in calcium signals during sniffing cagemate, an unknown male or an object after cohabitation for 3 days (I) (n = 5 voles) and 7 days (K) (n = 5 voles). All error bars = SEM. Supplementary Table 1 for detailed statistics.

**Figure S12.**
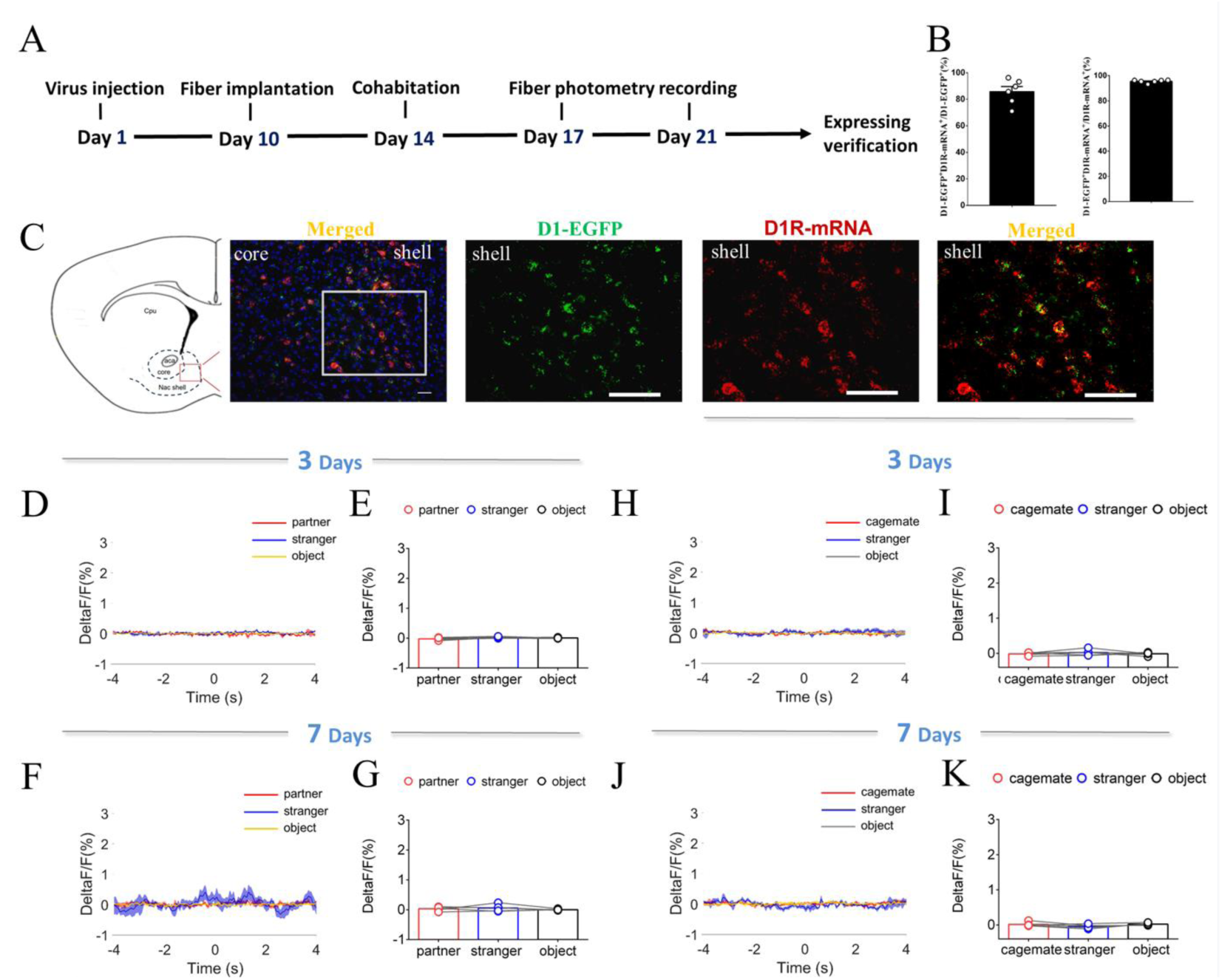
NAc shell D1 EGFP MSNs fluorescence signal with injection of control virus after cohabitation. (A)Timeline of experiments. (B) Statistical chart showed that EGFP was relatively restricted to D1R-mRNA positive neurons (n=6 voles). (C) Overlap of EGFP (green), D1R-mRNA (red), and DAPI (blue) in the NAc shell. Scale bar: 100 μm. (D, F) Mean fluorescence signal changes of calcium response during sniffing partner (red line), an unknown female (blue line) or an object (yellow line) after cohabitation for 3 days (D) and 7 days (F). (E, G) Quantification (Repeated One-way ANOVA) of change in calcium signals during sniffing partner, an unknown female or an object after cohabitation for 3 days (E) (n = 5 voles) and 7 days (G) (n = 5 voles). (H, J) Mean fluorescence signal changes of calcium response during sniffing cagemate (red line), an unknown male (blue line) or object (yellow line) after cohabitation for 3 days (H) and 7 days (J). (I, K) Quantification (Repeated One-way ANOVA) of change in calcium signals during sniffing cagemate, an unknown male or an object after cohabitation for 3 days (I) (n = 5 voles) and 7 days (K) (n = 5 voles). All error bars = SEM. Supplementary Table 1 for detailed statistics.

**Figure S13.**
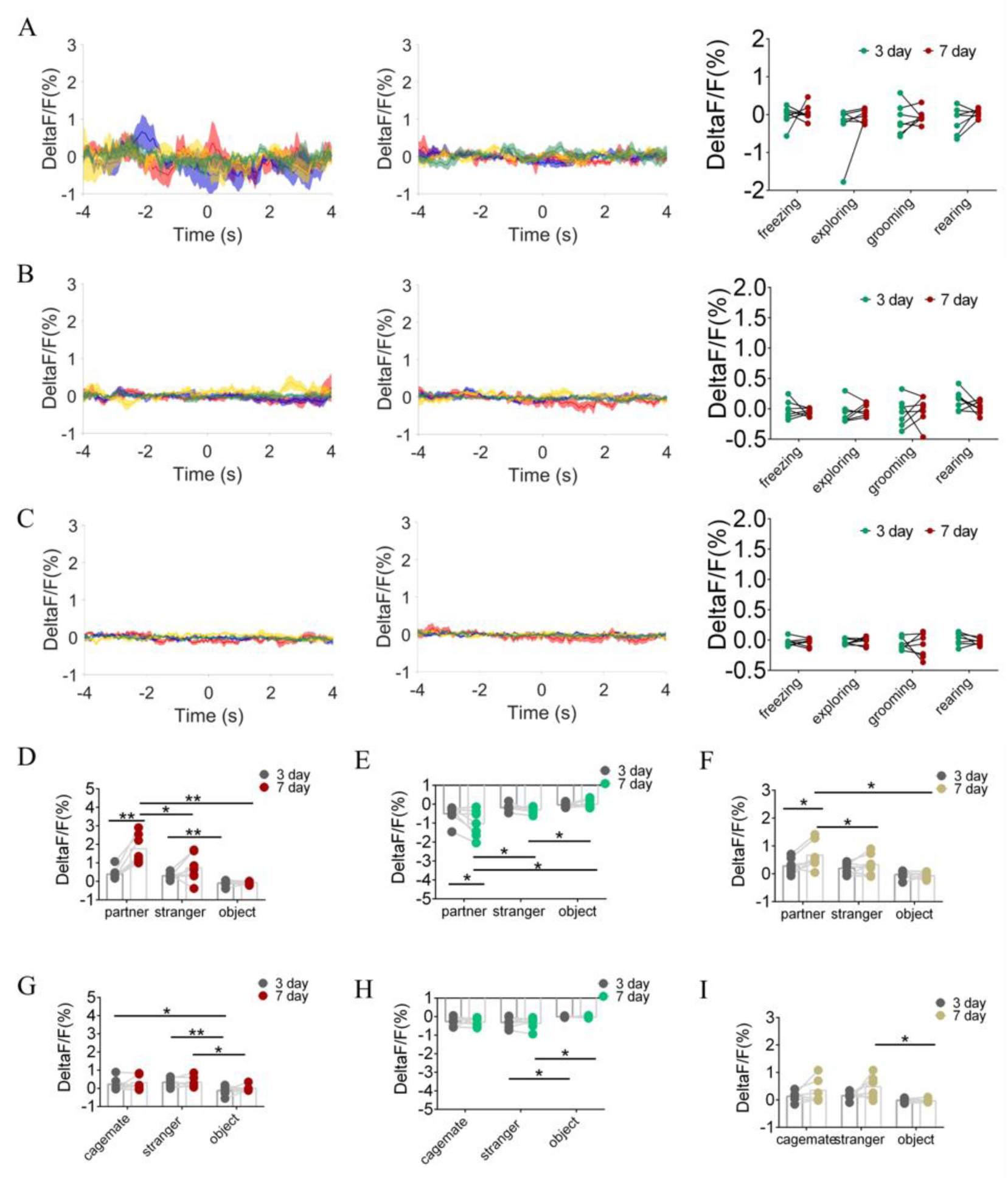
Quantification of fluorescence change during non-social behavioral bout and sniffing voles after cohabitation. (A) DA sensor: Mean fluorescence signals changes of DA concentration during freezing (red line), exploring (blue line), grooming (yellow line) and rearing (green line) after 3 days (left) and 7 days (middle) of cohabitation. Right: Quantification (Two-Way Repeated Measures ANOVA) of change during non-social behavioral bout after cohabitation with partner for 3 and 7 days (n= 7 voles). (B) D2 MSNs: Mean fluorescence signals changes of D2-MSNs during freezing (red line), exploring (blue line), grooming (yellow line) and rearing (green line) after 3 days (left) and 7 days (middle) of cohabitation. Right: Quantification (Two-Way Repeated Measures ANOVA) of change during non-social behavioral bout after cohabitation with partner for 3 and 7 days (n= 7 voles). (C) D1 MSNs: Mean fluorescence signals changes of D1-MSNs during freezing (red line), exploring (blue line), grooming (yellow line) and rearing (green line) after 3 days (left) and 7 days (middle) of cohabitation. Right: Quantification (Two-Way Repeated Measures ANOVA) of change during non-social behavioral bout after cohabitation with partner for 3 and 7 days (n= 7 voles). (D-F) Quantification (Two-Way Repeated Measures ANOVA) change sniffing partner, an unknown female or an object after cohabitation for 3 and 7 days (n= 7 voles). (D: DA sensor; E: D2 MSNs; F: D1 MSNs) (G-I) Quantification (Two-Way Repeated Measures ANOVA) of change sniffing cagemate, male stranger or object after cohabitation for 3 and 7 days (n= 6 voles). (G: DA sensor; H: D2 MSNs; I: D1 MSNs) Error bars = SEM. * represent *p* < 0.05, ** represent *p* < 0.01. See Supplementary Table 1 for detailed statistics.

**Figure S14.**
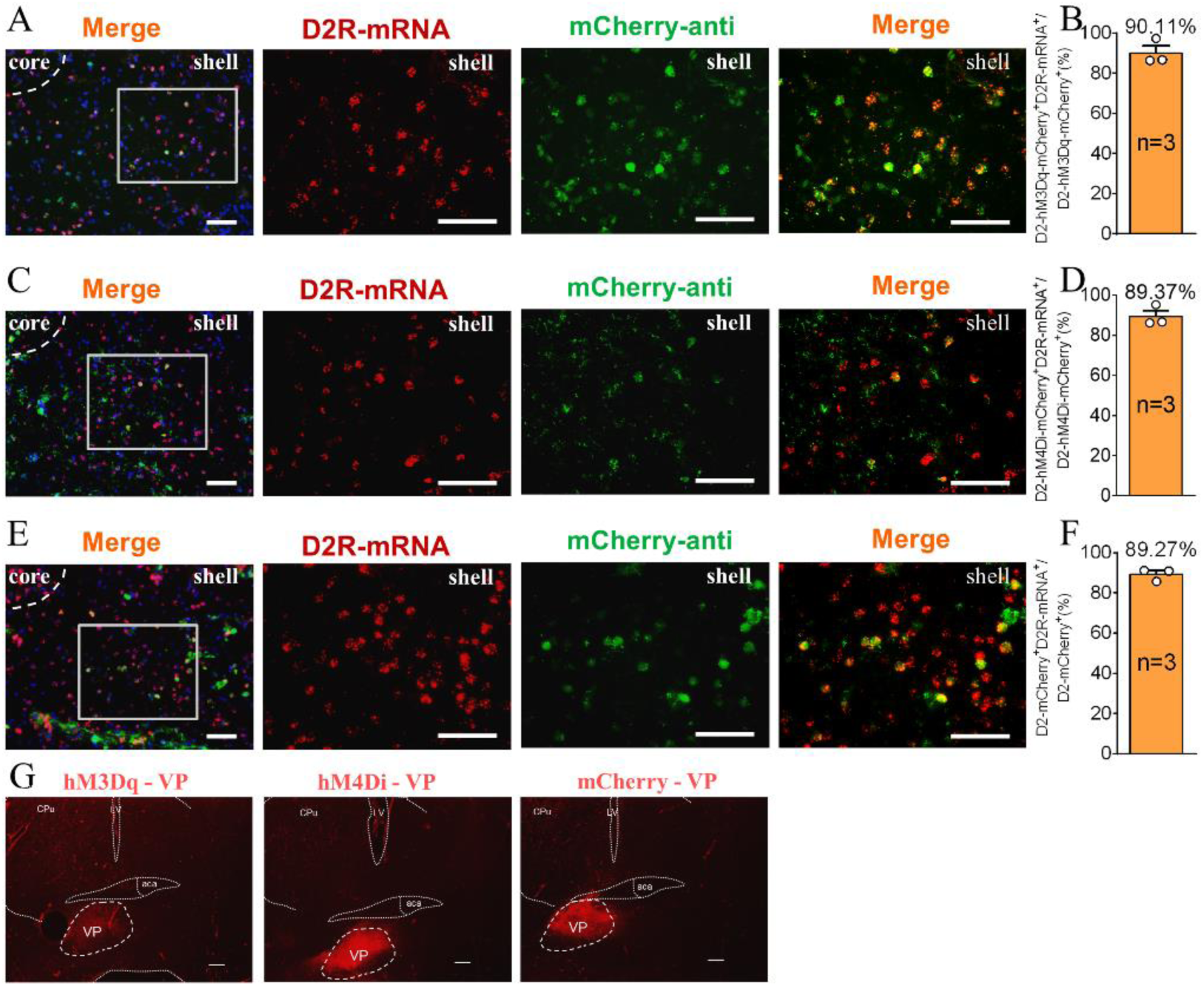
The supplement Immunohistochemistry image of the D2 MSNs chemogenetics test. (A, C, E) Immunohistochemistry image showing co-localization of hM3Dq-mCherry-anti (A), hM4Di-mCherry-anti (C) and mCherry-anti (E) expression (green), D2R-mRNA (red), and DAPI (blue) in the NAc shell. Scale bar: 100 μm. (B, D, F) Statistical chart showed that hM3Dq (B), hM4Di (D) and mCherry (F) was relatively restricted to D2 positive cells (hM3Dq n = 3 voles, hM4Di n = 3 voles, and mCherry n = 3 voles). (G) Immunohistochemistry image showing viral injection in the VP of hM3Dq (left), hM4Di (middle) and mCherry (right) group. Scale bar: 200 μm.

**Figure S15.**
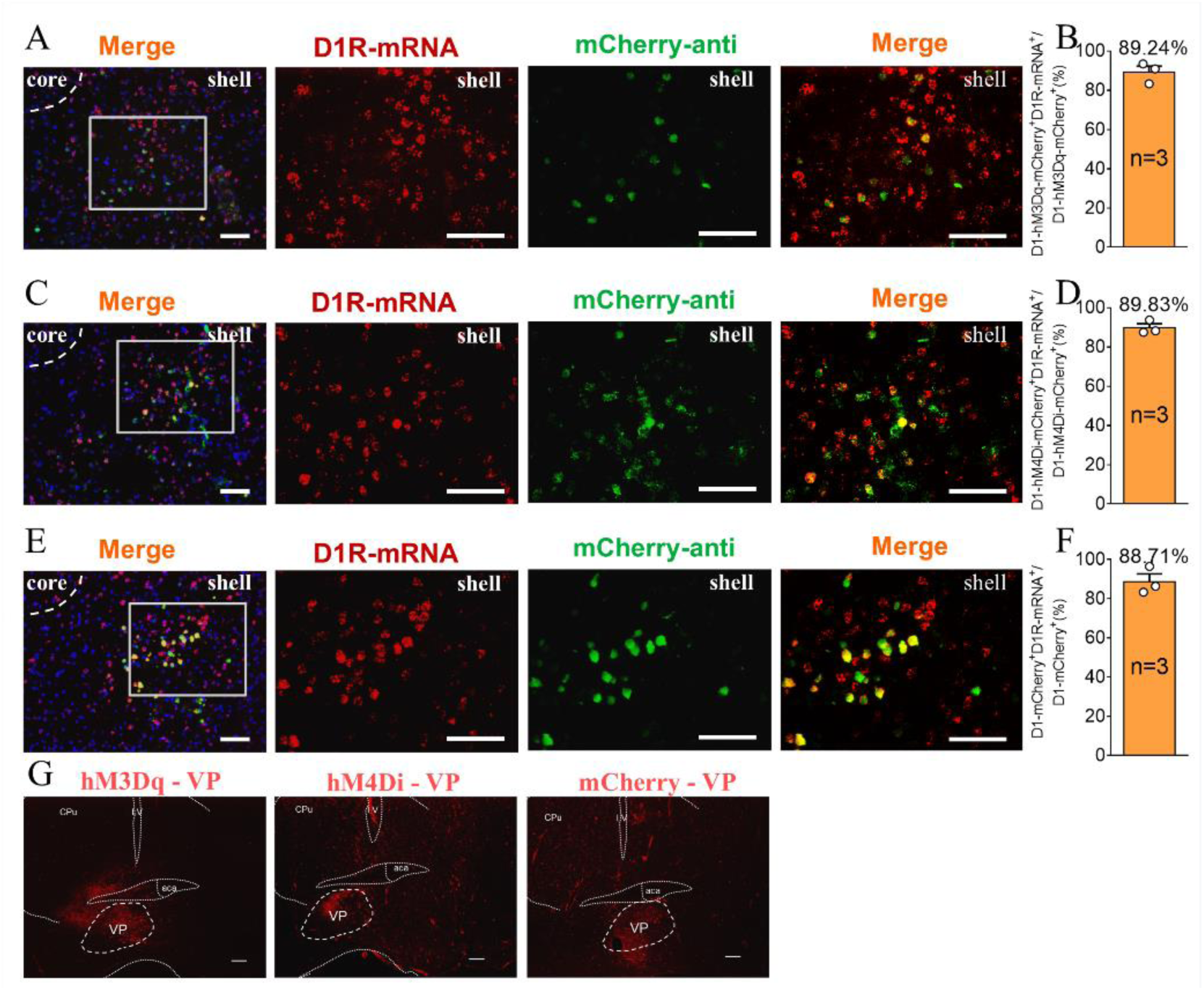
The supplement Immunohistochemistry image of the D1 MSNs chemogenetics test. (A, C, E) Immunohistochemistry image showing co-localization of hM3Dq-mCherry-anti (A), hM4Di-mCherry-anti (C) and mCherry-anti (E) expression (green), D1R-mRNA (red), and DAPI (blue) in the NAc shell. Scale bar: 100 μm. (B, D, F) Statistical chart showed that hM3Dq (B), hM4Di (D) and mCherry (F) was relatively restricted to D1 positive cells (hM3Dq n = 3 voles, hM4Di n = 3 voles, and mCherry n = 3 voles). (G) Immunohistochemistry image showing viral injections in the VP of hM3Dq (left), hM4Di (middle) and mCherry (right) group. Scale bar: 200 μm.

**Figure S16.**
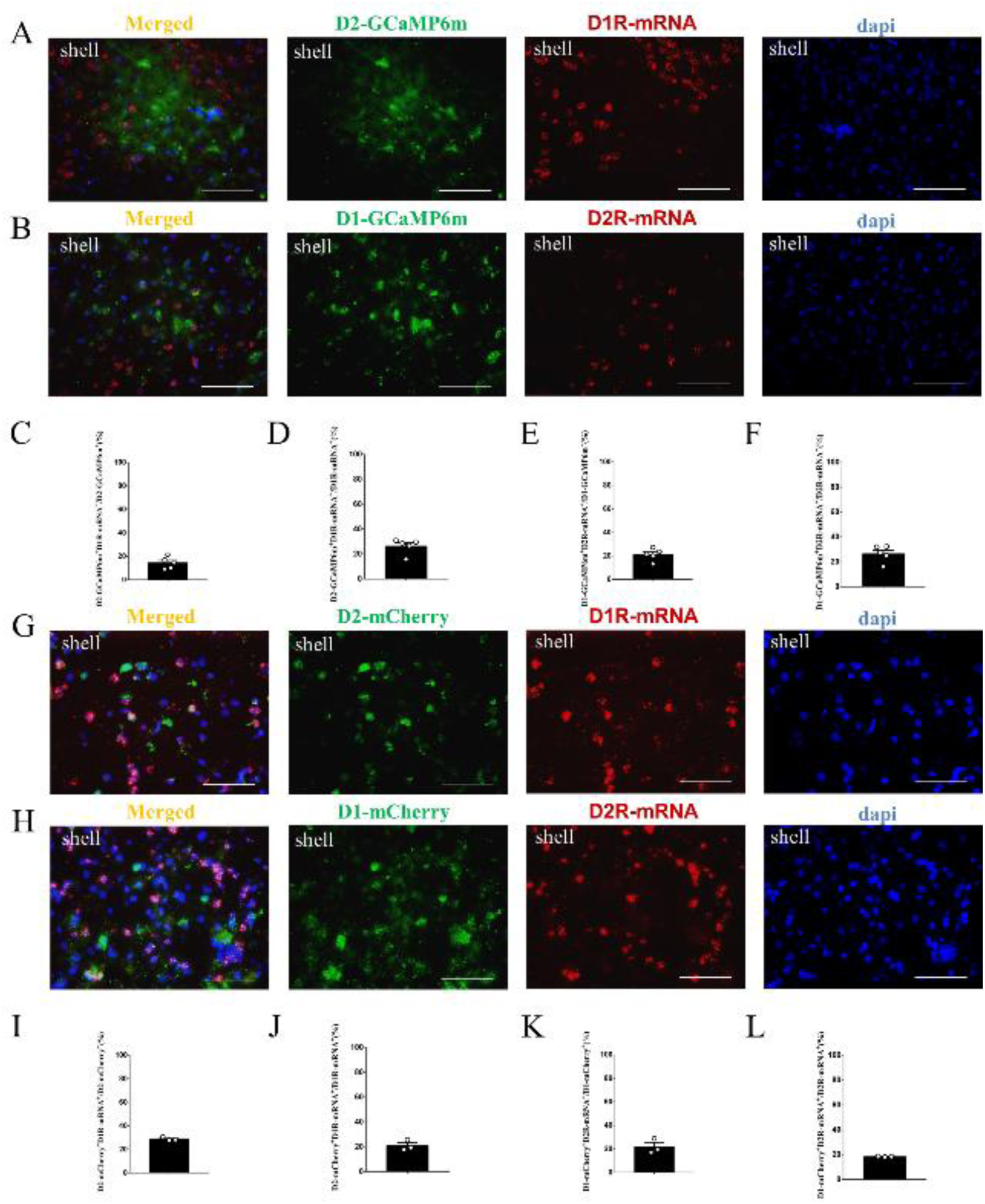
The supplement FISH image of the D1-GCaMP6m / D2-GCaMP6m MSNs in the NAc shell. (A) Overlap of D2-GCaMP6m (green), D1R-mRNA (red), and DAPI (blue) in the NAc shell. Scale bar: 100 μm. (B) Overlap of D1-GCaMP6m (green), D2R-mRNA (red), and DAPI (blue) in the NAc shell. Scale bar: 100 μm. (C-D) Statistical chart showed that D2-GCaMP6m was relatively restricted to D1R-mRNA positive neurons (n = 5 voles). (E-F) Statistical chart showed that D1-GCaMP6m was relatively restricted to D2R-mRNA positive neurons (n = 5 voles). (G) Overlap of D2-mCherry (green), D1R-mRNA (red), and DAPI (blue) in the NAc shell. Scale bar: 100 μm. (H) Overlap of D1-mCherry (green), D2R-mRNA (red), and DAPI (blue) in the NAc shell. Scale bar: 100 μm. (I-J) Statistical chart showed that D2-mCherry was relatively restricted to D1R-mRNA positive neurons (n = 3 voles). (K-L) Statistical chart showed that D1-mCherry was relatively restricted to D2R-mRNA positive neurons (n = 3 voles).

**Figure S17.**
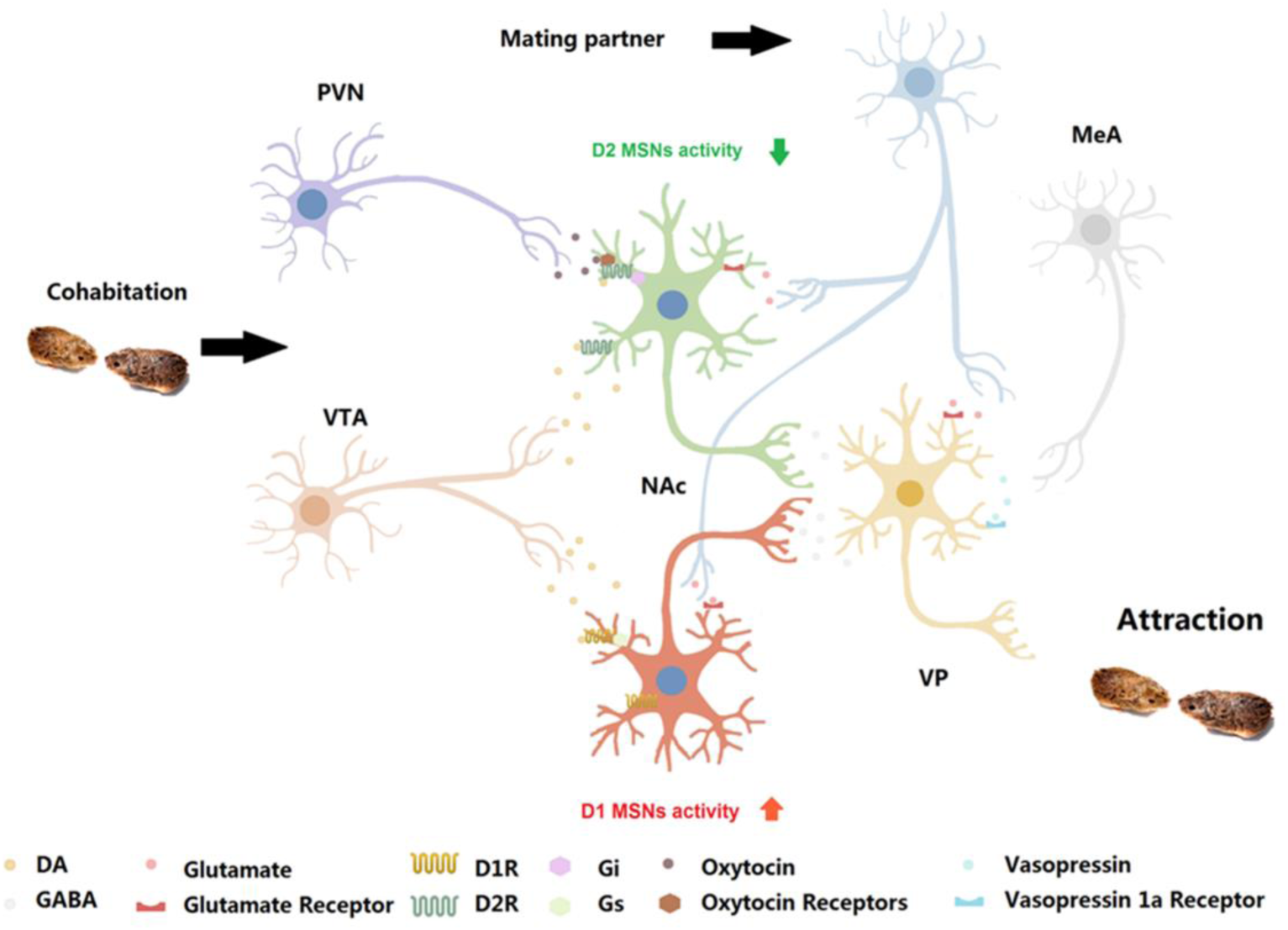
Neurobiological mechanism and circuit mechanism underlying the formation of a pair bond. Possible neural model within NAc, where the co-action of DA on D2R and oxytocin on oxytocin receptors depress the output of MSNs to VP; then, the VP mediates the attraction of partners during the formation of the pair bond. BLA: basolateral amygdala; MeA: medial amygdala; VTA: ventral tegmental area; PVN: paraventricular nucleus.

